# Finding structure in multi-armed bandits

**DOI:** 10.1101/432534

**Authors:** Eric Schulz, Nicholas T. Franklin, Samuel J. Gershman

## Abstract

How do humans search for rewards? This question is commonly studied using multi-armed bandit tasks, which require participants to trade off exploration and exploitation. Standard multi-armed bandits assume that each option has an independent reward distribution. However, learning about options independently is unrealistic, since in the real world options often share an underlying structure. We study a class of *structured* bandit tasks, which we use to probe how generalization guides exploration. In a structured multi-armed bandit, options have a correlation structure dictated by a latent function. We focus on bandits in which rewards are linear functions of an option’s spatial position. Across 5 experiments, we find evidence that participants utilize functional structure to guide their exploration, and also exhibit a learning-to-learn effect across rounds, becoming progressively faster at identifying the latent function. Our experiments rule out several heuristic explanations and show that the same findings obtain with non-linear functions. Comparing several models of learning and decision making, we find that the best model of human behavior in our tasks combines three computational mechanisms: (1) function learning, (2) clustering of reward distributions across rounds, and (3) uncertainty-guided exploration. Our results suggest that human reinforcement learning can utilize latent structure in sophisticated ways to improve efficiency.

## Introduction

Imagine walking into a nearby supermarket to buy groceries for tonight’s dinner. How do you decide what is good to eat? If you had enough time, you could try everything in the store multiple times to get a good sense of what you liked and then repeatedly buy what you liked the most. However, this level of exhaustive exploration is unrealistic. In real-world scenarios, there are far too many choices for anyone to try them all. Alternatively, you could explore in proportion to your uncertainty about the value of each item (Auer, Cesa-Bianchi, & Fischer, 2002; Gershman, 2018; Schulz & Gershman, 2019), choosing foods you know little about until you have accumulated enough knowledge to reliably select the ones you like.

Another strategy for intelligent exploration is based on generalization (Boyan & Moore, 1995; Shepard, 1987; Tenenbaum & Griffiths, 2001). For example, if you know that a grocery store has good apples, it probably also has good pears, but knowing that it has good apples might not tell you much about its fish. This paper studies how people combine generalization and uncertainty to guide exploration.

In experimental settings, the exploration-exploitation trade-off is commonly studied using *multi-armed bandit tasks* (Bechara, Damasio, Tranel, & Damasio, 2005; J. D. Cohen, McClure, & Angela, 2007; Mehlhorn et al., 2015). The term multi-armed bandit is based on a metaphor for a row of slot machines in a casino, where each slot machine has an independent payoff distribution. Solutions to these problems propose different policies for how to learn about which arms are better to play (exploration), while also playing known high-value arms to maximize reward (exploitation). In a typical multi-armed bandit task, participants sample options sequentially to maximize reward (Speekenbrink & Konstantinidis, 2015; Steingroever, Wetzels, Horstmann, Neumann, & Wagenmakers, 2013). To do so, they start out with no information about the underlying rewards and can only observe outcomes by iteratively choosing options (but see Navarro, Newell, & Schulze, 2016, for a different setup). Optimal solutions to multi-armed bandit tasks are generally intractable (Gittins, 1979; Whittle, 1980) and thus performing well requires heuristic solutions (e.g., Auer et al., 2002; Chapelle & Li, 2011).

In almost all of these experimental and machine learning settings, the available options are assumed to have independent reward distributions, and therefore computational models of human behavior in such tasks inherit this assumption (see Steingroever et al., 2013, for a detailed comparison of such models). This assumption, while useful experimentally, does not correspond well with actual human learning. Many real-world tasks are governed by a latent structure that induces correlations between options (Gershman & Niv, 2010). To return to the supermarket example, the values of different food items are correlated because food categories are organized spatially and conceptually. Understanding this structure enables an agent to explore more efficiently by generalizing across foods. Structure is the *sine qua non* of generalization (Gershman, Malmaud, & Tenenbaum, 2017).

This point has already received some attention in the bandit literature. Previous work on structure learning in bandit tasks has found evidence for two types of structure learning: learning a shared structure across the arms of a bandit (Acuna & Schrater, 2010; Gershman & Niv, 2015; Wu, Schulz, Speekenbrink, Nelson, & Meder, 2018) or learning the latent structure underlying a set of stimulus-response mappings (Badre, Kayser, & D’Esposito, 2010; Collins & Frank, 2013). Furthermore, human participants can sometimes assume a more complex structure even if it is not present (Zhang & Yu, 2013), and even when working memory load makes structure more costly to infer (Collins, 2017; Collins & Frank, 2013). If there actually is explicit structure in a bandit task (i.e., participants are told about the presence of an underlying structure), they are able to effectively exploit this structure to speed up learning (Acuna & Schrater, 2010; Gershman & Niv, 2015; Goldstone & Ashpole, 2004; Gureckis & Love, 2009; Wu, Schulz, Speekenbrink, et al., 2018).

We go beyond these past studies to develop a general theory of structured exploration in which exploration and structure learning are interlocking components of human learning. We propose that humans use structure for directed exploration and generalize over previously learned structures to guide future decisions. To test this theory experimentally, we use a paradigm called the *structured multi-armed bandit task*. A structured multi-armed bandit looks like a normal multi-armed bandit but—unknown to participants—the expected reward of an arm is related to its spatial position on the keyboard by an unknown function. Learning about this function can improve performance by licensing strong generalizations across options. Further performance improvements can be achieved by generalizing latent structure across bandits—a form of “learning-to-learn” (Andrychowicz et al., 2016; Harlow, 1949). The task further affords us the ability to ask what types of structure subjects use when learning. We use computational modeling to address this question.

In the following section, we outline the structured multi-armed bandit task before discussing a series of human behavioral experiments and accompanying computational models that investigate human structure learning and exploration across 5 variants of our task.

### Structured multi-armed bandits

Let us formally define the structured multi-armed bandit task. On each trial *t*, participants choose one of *J* options, *a_t_* ∈ {1, …, *J*}, and observe reward 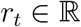, drawn from a latent function *f* corrupted by noise *ϵ_t_*:

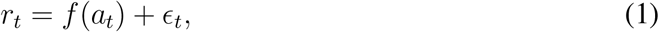

where 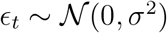. Importantly, the *J* options are spatially contiguous (see Fig. 1), such that the reward function can be understood as mapping a one-dimensional spatial position to reward.

**Figure 1.**
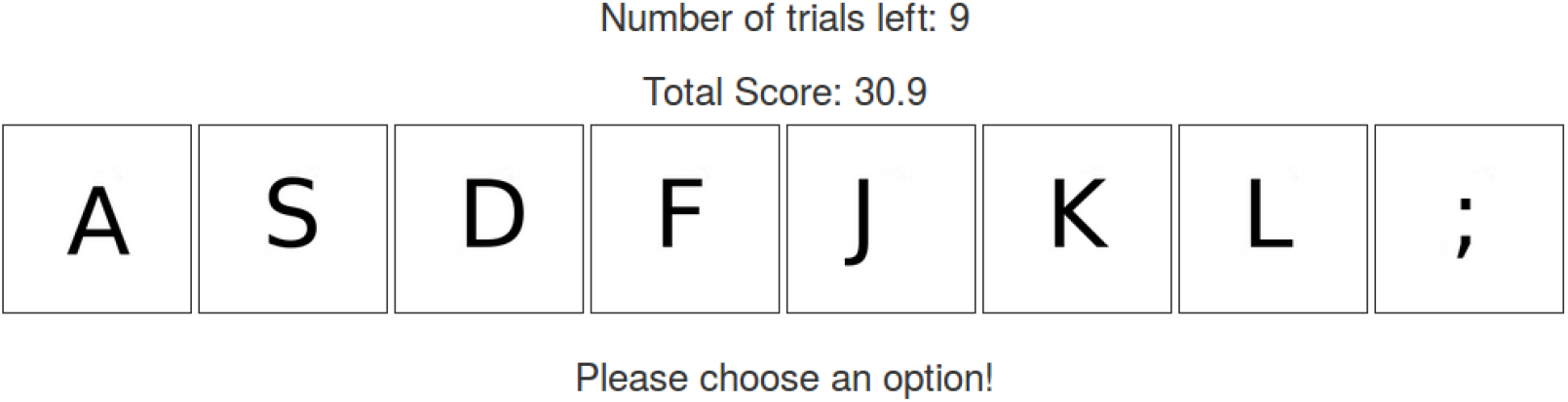
Screenshot of the structured bandit task. There are 8 options (arms) in total, each marked by a letter/symbol. When participants pressed the corresponding key on their keyboard, the matching arm was sampled and the output for that option appeared underneath all arms for 2 seconds. The number of trials left as well as the total score in that round so far were displayed above all options.

In standard bandit tasks, the reward function *f* is assumed to be tabular (i.e,. the value of each arm in each trial is independent of all others). This simplifies the learning and exploration problem, but prevents generalization between options. Here, we allow for such generalization by modeling the reward function with a Gaussian Process (GP; Rasmussen & Williams, 2006; Schulz, Speekenbrink, & Krause, 2018). A GP is a distribution over functions defined by a mean function *μ* and a covariance function (or *kernel*) *k*:

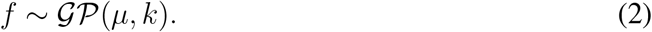

The mean function controls the function’s expected value, and the covariance function controls its expected smoothness:

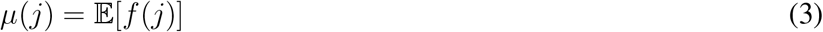

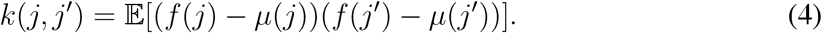

Following convention (Rasmussen & Williams, 2006), we will assume that the mean function is 0 everywhere, *μ*(*j*) = 0. Our focus will be on the choice of kernel, which can be parametrized to encode specific assumptions about the correlation structure across options (Duvenaud, 2014; Schulz, Tenenbaum, Duvenaud, Speekenbrink, & Gershman, 2017). For example, a periodic kernel assumes that functions oscillate over space, while a linear kernel assumes that functions change linearly. We will discuss specific choices of kernels later when we develop models of our experimental data. Note, however, that the focus of the current work is on whether and how participants use latent structure when making decisions to gather rewards. Which exact kernel describes participants’ generalization patterns best has been the main question of our previous work (Schulz, Tenenbaum, et al., 2017; Schulz, Tenenbaum, Reshef, Speekenbrink, & Gershman, 2015).

Our experiments are designed to test the following hypotheses about learning in structured multi-armed bandits:

1. Participants will perform better across trials on rounds where there is a learnable latent structure, compared to rounds where each option’s reward is independent of its position, without being prompted that there is an underlying structure.
2. Participants will learn more effectively in later than in earlier rounds, as they learn about the higher-order distribution over latent functions (i.e., learning-to-learn).

If these hypotheses are correct, we expect that a model that best captures human behavior in our task will incorporate both a mechanism for learning within a given round (generalization over options) as well as a mechanism for learning across rounds (generalization over functions).

### Overview of experiments and models

Our experiments are designed to assess if and how participants benefit from latent functional structure in a bandit task. We test if they learn about the underlying structure over options by applying function learning. We assess if they learn about the higher-order distribution of functions over rounds by a learning-to-learn mechanism. We also rule out several alternative explanations of our results. An overview of all experiments and their results can be found in Table 1.

**Table 1.**
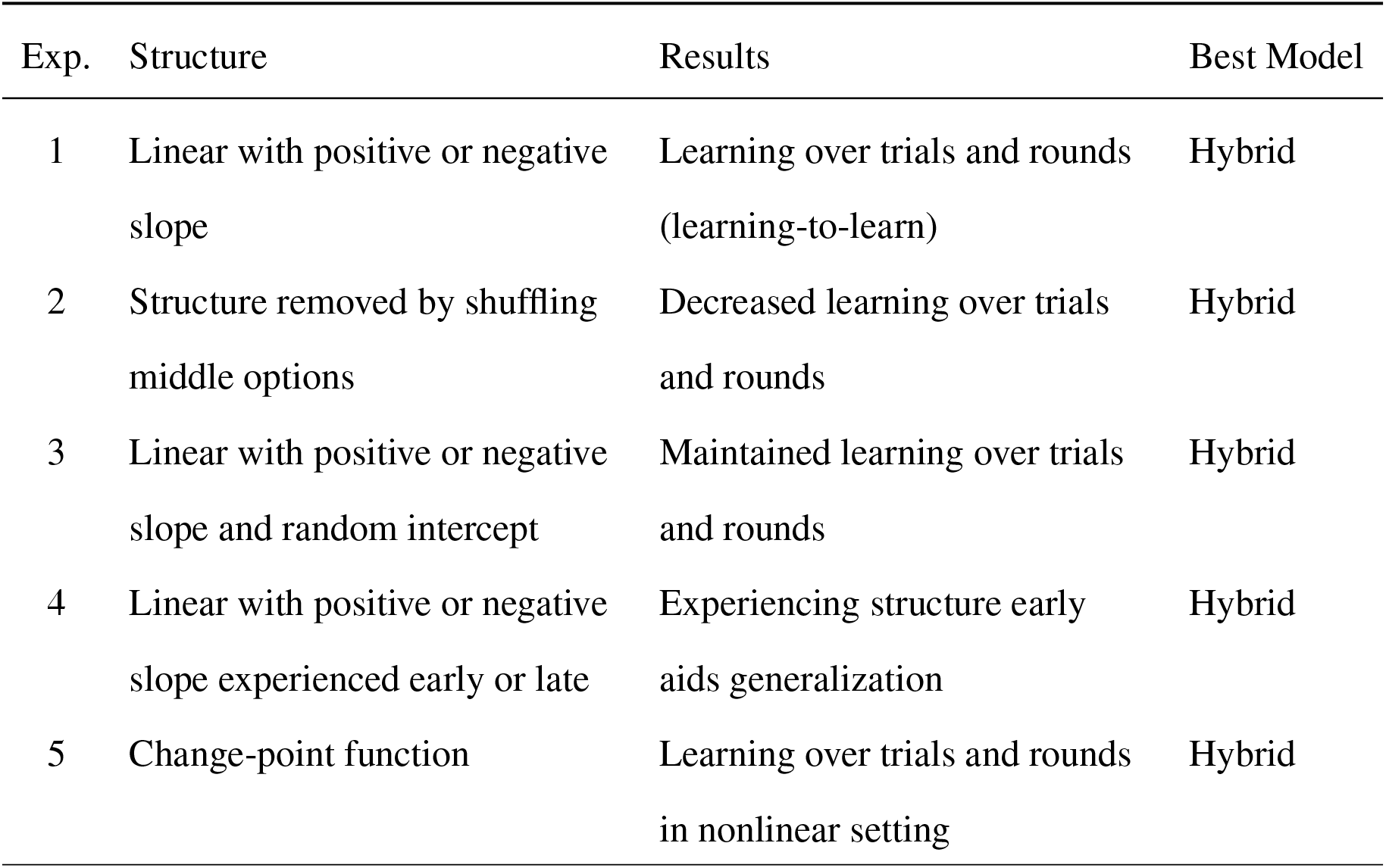
Overview of experiments and results. The Hybrid model combines a Gaussian Process model of function learning with a Clustering model which learns how options differ over rounds.

| Exp. | Structure | Results | Best Model |
| --- | --- | --- | --- |
| 1 | Linear with positive or negative slope | Learning over trials and rounds (learning-to-learn) | Hybrid |
| 2 | Structure removed by shuffling middle options | Decreased learning over trials and rounds | Hybrid |
| 3 | Linear with positive or negative slope and random intercept | Maintained learning over trials and rounds | Hybrid |
| 4 | Linear with positive or negative slope experienced early or late | Experiencing structure early aids generalization | Hybrid |
| 5 | Change-point function | Learning over trials and rounds in nonlinear setting | Hybrid |

In Experiment 1, we interleave standard (uncorrelated) multi-armed bandit rounds with rounds in which an option’s reward was mapped to its position by an underlying linear function with either positive or negative slope. Our results show that participants are able to learn this structure and show improvements in detecting this structure over rounds (i.e., learning-to-learn). In Experiments 2 and 3, we control for potential heuristic strategies that can produce similar learning profiles to those found in Experiment 1.

In Experiment 2, we examine whether participants rely on only the leftmost and rightmost options to solve the task or are sensitive to the structure across all options. By randomly shuffling the expected rewards of all options except the leftmost and rightmost options, we remove the functional structure within rounds while maintaining similar levels of uncertainty. We find that participants perform worse overall, suggesting they use the full functional structure to solve the task.

In Experiment 3, we randomly change the range of possible rewards, making it difficult to learn the global optimum, and show that participants can still generalize about the underlying function.

In Experiment 4, we examine order effects and probe how exposure to structured rounds during either earlier or later stages of the task influence performance.

Finally, in Experiment 5 we push the limits of participants’ generalization ability by assessing their performance in a structured bandit task with a nonlinear change-point function, which increases linearly up to a point and then decreases again. Participants are once again able to generalize successfully.

Across all five experiments, we use computational modeling to test multiple, alternative hypotheses of structure learning, and ask what form of structure learning best describes participants’ behavior. We find that the same computational model best fits behavior in each of the experiments. This model combines a mechanism of function learning to learn the structure within each round, a mechanism of option clustering to improve initial reward expectations over rounds (learning-to-learn), and a decision strategy that solves the exploration-exploitation dilemma by trading off between an option’s expected reward and uncertainty. We use generative simulations to show that the winning model can reproduce human-like learning, and that our model comparison results are robust and recoverable. These modeling results suggest that simpler accounts of learning that rely on a single mechanism for learning structure or exploration are insufficient to explain learning in these task, and suggest that structure learning and exploration are naturally linked.

## Experiment 1: Linear structure

Our first investigation uses one of the simplest latent structures: linear functions. Intuitively, a participant who observes that outcomes are increasing as a function of choosing more left or right options will be able to make the inference that she should jump right to the leftmost or rightmost option rather than continuing to sample intermediate options. Moreover, repeatedly encountering this structure should teach the participant that the most highly rewarding options tend to be on the extremes, thus biasing sampling for choices at the start of each round.

### Participants

One hundred and sixteen participants (50 females, mean age=32.86, SD=8.25) were recruited from Amazon Mechanical Turk and were paid $2.00 plus a performance-dependent bonus of up to $1.50 for their participation. Participants in this and all of the following experiments were required to have a historical acceptance rate on Mechanical Turk of greater than 95% and a history of at least 1000 successfully completed tasks. The experiment took 28.64 minutes to complete on average.

### Design

Participants played 30 rounds of a multi-armed bandit task, where each round contained 10 sequential trials in which participants were instructed to gain as many points as possible by sampling options with high rewards. The experiment used a within-subjects design, where 10 of the rounds contained an underlying linear function with a positive slope (*linear-positive*), 10 a linear function with a negative slope (*linear-negative*), and 10 contained no structure at all (i.e., shuffled linear functions; *random*). A screenshot of the experiment is shown in Figure 1.

### Procedure

Participants were instructed to place their hands as shown in Figure 2 and could sample from each option by pressing the corresponding key on their keyboard. We excluded participants who did not have a standard “QWERTY” keyboard from this and all of the following experiments.

**Figure 2.**
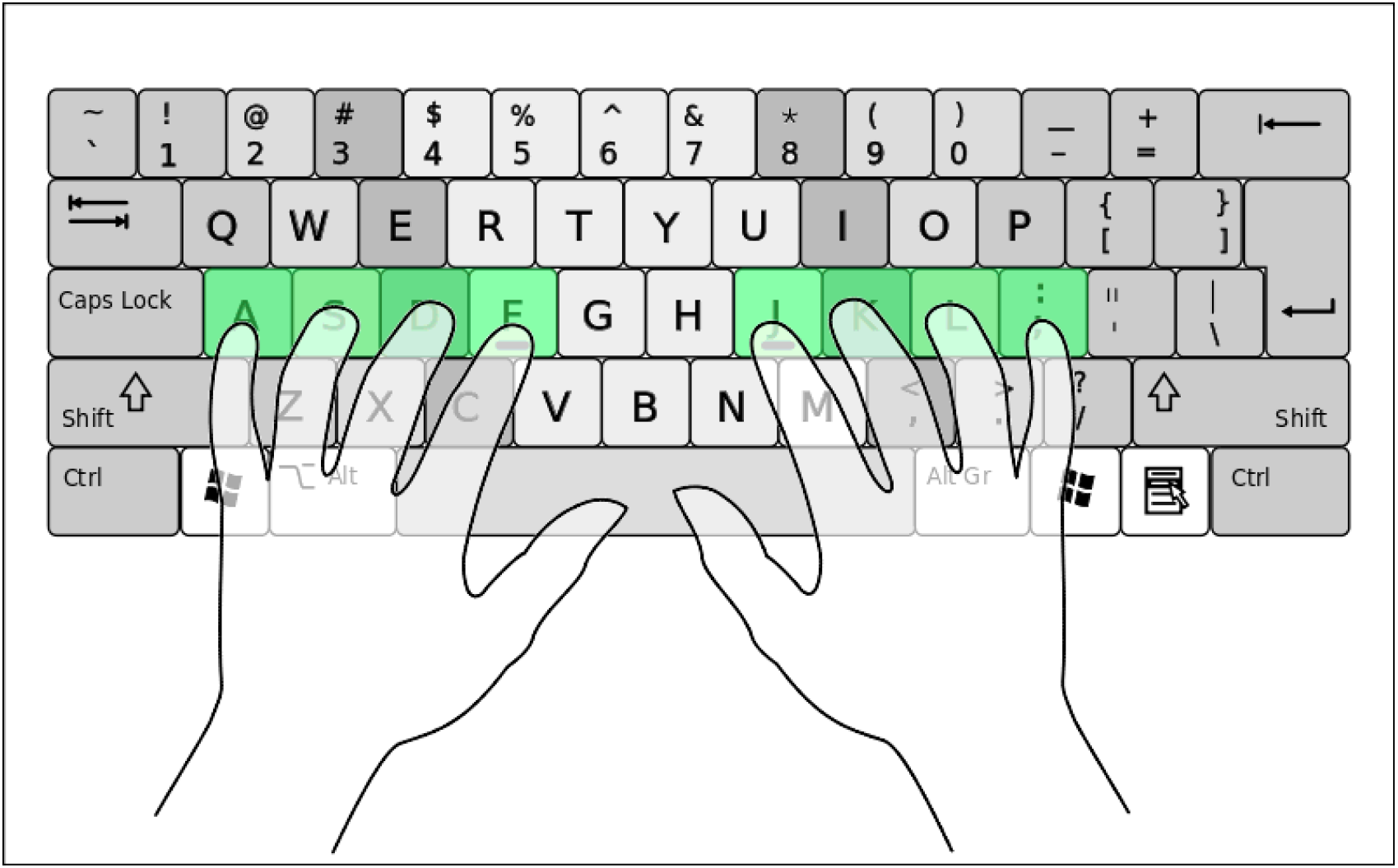
Finger position participants were asked to adhere to during all experiments. Green keys correspond to the options labeled by the same symbol. Pressing the corresponding key would sample the option with that label. Pressing any other key or pressing one of the valid keys during the 2 seconds after which the last valid key was pressed would trigger a warning.

There were 30 rounds in total, each containing 10 trials, for a total number of 300 trials. Participants were told that after each round the game resets and that each option could then produce different rewards. After they had passed four comprehension check questions, participants were allowed to start the game. We did not exclude any participant who had failed the comprehension check questions, but sent them back to the first page of the instructions, repeating this loop until they had answered all questions correctly.

Ten of the thirty rounds were randomly selected and their underlying structure was designed to follow a linear structure with a positive slope (i.e., sampling arms further towards the right would lead to higher rewards). We refer to those rounds as *linear-positive* rounds. Ten of the remaining twenty rounds were randomly selected and their underlying structure was designed to follow a linear structure with a negative slope (i.e., sampling arms towards the left would lead to higher rewards). We refer to those rounds as *linear-negative* rounds. The remaining 10 rounds did not follow any spatial structure; the reward function was generated by taking the linearly structured functions (5 from the linear-positive and 5 from the linear-negative set) and randomly shuffling their reward function across positions. We refer to those rounds as *random* rounds. Thus, participants encountered 10 rounds of each condition interleaved at random.

All linear functions were sampled from a Gaussian Process parameterized by a linear kernel:

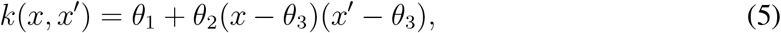

where *θ*_1_ defines the intercept and *θ*_2_ and *θ*_3_ the slope of the linear function respectively. We rescaled the minimum of each function per block to be sampled uniformly from 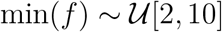 and the maximum to be 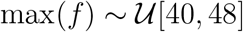 to create extremes that were not easily guessable (for example, 0 or 50). This implies linear functions with varying intercepts and positive or negative slopes.

Figure 3 shows the pool of functions from which we sampled the underlying structure for each block without replacement. On every trial, participants received a reward by sampling an arm, with the reward determined by the position of the arm:

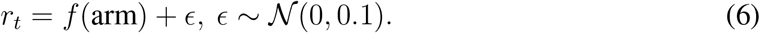

**Figure 3.**
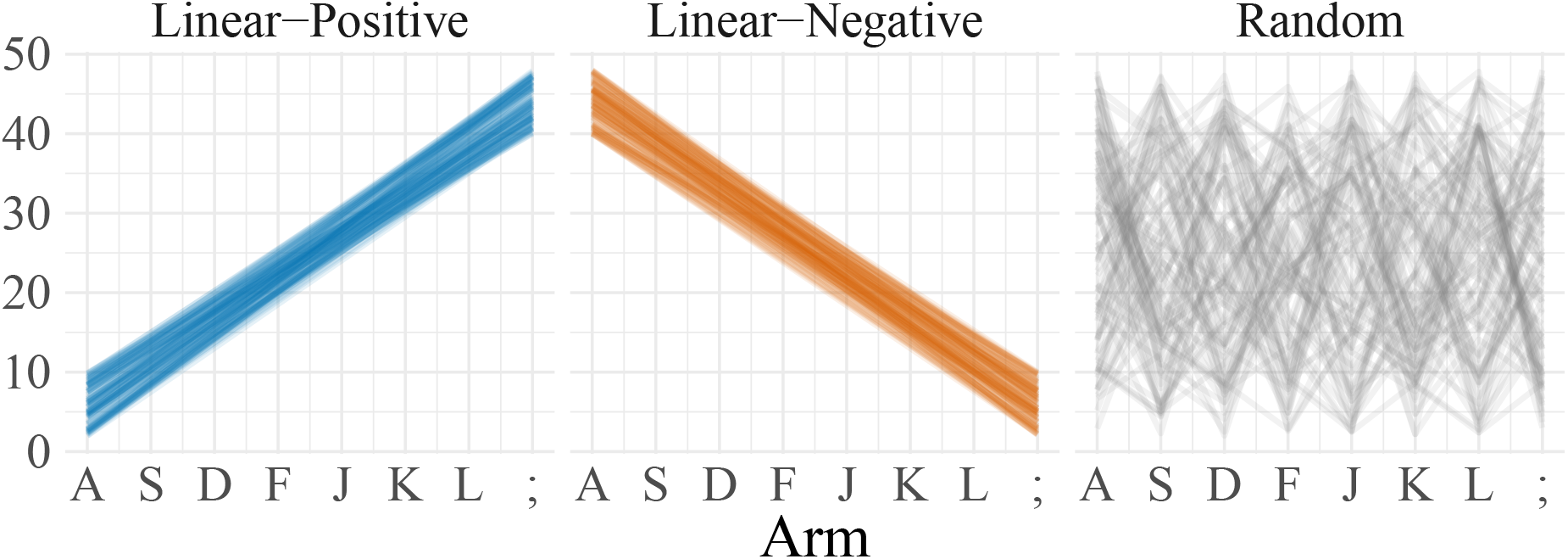
Pool of functions from which underlying structure was sampled in Experiment 1 (without replacement). We created 100 linear functions with a positive slope, 100 linear functions with a negative slope, and 100 random functions (50 of which were functions with a positive slope but shuffled, 50 of which were shuffled linear functions with a negative slope).

Participants were paid a performance bonus (on top of the $2.00 participation fee) of $0.01 for every 100 points they gained in the experiment (with points varying between 0 and 50 for every sample), leading to a maximum bonus of up to $1.50. As the maximum of $1.50 was unlikely to be achieved, we also told participants the expected range of rewards over all rounds, which—based on pilot data (see Appendix C; the pilot data with 60 participants also showed all main behavioral effects reported here)—was assumed to vary between $0.65 and $1.05.

### Results and discussion

Participants learned about and benefited from the latent structure in our task. Participants performed better in linear-positive than in random rounds (Fig. 4A; *t*(115) = 7.71, *p* < .001, *d* = 0.72; *BF* > 100) and better during linear-negative than during random rounds (*t*(115) = 11.46, *p* < .001, *d* = 1.06; *BF* > 100). Participants did not perform better in linear-positive than in linear-negative rounds (*t*(115) = 1.42, *p* = .16 *d* = 0.13; *BF* = 0.3). There were no differences in overall reaction times between the different conditions (Fig. 4B; all *p* > .05; max-*BF* = 0.2).

**Figure 4.**
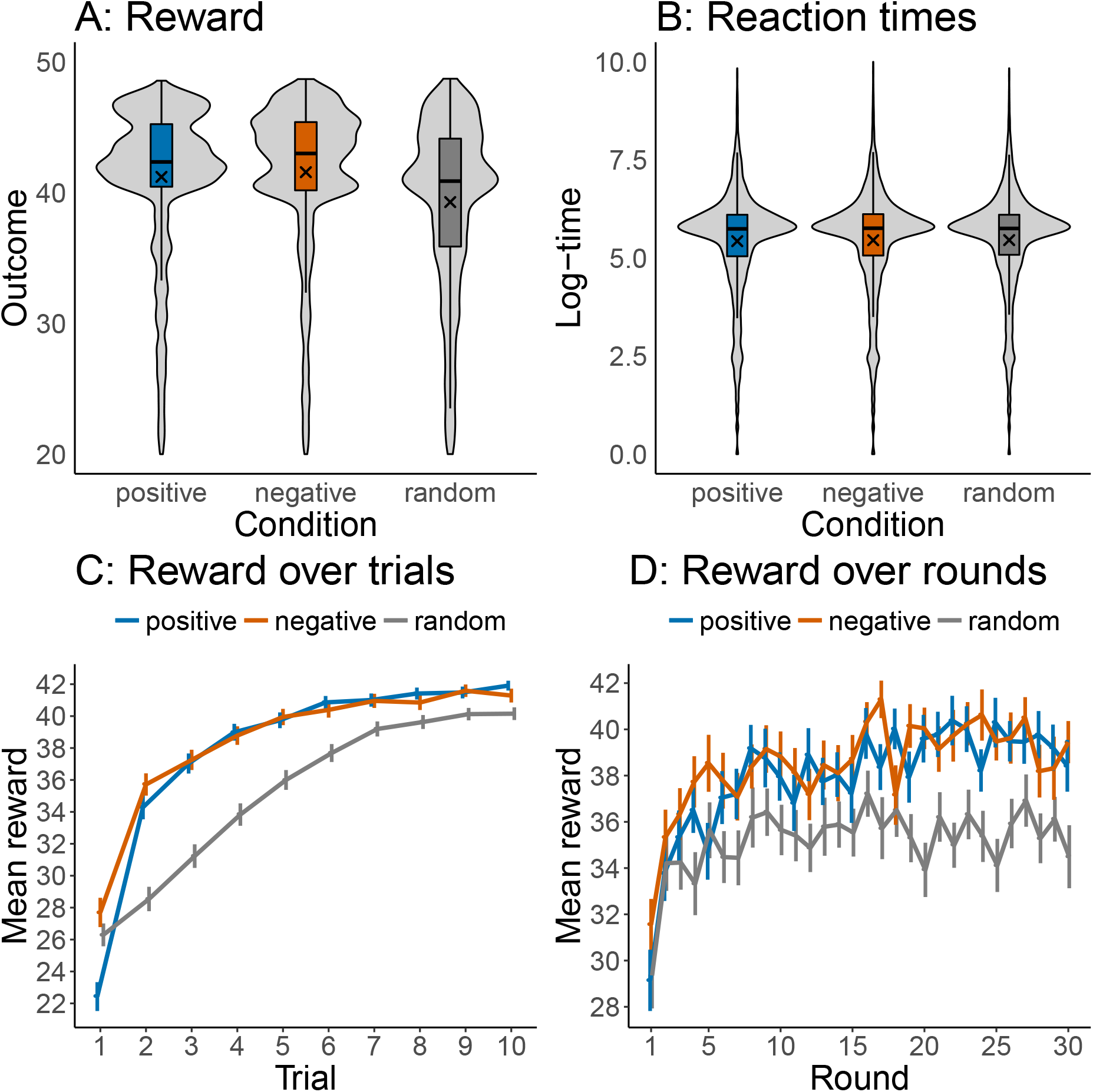
Results of Experiment 1. **A:** Box-plots of rewards by condition including violin density plots. Crosses mark the means by condition. Distributions show the rewards over all trials and rounds. **B:** Same plot but for reaction times measured in log-milliseconds. **C:** Reward over trials aggregated over all rounds. **D:** Reward over rounds aggregated over all trials. Error bars represent the 95% confidence interval of the mean.

Next, we analyzed participants’ learning rates. Participants improved over trials, with a mean correlation^1^ between trials and rewards of *ρ* = 0.29, *t*(115) = 34.6, *p* < .001, *d* = 3.21, *BF* > 100 (see Fig. 4C). The correlation between trials and rewards was higher for linear-positive than for random rounds (*t*(115) = 4.48, *p* < .001, *d* = 0.42; *BF* > 100) and higher for linear-negative than for random rounds (*t*(115) = 10.58, *p* < .001, *d* = 0.95; *BF* > 100). It was also higher for linear-positive than for linear-negative rounds *t*(115) = 4.95, *p* < .001, *d* = 0.46; *BF* > 100), although this difference was mostly driven by participants frequently sampling the leftmost option first and thus disappeared when removing the first trial from the correlation analysis (*t*(115) = 1.92, *p* = .06, *d* = 0.18; *BF* = 0.6).

Participants improved over rounds. The mean correlation between rounds and rewards was *ρ* = 0.10 (*t*(115) = 6.78, *p* < .001, *d* = 0.63, *BF* > 100; see Fig. 4D). The correlation between rounds and rewards was higher for the linear-positive than for the random condition (*t*(115) = 4.20, *p* < .001, *d* = 0.39; *BF* > 100) and higher for the linear-negative than for the random condition (*t*(115) = 3.76, *p* < .001, *d* = 0.34; *BF* = 71). There was no such difference between the linear-positive and the linear-negative condition (*t*(115) = 0.16, *p* = .86, *d* = 0.02; *BF* = 0.1). This indicates that participants showed some characteristics of learning-to-learn, since they improved over rounds in the structured but not in the random condition.

Participants also required fewer trials to perform near ceiling during later rounds of the experiment (Fig. 5). For instance, looking at the trial number where participants first generated a reward higher than 35, *t*_*R*>35_, we found that *t*_*R*>35_ negatively correlated with round number (*r* = −0.10, *t*(115) = 5.34, *p* < .001, *d* = 0.50, *BF* > 100), indicating participants were able to achieve high scores more quickly over the course of the experiment. Importantly, this effect was stronger for the structured than for the random rounds (*t*(115) = 3.45, *p* < .001, *d* = 0.32; *BF* = 25.9).

**Figure 5.**
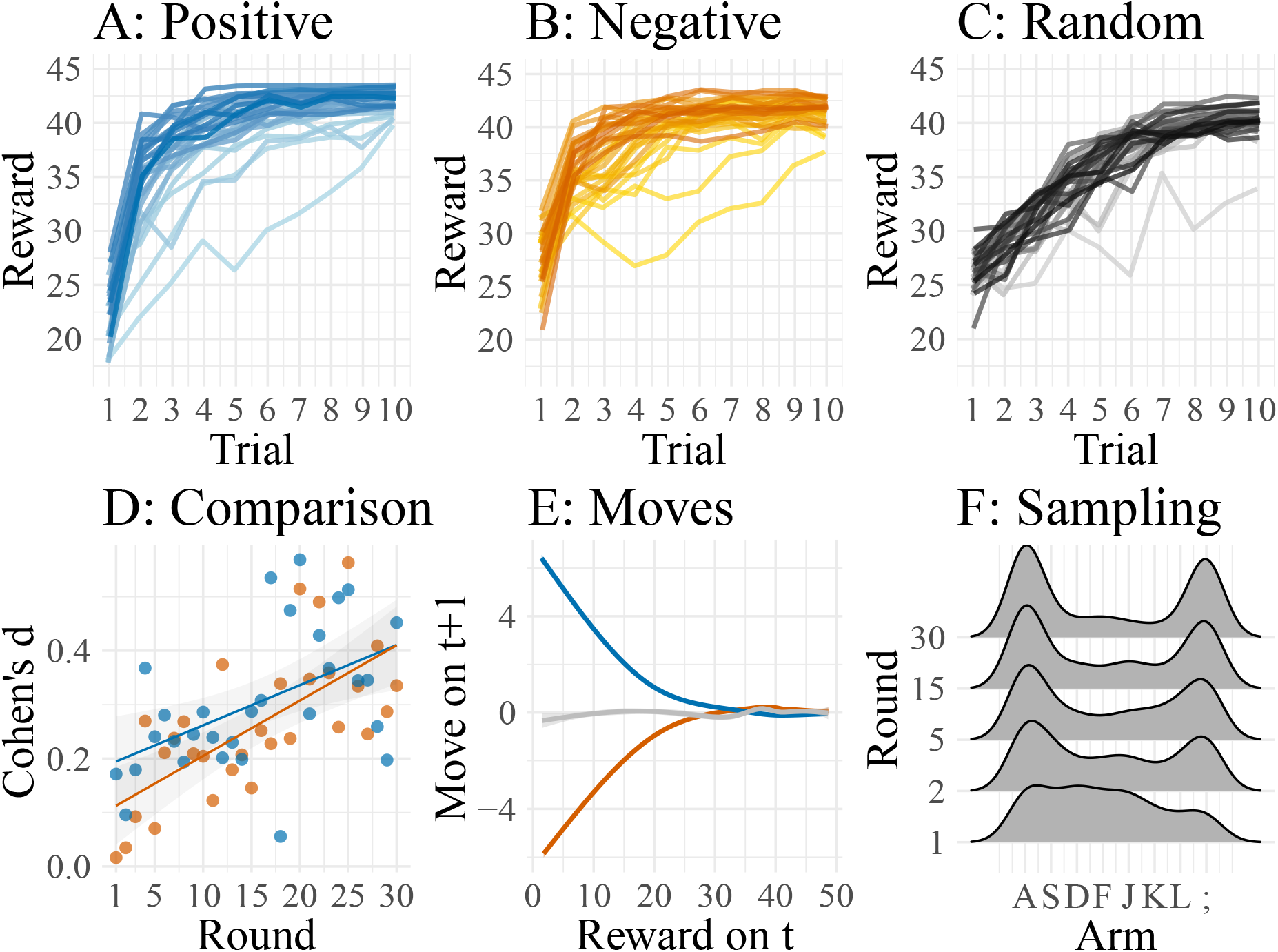
Learning-to-learn in Experiment 1. **A-C:** Mean rewards over trials averaged by round, with darker colors representing later rounds. **D:** Effect size (Cohen’s d) when comparing rewards of the linear-positive (blue) and the linear-negative (orange) condition with rewards of the random condition per round. Lines show linear regression fit. **E:** Participants’ moves on a trial *t* +1 after having observed a reward on the previous trial *t*. Lines were smoothed with a generalized additive regression model (Hastie, 2017). **F:** Kernel smoothed density of sampled arms during the first 5 trials on different rounds.

The average difference (measured by Cohen’s d) between participants’ performance during random rounds compared to the structured rounds correlated positively with round number (Fig. 5D; *r* = 0.71, *t*(28) = 6.17, *p* < .001, *d* = 4.58, *BF* > 100), indicating that participants became disproportionately better at gaining high rewards during structured rounds as the experiment progressed. Furthermore, the directionality of subsequent samples (i.e., participants’ moves) on a trial *t* +1 correlated negatively with observed rewards in the linear-positive condition (Fig. 5E; *r* = −0.71, *t*(115) = 54.5, *p* < .001, *BF* > 100) and positively in the linear-negative condition (*r* = 0.67, *t*(115) = 45.3, *p* < .001, *d* = 4.2, *BF* > 100). There was no systematic tendency to increasingly sample towards the left or right direction in the random condition (*r* = 0.02, *t*(115) = 1.44, *p* = .15, *d* = 0.13, *BF* = 0.3). Finally, we assessed how frequently participants sampled either the leftmost or the rightmost arm during the first 5 trials across rounds (see Fig 5). This analysis showed that participants sampled the leftmost and rightmost options during earlier trials more frequently as the experiment progressed, leading to a correlation between round number and the proportion of trials these two extreme options were sampled during the first 5 trials (*r* = 0.84, *t*(28) = 9.15, *p* < .001, *BF* > 100).

Taken together, Experiment 1 provides strong evidence for participants not only benefiting from latent functional structure on any particular round; it also indicates that they improved at detecting structure over rounds as the experiment progressed, i.e., they learned to learn.

## Experiment 2: Controlling for unequal variances

One potential problem when interpreting the results of Experiment 1 is that the variance of the rewards is higher for both the leftmost and the rightmost arm aggregated over rounds, compared to all of the other arms. This is because linear functions will always have their extreme outcomes on the endpoints of the input space. One way to solve the exploration-exploitation dilemma is to sample from arms with high variances (Daw, O’Doherty, Dayan, Seymour, & Dolan, 2006; Gershman, 2018; Schulz & Gershman, 2019, see also modeling section). Thus, this might serve as an alternative explanation for our results in Experiment 1. To address this possibility, Experiment 2 shuffled the middle options (between the extremes) in structured rounds, but left the outer (leftmost and rightmost) options unchanged. This removed any underlying structure, but kept the same distribution of rewards. If participants are using a heuristic based on playing the outermost options, then their performance should be mostly intact on this version of the task. But if they are using functional structure to guide exploration, then their performance should dramatically decrease.

### Participants

Ninety-one participants (36 females, mean age=32.74, SD=8.17) were recruited from Amazon Mechanical Turk and received $2.00 plus a performance-dependent bonus of up to $1.50 for their participation. The Experiment took 26.67 minutes to complete on average.

### Procedure

The design of Experiment 2 was similar to the one used in Experiment 1 (i.e., participants had to play 30 rounds of a bandit task with 10 trials per round, sampling options sequentially by pressing the corresponding key). As in Experiment 1, participants were told they had to sequentially sample from different options to gain rewards. However, instead of randomly sampling linear-positive and linear-negative functions for 10 of the 30 rounds each, we sampled from a pool of functions that was generated by taking the linear functions from Experiment 1 and randomly shuffling the rewards of the mean rewards of the middle options (i.e., all but the leftmost and the rightmost option). We refer to the shuffled linear-positive functions as *scrambled-positive* and to the shuffled linear-negative functions as *scrambled-negative* condition. The resulting pool of functions is shown in Figure 6.

**Figure 6.**
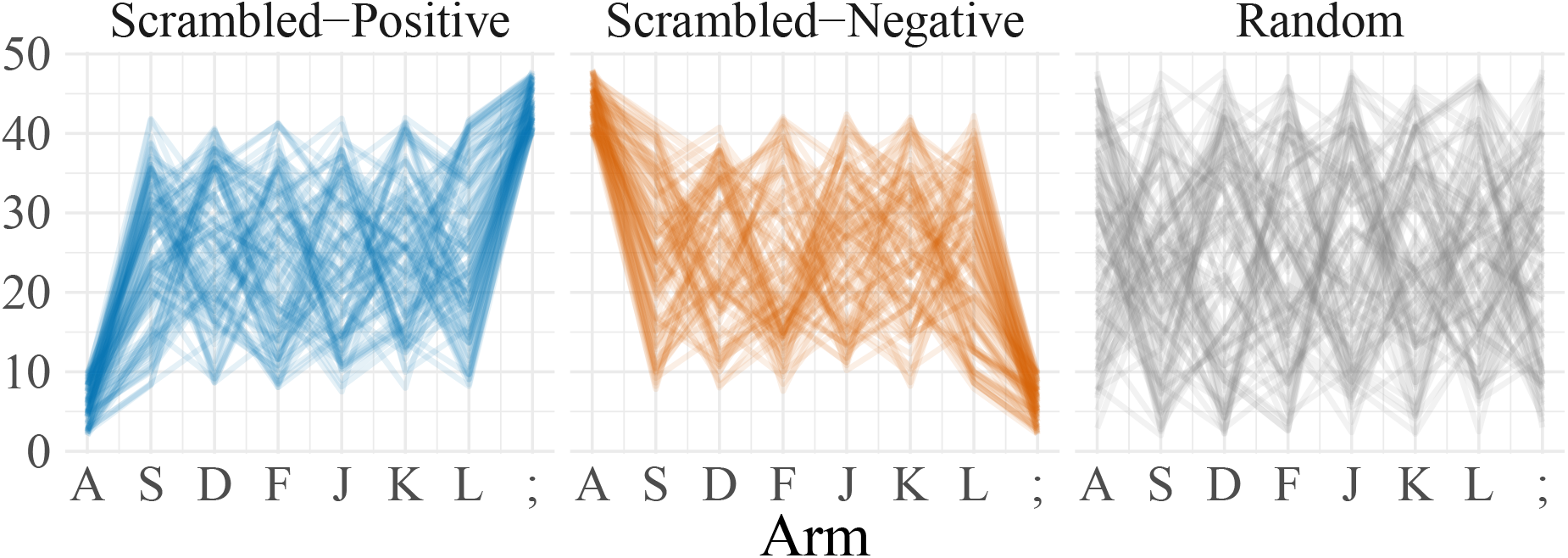
Pool of functions from which underlying structure was sampled in Experiment 2. We created 100 scrambled-positive, 100 scrambled-negative functions, and 100 random functions (50 of which were shuffled scrambled-positive and 50 of which were scrambled-negative). Scrambling a function was accomplished by taking the rewards for the middle options (i.e., S, D, F, J, K, and L) of the linear functions and randomly shuffling them to disrupt the latent functional structure.

We hypothesized that participants would still be able to detect that on some rounds the leftmost or the rightmost arm would result in the highest available reward and would potentially improve in detecting which round they were in. However, as the latent functional structure had been removed by shuffling rewards of the middle options, we expected this structural advantage to be smaller than in Experiment 1. We therefore compare the results of the two experiments, referring to Experiment 1 as the *structured* experiment and to Experiment 2 as the *scrambled* experiment. Although both experiments were conducted separately, no participant was allowed to participate in both experiments and participants were recruited from the same subject pool (i.e., Amazon Mechanical Turk). As such, these two experiments can equivalently be interpreted as a single, between-subjects experiment.

### Results and discussion

Participants performed marginally better on the scrambled-positive than on the random rounds (*t*(90) = 2.88, *p* = .005, *d* = 0.30; *BF* = 5.54; Fig. 7). Performance on the scrambled-negative rounds was better than on random rounds (*t*(90) = 5.52, *p* < .001, *d* = 0.58, *BF* > 100; Fig. 7A). There was no significant difference in reaction times between the conditions (Fig. 7B; all *p* > .05; max-*BF* = 0.04).

**Figure 7.**
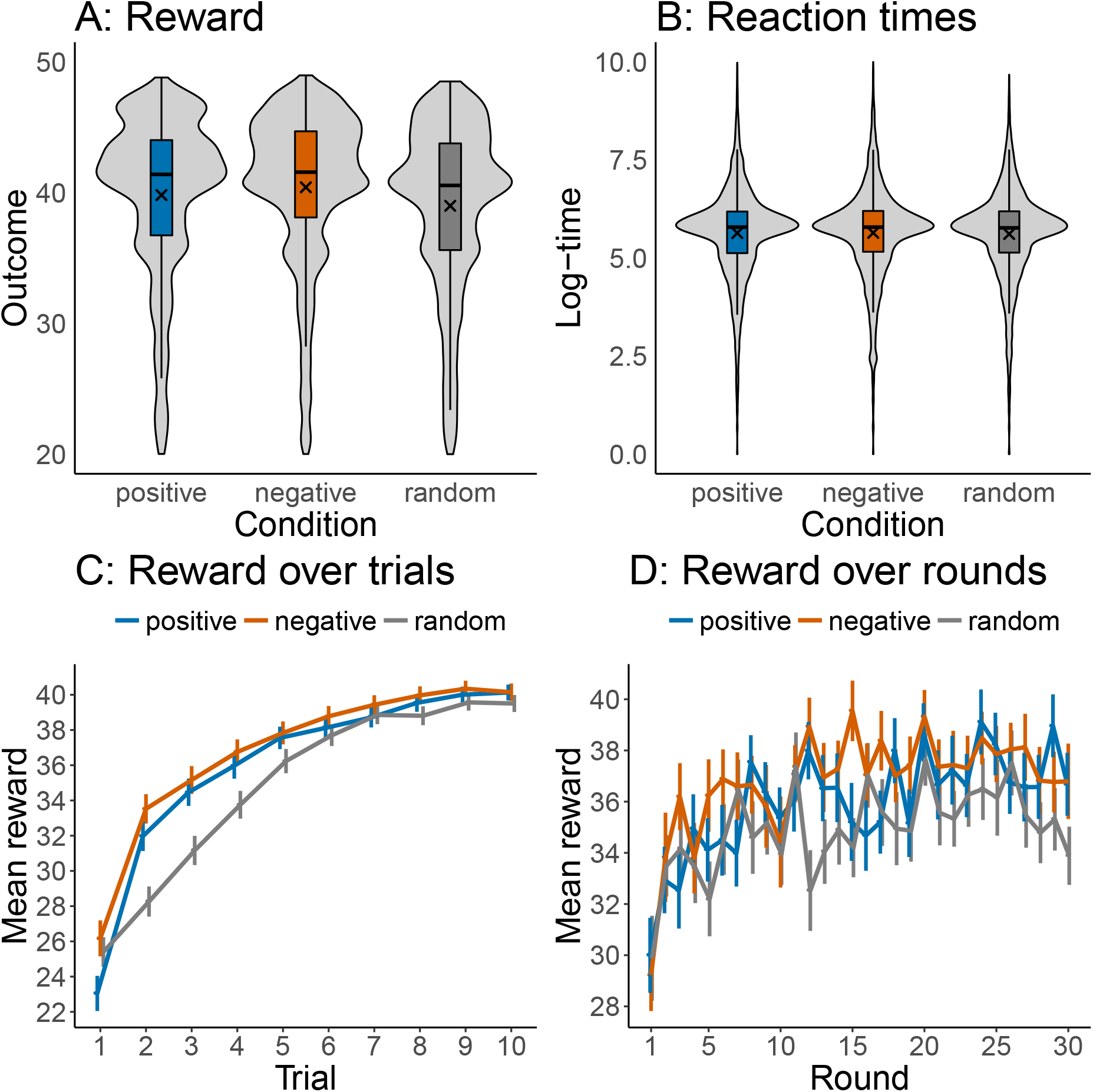
Results of Experiment 2. **A:** Box-plots of rewards by condition including violin density plots. Crosses mark the means by condition. Distributions show the rewards over all trials and rounds. **B:** Same plot but for reaction times measured in log-milliseconds. **C:** Reward over trials aggregated over all rounds. **D:** Reward over rounds aggregated over all trials. Error bars represent the 95% confidence interval of the mean.

Participants learned over trials, leading to an overall correlation between trials and reward of *ρ* = 0.29, *t*(90) = 28.63, *p* < .001, *d* = 3.00, *BF* > 100 (see Fig. 7C). This correlation was stronger for the scrambled-positive than for the scrambled-negative rounds (*t*(90) = 7.35, *p* < .001; *BF* > 100). Interestingly, this correlation was higher for the random than for the scrambled-negative (*t*(90) = 7.35, *p* < .001, *d* = 0.77; *BF* > 100) as well as the scrambled-positive rounds (*t*(90) = 3.98, *p* < .001, *d* = 0.41; *BF* > 100). Participants also improved over rounds, albeit with only a small average correlation of *ρ* = 0.09, *t*(90) = 4.65, *p* < .001, *d* = 0.49, *BF* > 100 (see Fig. 7D). There was no difference between conditions in the correlation between reward and round number (all *p* > .05; *BF* = 0.2). Thus, participants only marginally benefited from the underlying (scrambled) structure.

Assessing again the trial where participants first generated a rewards greater than 35, *t*_*R*>35_, we did not find a significant correlation with round number (*r* = −0.02, *t*(90) = 0.91, *p* = .36, *d* = 0.1, *BF* = 0.2). There was also no significant correlation between round number and the average difference between participants’ performance on random rounds and the scrambled rounds (*r* = 0.28, *t*(28) = 1.58, *p* = .13, *BF* = 1.1). Thus, participants showed no patterns of learning-to-learn in the Experiment 2 with scrambled latent functions.

We next directly compare the results of Experiments 1 and 2 (i.e., structured vs. scrambled; Figure 8). Participants in the structured experiment performed better than participants in the scrambled experiment overall (*t*(205) = 2.50, *p* < 0.01, *d* = 0.35; *BF*_10_ = 2.76), in the positive condition (*t*(205) = 3.59, *p* < .001, *d* = 0.50; *BF* = 55.8) but not in the negative condition (*t*(205) = 1.97, *p* < .05, *d* = 0.28; *BF* = 1.1). There was no difference in performance between the two random conditions (*t*(205) = 0.50, *p* = .61, *d* = 0.07; *BF* = 0.17). In sum, the linear structure in Experiment 1 helped participants to perform better, in particular in the positive condition. That the difference between the two negative conditions was not present might be due to participants frequently sampling the leftmost option first.

**Figure 8.**
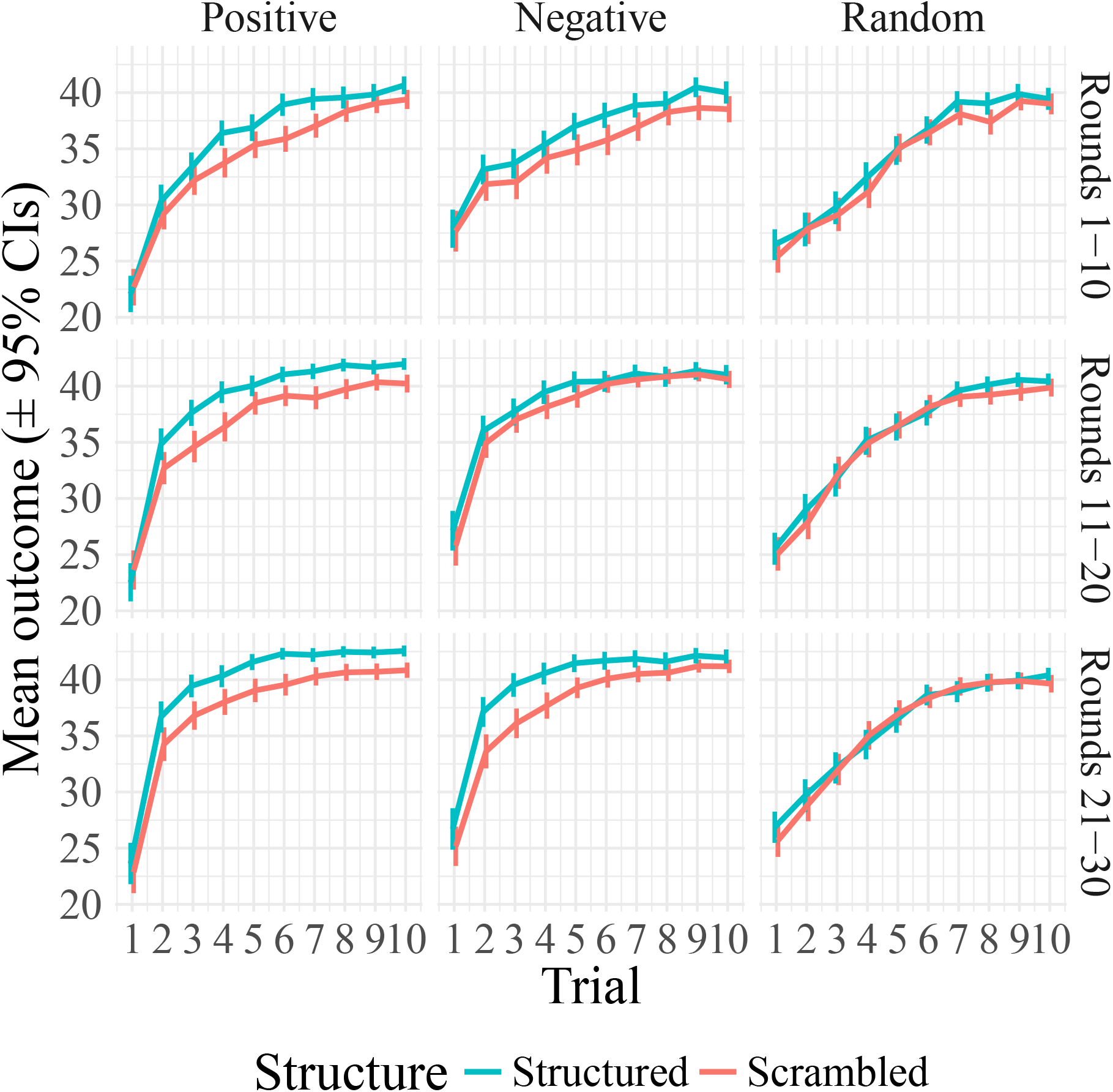
Mean reward over trials aggregated over different rounds for the structured and the scrambled experiment. Error bars represent the 95% confidence interval of the mean.

We also assessed if the two experiments differed in terms of learning-to-learn effects. Figure 9 shows the density over all 8 arms for the first 5 trials over different rounds. Participants in the structured experiment started out sampling the leftmost and rightmost arm earlier and did so more frequently than in the scrambled experiment. This intuition was confirmed by a χ^2^-test (χ^2^(1) = 52.84, *p* < .001). Furthermore, participants in the structured experiment reached rewards higher than 35 earlier on average than participants in the scrambled experiment (*t*(205) = 2.07, *p* = .02, *d* = 0.28; *BF* = 3.05). Finally, participants in the structured experiment showed a stronger correlation between round number and the difference between the random and the structured conditions than did participants in the scrambled experiment (*z* = 2.29, *p* = .01).

**Figure 9.**
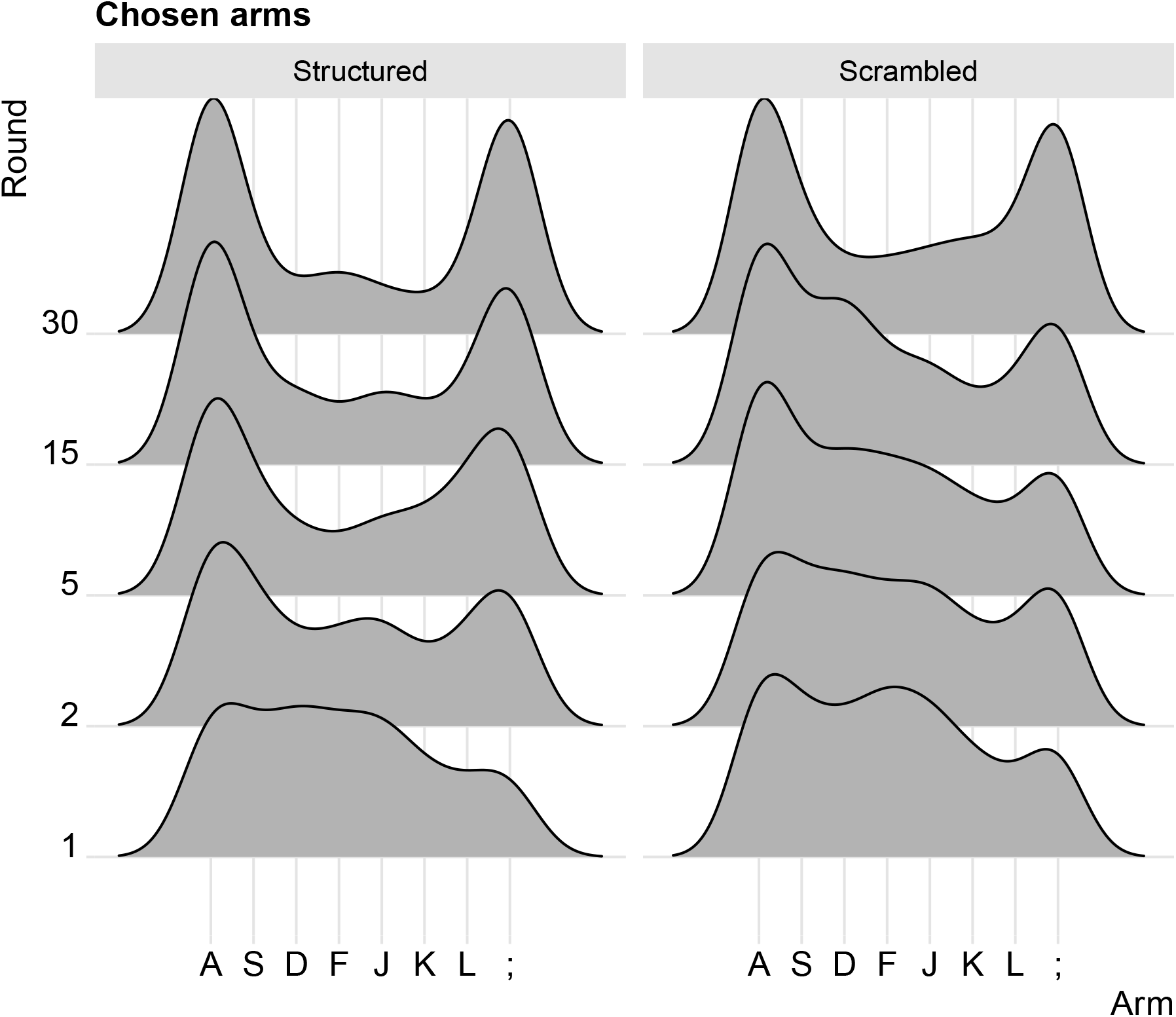
Kernel smoothed density of chosen arms during the first 5 trials for rounds 1, 2, 5, 10, 15, and 30. Left shows the density for Experiment 1 (Structured). Right shows density for Experiment 2 (Scrambled).

To summarize, we find strong evidence that participants in the structured experiment benefited from the underlying linear structure, which made them learn faster, perform better, and show stronger signatures of learning-to-learn compared to participants in the scrambled experiment. This suggests that participants are not only learning a sufficiently good policy (sampling the arms at either end) but are also using the structure within each round to aid their exploration.

## Experiment 3: Linear structure with random intercepts

Experiment 1 showed that participants can identify an underlying linear structure, and that detection improves over rounds. Experiment 2 showed that the observed behavior cannot be explained by a tendency towards sampling options with a high reward variance (i.e., leftmost and rightmost options), since learning was impaired by removing the underlying structure while keeping reward variance similar.

Another possible concern is that participants could have focused on finding arms that deliver reward points between 40 and 48, akin to a satisficing strategy frequently observed in other decision making tasks (Börgers & Sarin, 2000). Thus, despite the restricted range of maximum values it could have still been relatively easy for them to find the best possible option. Nonetheless, if participants are truly able to learn about the latent functional structure, then the actual values of the resulting rewards should not matter as much as the functional structure. We therefore assess if participants can identify and exploit the underlying structure if the experienced range of rewards is less informative in Experiment 3.

### Participants

We recruited 144 participants (71 females, mean age=34.89, SD=9.24) from Amazon Mechanical Turk and paid them a basic fee of $2.00 plus a performance-dependent bonus of up to $1.50. The experiment took 28.10 minutes on average.

### Procedure

The procedure was similar to Experiment 1, with linear-positive and linear-negative rounds randomly interleaved with rounds exhibiting shuffled rewards. However, a random intercept was sampled before each round from a uniform distribution between 0 and 50 and added to every option’s rewards. Specifically, we shifted the rewards of all options by a randomly and uniformly sampled value 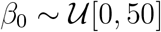 on every round. This value can also be interpreted as a random intercept that varied between rounds but was the same for every option on a particular round. For example, if *β*_0_ was 20 during a particular round, then 20 was (unknown to participants) added to every option’s value on that round.

We refer to the linear-positive functions with an added intercept as *shifted-positive* and to the linear-negative with an added intercept as *shifted-negative*. The outputs in the random condition were also shifted using the same principle as before. An example of the functions used in Experiment 3 is shown in Figure 10. Participants were told that the range of the rewards can change on every round but they would be paid based on their relative performance on each round in the end. In fact, the payment of participants’ bonus was determined without the random intercept and calculated just as in Experiments 1 and 2.

**Figure 10.**
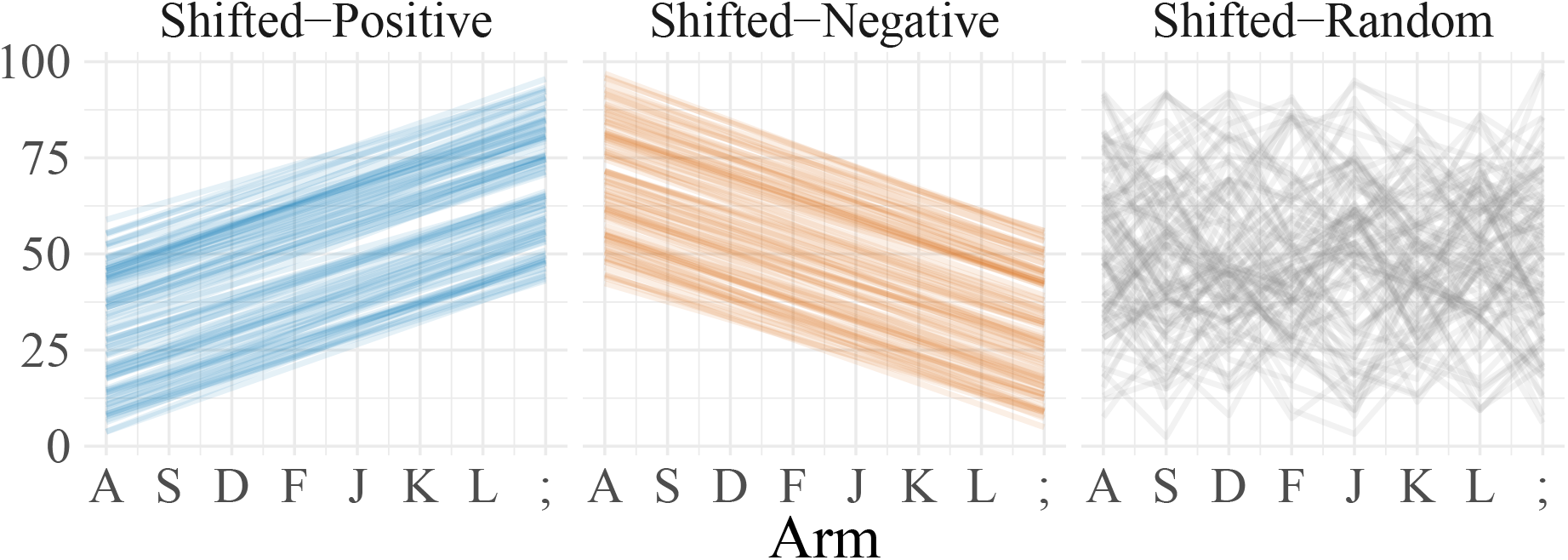
Examples of functions with underlying structure used in Experiment 3. We created 100 linear functions with a positive slope (shifted-positive), 100 functions with a negative linear slope (shifted-negative), and 100 random functions (50 of which were shuffled shifted-positive functions, 50 of which were shuffled shifted-negative functions). On each block, a random number sampled uniformly from within a range of 0-50 was added to all rewards of a current block.

### Results and discussion

As in Experiment 1, participants benefited from the underlying structure. They performed better on the shifted-positive than on random rounds (Fig. 11A; *t*(143) = 7.77, *p* < .001, *d* = 0.65; *BF* > 100) and also better on the shifted-negative than on random rounds *t*(143) = 12.49, *p* < .001, *d* = 1.04; *BF* > 100). They also performed better on shifted-negative compared to shifted-positive rounds (*t*(143) = 3.68, *p* < .001, *d* = 0.31; *BF* = 53.2), likely because they frequently sampled the leftmost arm first (see also Fig. 11C). There was no difference in reaction times by condition (Fig. 11B; all *p* > 0.05; max-*BF* = 1.4).

**Figure 11.**
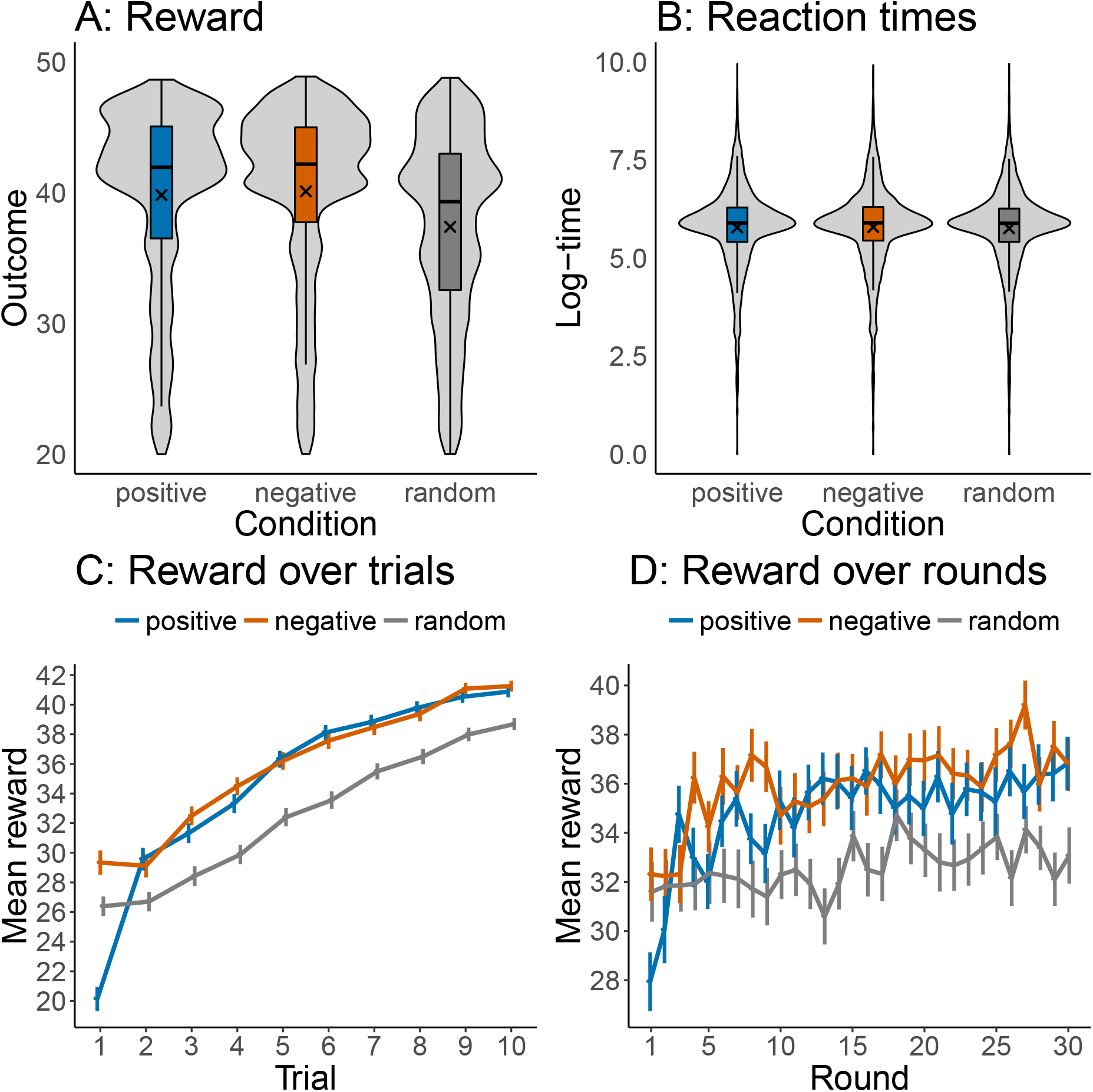
Results of Experiment 3. **A:** Box-plots of rewards by condition including violin density plots. Crosses mark the means by condition. Distributions show the rewards over all trials and rounds. **B:** Same plot but for reaction times measured in log-milliseconds. **C:** Reward over trials aggregated over all rounds. **D:** Reward over rounds aggregated over all trials. Error bars represent the 95% confidence interval of the mean.

Participants continuously improved their rewards over trials, leading to an overall correlation between rewards and trial number of *ρ* = 0.35, *t*(143) = 42.41, *p* < .001, *d* = 3.62 (see Fig. 11C). This correlation was higher for shifted-positive than for random rounds (*t*(143) = 3.17, *p* = .002, *d* = 0.26; *BF* = 11), and also higher for shifted-negative than for random rounds (*t*(143) = 3.74, *p* < .001, *d* = 0.31; *BF* = 64.4). The correlation between trials and rewards was also higher for the shifted-positive than for the shifted-negative condition (*t*(143) = 6.19, *p* < .001, *d* = 0.52; *BF* > 100). Thus, participants learned faster on structured as compared to random rounds.

Participants significantly improved their performance over rounds with a small positive correlation between rounds and rewards (*ρ* = 0.09, *t*(143) = 6.60, *p* < .001, *d* = 0.55, *BF* > 100; see Fig. 11D). This correlation was higher for the shifted-positive than for the random condition (*t*(143) = 4.71, *p* < .001, *d* = 0.39; *BF* > 100) and higher for the shifted-negative than for the random condition (*t*(143) = 3.83, *p* < .001, *d* = 0.32; *BF* = 87.8). There was no difference between the shifted-positive and the shifted-negative condition (*t*(143) = 0.97, *p* = .33, *d* = 0.08; *BF* = 0.1). This shows that, even if the intercept changes randomly on every round, participants nonetheless show learning-to-learn behavior.

Next, we further looked for signatures of learning-to-learn in participants’ behavior. We again found that participants broke above the 35 point limit (plus the added constant) earlier over time (Fig. 12A-C), with a significantly negative correlation between the first trial of breaking the 35 point limit and round number (*r* = −0.05, *t*(143) = 2.91, *p* < .01, *d* = 0.24, *BF* = 5.4). This effect was larger for the structured than for the random rounds (*t*(143) = 4.03, *p* < .001, *d* = 0.34; *BF* = 180.1). There was also a significantly positive correlation between round and the difference in rewards between the structured and the random conditions (*r* = 0.50, *t*(28) = 3.08, *p* = .004, *BF* = 12.1; Fig. 12D). Thus, participants did not only learn the latent structure on each round but also about the different kinds of encountered structures over rounds.

**Figure 12.**
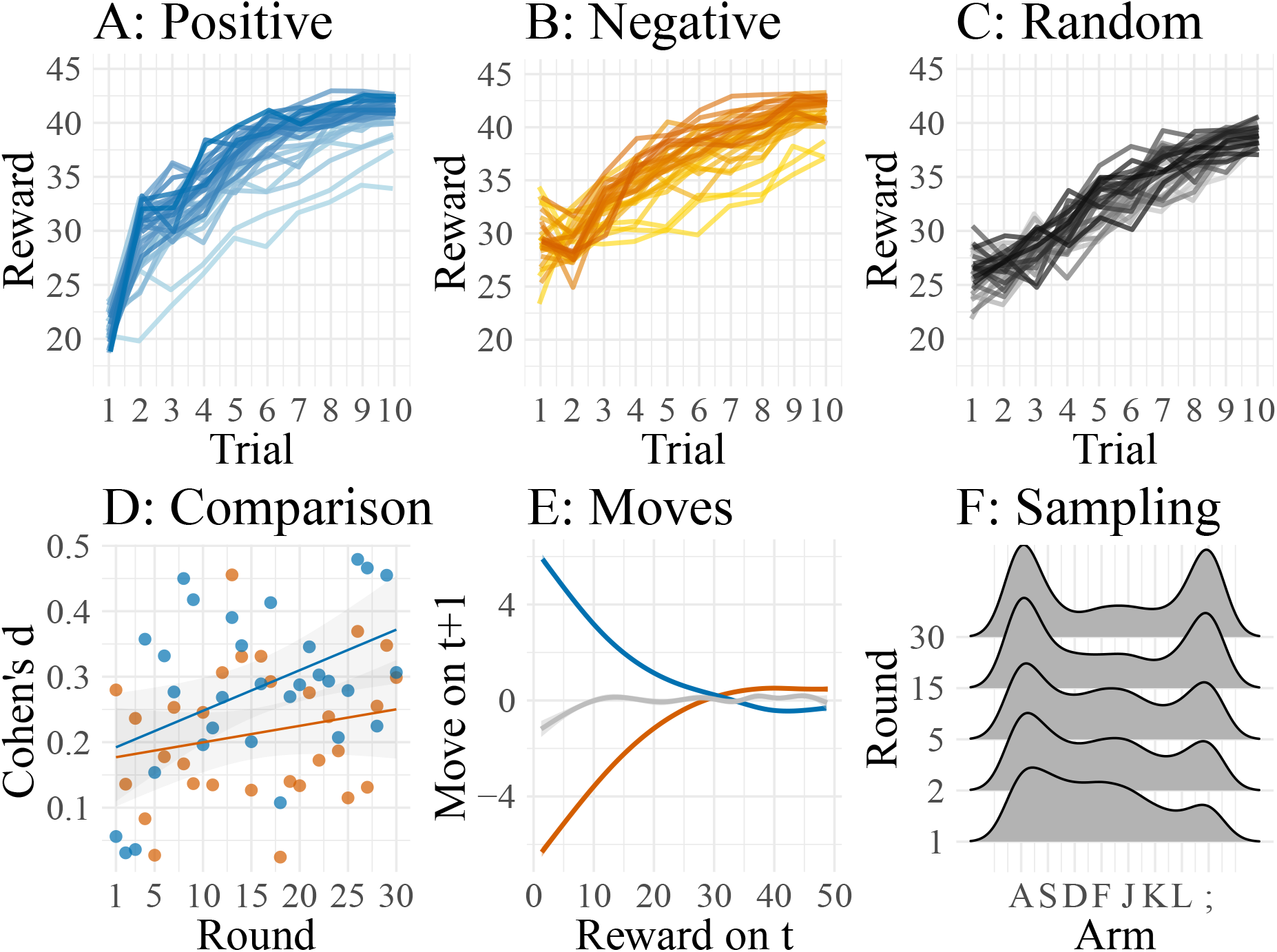
Learning-to-learn in Experiment 3. **A-C:** Mean rewards over trials averaged by round with darker colors representing later rounds. **D:** Effect size (Cohen’s d) when comparing rewards of the linear-positive (blue) and the linear-negative (orange) condition with rewards of the random condition by round. Lines indicate linear regression fits. **E:** Participants’ moves on a trial *t* + 1 after having observed a reward on the previous trial *t*. Lines were smoothed with a generalized additive regression model. **F:** Kernel smoothed densities of sampled arms during the first 5 trials on different rounds.

Next, we assessed the direction in which participants moved on trial *t* + 1 conditional on the reward encountered on trial *t* (Fig. 12E). For the shifted-positive rounds, there was a significantly negative correlation between rewards on a trial and participants’ moves afterwards (*r* = −0.62, *t*(143) = −55.2, *p* < .001, *d* = 4.6, *BF* > 100). The same correlation was significantly positive for the shifted-negative rounds (*r* = 0.61, *t*(143) = 68.08, *p* < .001, *d* = 5.7, *BF* > 100). There was also a significant positive correlation for the random rounds (*r* = 0.04, *t*(143) = 3.98, *p* < .001, *BF* > 100).

Finally, we assessed which options participants sampled during the first 5 trials on different rounds. Figure 12F shows that participants sampled the rightmost and leftmost option more frequently earlier and more frequently as the experiment progressed. This is also confirmed by correlating the round number with the proportion of how frequently participants sampled the leftmost and rightmost options during the first 5 trials (*r* = 0.89, *t*(28) = 10.06, *p* < .001, *BF* > 100).

In summary, Experiment 3 reveals that participants benefit from a latent linear structure even in a more complicated set-up in which rewards are shifted randomly on every round and only learning about the underlying structure could lead to high rewards. Participants show clear signs of learning over both trials and rounds, better performance on structured vs. random rounds, as well as some signatures of learning-to-learn.

## Experiment 4: The dynamics of structure learning

In our next experiment, we tested how people adjust to either structure or randomness in their environment over time. In particular, we wanted to know if early exposure to structured rounds could benefit performance as compared to early exposure to random rounds. Previous computational accounts of structure learning have often relied on clustering models and approximate computation that can introduce order effects into learning (e.g., Collins & Frank, 2013; Gershman, Blei, & Niv, 2010). Here, we wish to test whether people demonstrate order sensitivity. Moreover, we sought to test how participants behave if a sudden change from structured to random rounds occurred and vice versa.

### Participants

We recruited 120 participants (53 females, mean age=34.09, SD=8.98) from Amazon Mechanical Turk. Participants received $2.00 for their participation and a performance-dependent bonus of up to $1.50. The experiment took 29.36 minutes on average.

### Design

We used a two groups between-subjects design, where one group first experienced 10 structured rounds (round 1-10), then 10 random rounds (round 11-20) and finally 10 structured rounds again (round 21-30), whereas the other group experienced 10 random rounds (round 1-10), followed by 10 structured rounds (round 11-20) and 10 random rounds again (round 21-30). We refer to the first group as SRS (Structured-Random-Structured) and to the second group as RSR (Random-Structured-Random).

### Procedure

Participants’ instructions and task were the same as in all previous experiments. However, this time one group of participants experienced 10 structured rounds, followed by 10 random rounds and finally 10 structured rounds again (SRS-group), whereas the other group experienced 10 random rounds first, followed by 10 structured rounds and finally 10 random rounds again (RSR-group). For each block of 10 structured rounds, the underlying functions were sampled without replacement from the pool shown in Figure 3, such that 5 of the functions were positive-linear and 5 negative-linear functions with their order determined at random. We hypothesized that learning about structure early on would benefit participants’ performance.

### Results and discussion

Participants in the SRS-group performed better than participants in the RSR-group overall (Fig. 13A, *t*(118) = 3.34, *p* = .001, *d* = 0.62, *BF* = 26.5). This was even true when only comparing the first 20 rounds and thereby matching the number of structured and random rounds between the two groups (*t*(118) = *t* = 2.73, *p* = .007, *d* = 0.50, *BF* = 5.4). There was no difference in reaction times between the two groups (Fig. 13B, *t*(118) = 1.55, *p* = .12, *d* = 0.29, *BF* = 0.58). Participants consistently improved their rewards over trials (*ρ* = 0.39, *t*(119) = 32.40 *p* < .001, *d* = 2.96, *BF* > 100). This correlation did not differ between the two groups (*t*(118) = 1.95, *p* = .06, *d* = 0.35, *BF* = 1.02).

**Figure 13.**
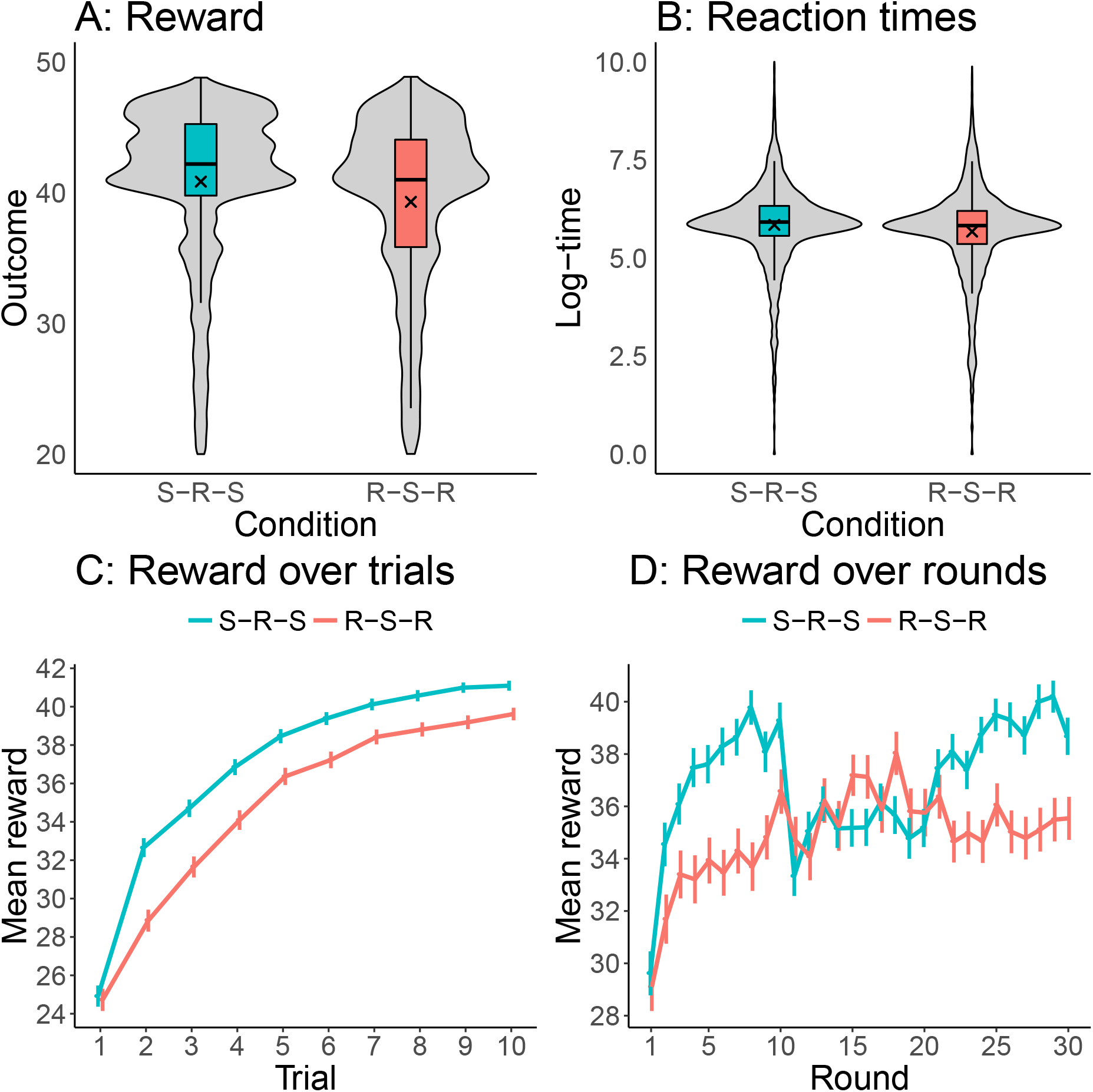
Results of Experiment 4. **A:** Box-plots of rewards by condition including violin density plots. Crosses mark the means by condition. Distributions show the rewards over all clicks and rounds. **B:** Same plot but for reaction times measured in log-milliseconds. **C:** Reward over trials aggregated over all rounds. **D:** Reward over rounds aggregated over all trials. Error bars represent the 95% confidence interval of the mean.

For the RSR-group, participants performed worse during the first 10 random rounds as compared to the subsequent second set of 10 structured rounds (*t*(50) = 4.46, *p* < .001, *d* = 0.63, *BF* > 100). They did not perform better during the 10 structured rounds (round 11-20) as compared to the second 10 random rounds (rounds 21-30; *t*(50) = 1.76, *p* = .08, *d* = 0.24, *BF* = 0.65). Finally, participants in the RSR-group also performed better during the second block of random rounds (21-30) as compared to the first block of random rounds (*t*(50) = 3.11, *p* = .003, *d* = 0.44, *BF* = 10.5). For the SRS-group, participants performed better in the first block of structured rounds (1-10) as compared to the first block of random rounds (11-20, *t*(68) = 4.50, *p* < .001, *d* = 0.54, *BF* > 100) and better in the second block of structured rounds as compared to the block of random rounds (*t*(68) = 9.22, *p* < .001, *d* = 1.11, *BF* > 100). Finally, participants in the SRS-group also performed better during the second block of structured rounds as compared to the first block of structured rounds (*t*(68) = 5.65, *p* < .001, *d* = 0.68, *BF* > 100).

Next, we compared the two groups directly. This showed that participants in the SRS-group performed better than participants in the RSR-groups during the first 10 (1-10, *t*(118) = 5.09, *p* < .001, *d* = 0.91, *BF* > 100) and the last 10 rounds (21-30, *t*(118) = 4.48, *p* < .001, *d* = 0.82, *BF* > 100). This was expected, since participants in the SRS group experienced linear latent structures during these rounds, whereas participants in RSR-group experienced random latent structures. However, participants in the SRS-group performed as well as participants in the RSR group during the ten middle rounds (11-20, *t*(118) = −1.27, *p* = .21, *d* = 0.23, *BF* = 0.4). Thus, participants who only experienced latent structure during later rounds did not benefit from it as much as participants who experienced latent structure earlier.

We examined how the two groups differed in their sampling behavior during the first 5 trials. Although participants tended to sample the extreme options more frequently even during the structured rounds in the RSR group (Fig. 14A-B), participants in the SRS group sampled the extreme option more frequently during structured rounds (*t*(118) = 4.07, *p* < .001, *d* = 0.75, *BF* > 100). Participants in the SRS group also performed better than participants in the RSR group during structured rounds (Fig. 14C; *t*(118) = 3.51, *p* < .001, *d* = 0.65, *BF* = 43.3). Finally, we checked for both groups how much a reward on trial *t* affected participants’ movements on trial *t* +1 (Fig. 14D) and found that this effect was stronger for the SRS group for both positive-linear (*t*(118) = 4.64, *p* < .001, *d* = 0.87, *BF* > 100) and negative-linear rounds (*t*(118) = 2.94, *p* = .004, *d* = 0.56, *BF* = 9.5). In summary, Experiment 4 shows that experiencing structure early on is particularly beneficial for participants’ performance. This indicates that learning-to-lean effects do not only depend on repeated encounters with structured rounds, but also on participants’ learning history.

**Figure 14.**
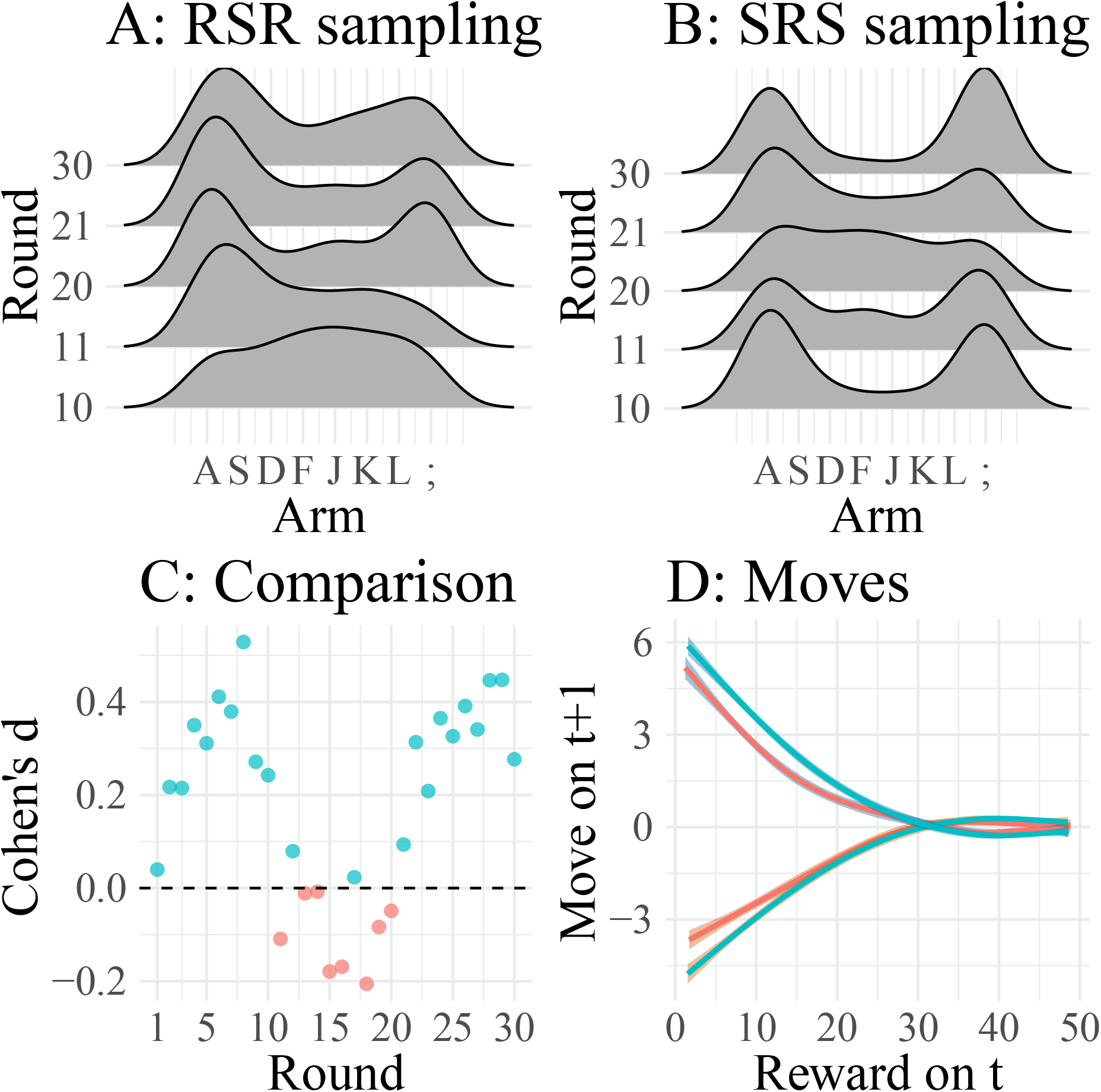
Learning-to-learn behavior in Experiment 4. **A-B:** Kernel smoothed density over sampled options during the first 5 trials on rounds 10,11,20,21, and 30 for both the RSR (A) and SRS participants (B). **C:** Effect size (Cohen’s d) when comparing rewards of the SRS and RSR condition by round. **D:** Participants’ rewards on trial *t* and distance moved on the next trial *t* +1 for both the SRS (blue) and RSR (red) participants. Lines were smoothed using a generalized additive regression model.

## Experiment 5: Change-point structure

In our final experiment, we assessed if participants are able to generalize successfully in a structured bandit task containing non-linear functions. The design is similar to our earlier experiments, but we replaced the linear functions with functions sampled from a change point kernel. A change point kernel leads to functions that increase linearly up to a point and then decrease linearly again. The key question was whether participants would discover that the functions tend to be peaked at intermediate positions and use this knowledge to guide exploration.

### Participants

We recruited 159 (62 females, mean age=32.64, SD=7.67) participants from Amazon Mechanical Turk. Participants were paid a basic fee of $2.00 as well as a performance-dependent bonus of up to $1.50. The experiment took 27.11 minutes to complete on average.

### Design

We used a two-groups between-subjects design in which participants were either assigned to a *structured* or a *scrambled* group. Whereas the structured group experienced latent functional structure that was sampled from a change point kernel, the scrambled group experienced a matched set of functions where all but the best option’s reward were randomly shuffled and therefore the latent functional structure was removed.

### Procedure

There were 30 rounds with 10 trials each as in all previous experiments. One group of participants was randomly assigned to the *structured* condition and the other group was assigned to the *scrambled* condition. In the structured condition, the latent functional structure were functions generated by a change point kernel, which produces functions with a single peak and linear slopes on either side. Thus, functions sampled from this kernel increase linearly up to a point and then sharply decreased again. In particular, 5 randomly chosen rounds would have their change point (i.e., the maximum) on the second arm, 5 on the third arm, and so forth until the seventh arm. We did not include functions with change points at the first or last arm, since these would simply be linear functions as before. For the scrambled condition, we matched the functions from the structured condition (i.e., there would be again 5 randomly chosen rounds with maxima on each of the middle arms), but randomly shuffled all but the maximum arm’s rewards in this condition. We did this to ensure that both conditions had an equal number of rounds with maxima on the same arm and therefore would be comparable. The overall pool from which we sampled functions (without replacement) is shown in Figure 15.

**Figure 15.**
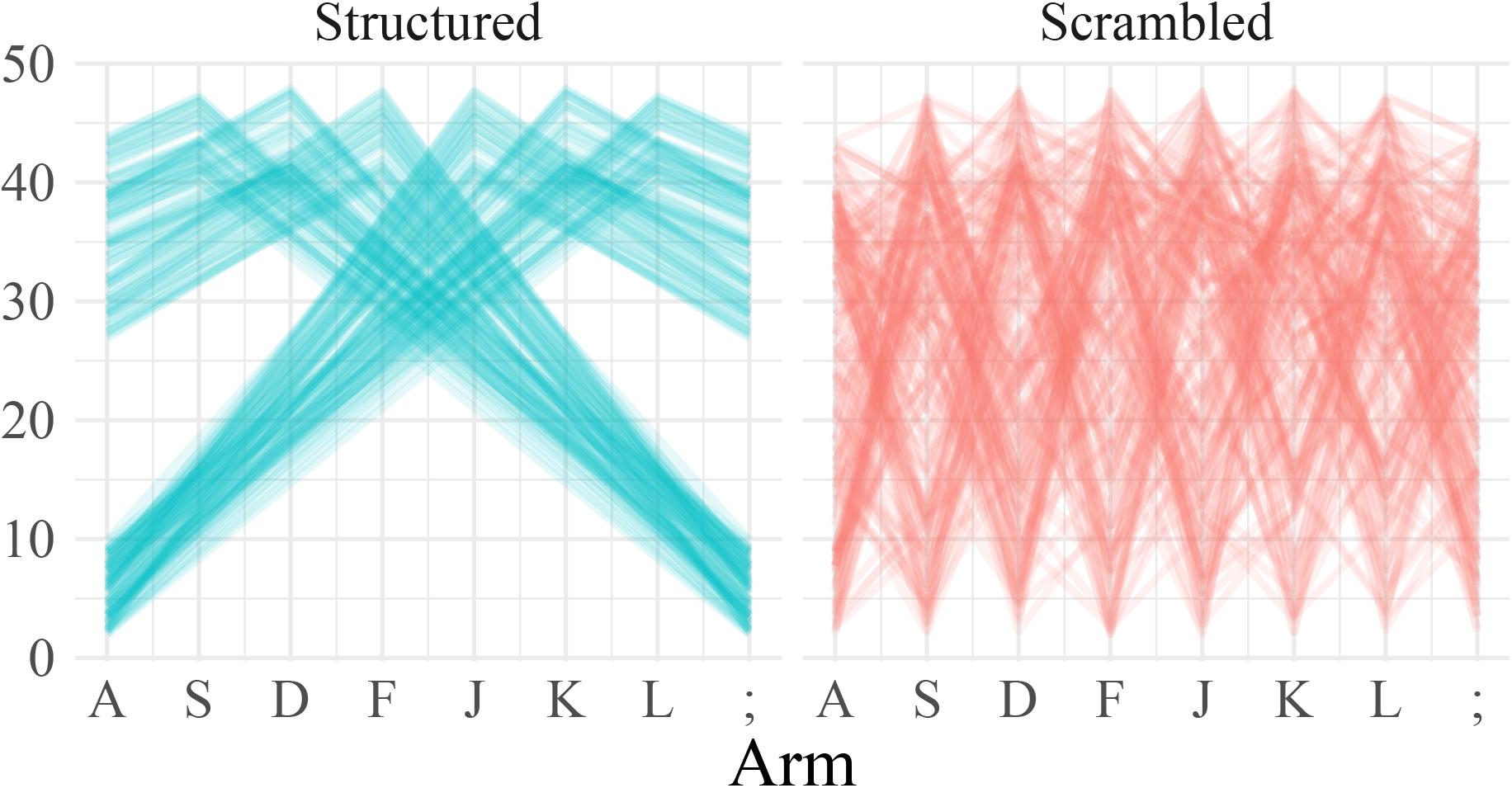
Pool of functions with change-point and scrambled structure used in Experiment 5. We created 90 change point functions, 15 for each middle arm being the overall maximum. For the scrambled functions, we matched the change point functions and scrambled the rewards for each arm that was not the maximum arm overall.

### Results and discussion

Figure 16 shows the behavioral results of Experiment 5. Participants benefited from an underlying change point structure, gaining somewhat higher rewards on average than participants in the scrambled condition (Fig. 16A; *t*(157) = 2.33, *p* = .02, *d* = 0.37; *BF* = 2.4). There was only slight evidence for a difference in reaction times between the two conditions (*t*(157) = 2.08, *p* = .04, *d* = 0.33; *BF* = 1.2).

**Figure 16.**
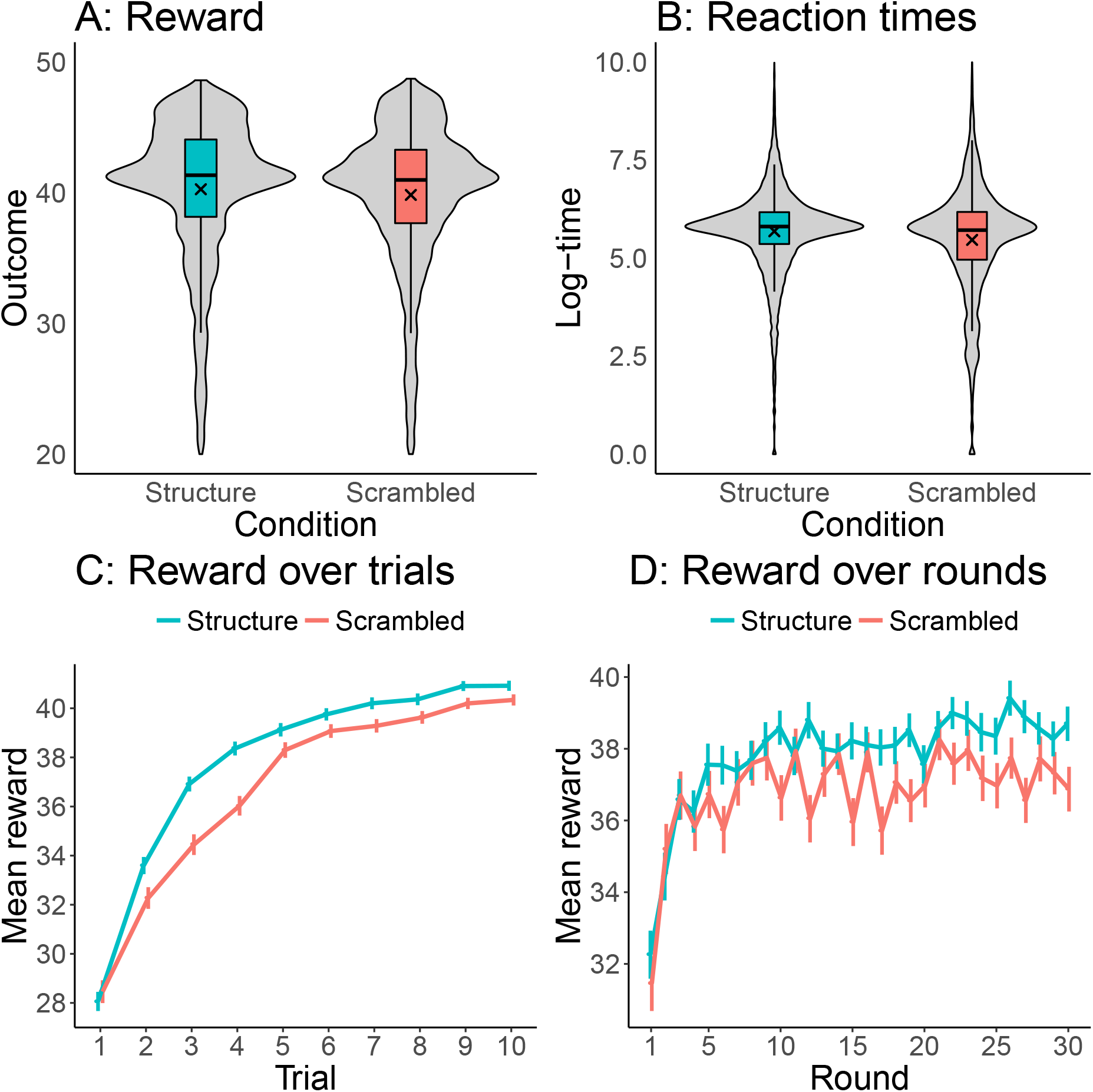
Results of Experiment 5. **A:** Box-plots of rewards by condition including violin plot of densities. Crosses indicate group means. Distributions show the rewards over all trials and rounds. **B:** Same plot but for reaction times. **C:** Reward over trials aggregated over all rounds. **D:** Reward over rounds aggregated over all trials. Error bars represent the 95% confidence interval of the mean.

Both groups improved their rewards over trials with an average correlation between trial number and rewards of *ρ* = 0.32 (*t*(158) = 36.07, *p* < .001, *d* = 2.86, *BF* > 100; Fig. 16C). This correlation was higher in the structured condition than in the scrambled condition (*t*(157) = 3.45, *p* < .001 *d* = 0.54; *BF* = 33.1). Participants also improved over rounds, yielding a positive correlation between round number and rewards of *ρ* = 0.05, *t*(158) = 4.59, *p* < .001, *d* = 0.36, *BF* > 100; Fig. 16D). This correlation was not higher for the structured compared to the scrambled condition (*t*(157) = 2.05, *p* < .05, *d* = 0.32; *BF* = 1.16).

Participants also first generated a reward higher than 35 on earlier trials as the experiment progressed, giving a negative correlation between round number and the first trial they broke the 35 reward points limit (*r* = −0.05, *t*(158) = −2.73, *p* < .01, *d* = 0.22 *BF* = 3.1). This correlation was marginally stronger for the structured condition (*t*(157) = 2.18, *p* = .03, *d* = 0.35; *BF* = 1.5). The difference between the structured and the scrambled condition increased over rounds with a correlation of *r* = 0.45, *t*(28) = 2.71, *p* = .01, *BF* = 6.0; Figure 17). Finally, we assessed how much participants improved their rewards for differently spaced moves on average (i.e., to what extent they made successful changes to better options) and found that participants in the structured condition made more successful moves than participants in the scrambled condition (*t*(157) = 4.24, *p* < .001, *d* = 0.68; *BF* > 100).

**Figure 17.**
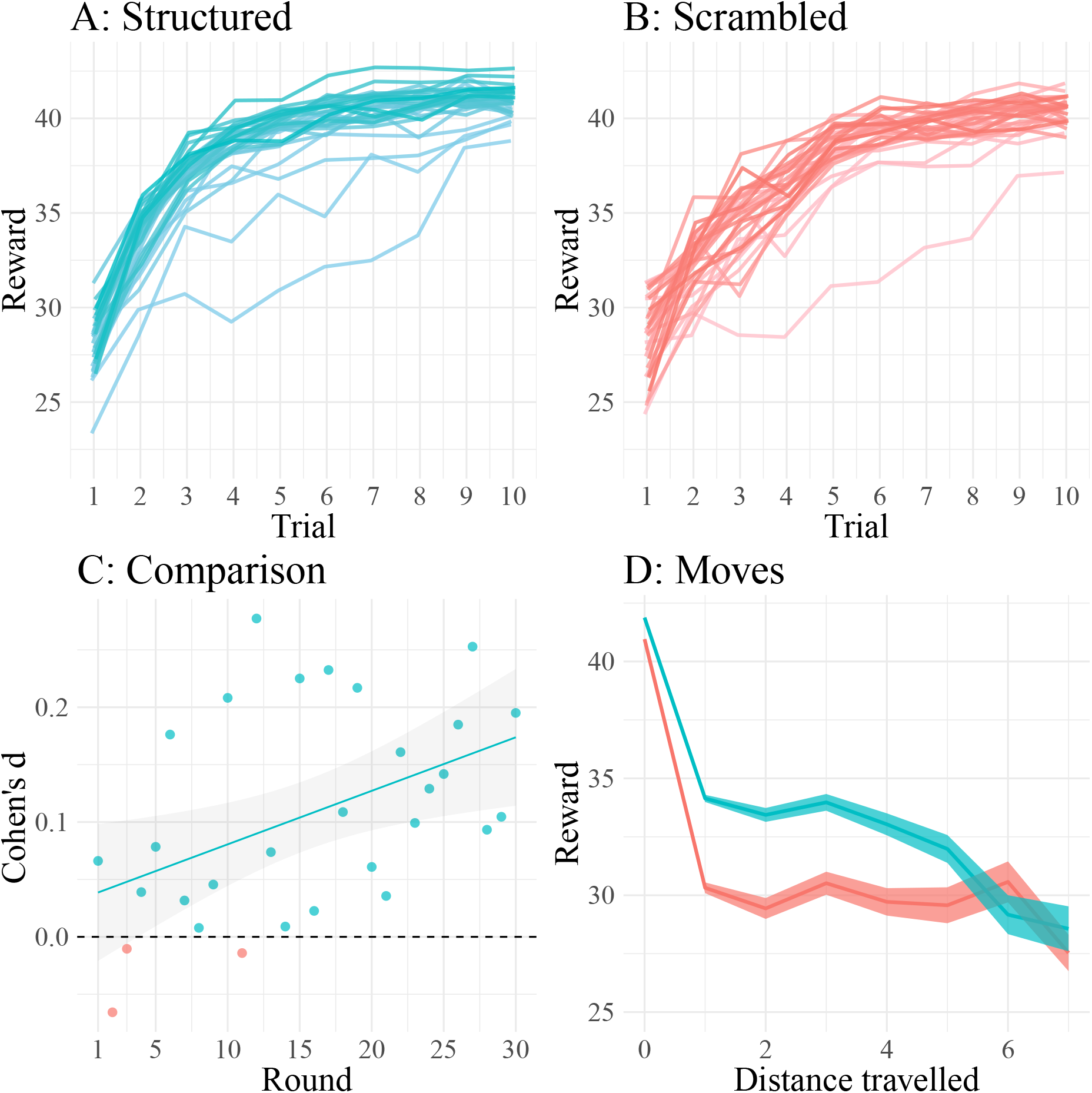
Learning-to-learn behavior in Experiment 5. **A-B:** Mean rewards over trials averaged by round with darker colors representing later rounds. **C:** Effect size (Cohen’s d) when comparing rewards of the structured and the scrambled condition by round. Line show linear regression fits. **D:** Participants’ moves and the resulting rewards (i.e., the average success of moves of different size). Lines were smoothed using a generalized additive regression model.

To summarize, Experiment 5 shows that participants can even learn about and use latent structure when the underlying function was sampled from a change-point kernel. Our results indicate that structure helped participants to learn faster over trials, and again led to a learning-to-learn effect as in the previous experiments. Even though the overall effect was weaker compared to earlier studies using similar structure (see Experiment 3 in Schulz, Konstantinidis, & Speekenbrink, 2017), it is nonetheless remarkable that participants can make use of nonlinear structure as well.

## Models of learning and decision making

We now turn to a formal model comparison contrasting several alternative accounts of learning in the structured multi-armed bandit task. These models are compared based on how well they can predict participants’ decisions in all experiments, using a Bayesian variant of leave-one-trial-out cross-validation.

### Kalman filter

The first model does not learn about the underlying functional structure, but instead updates beliefs about each option’s distribution of rewards independently. This model is called the *Kalman filter*. Kalman filter models have been used to successfully describe participants’ choices in various bandit tasks (Gershman, 2018; Speekenbrink & Konstantinidis, 2015). They have also been proposed as a model unifying various phenomena observed in associative learning (Gershman, 2015). We use the Kalman filter model as a baseline model without any generalization to compare against other models which are able to generalize across arms and rounds. It neither learns about possible latent functional structures, nor does it improve its performance over rounds (see also section on simulating human-like behavior).

Under Gaussian assumptions, a Kalman filter is a suitable model for tracking a time-varying reward function *f_n,t_*(*j*) for option *j* on trial *t* in round *n*. The mean is assumed to change over trials according to a Gaussian random walk:

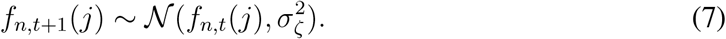

The reward functions are resampled from the prior, a Gaussian with mean 25 and variance 10, at the beginning of each round. Observed rewards are generated from a Gaussian distribution centered on the expected reward of the chosen option:

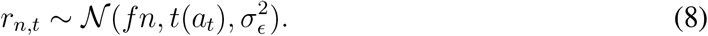

In this model, the posterior distribution of the reward function given the history of actions and rewards 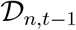 is a Gaussian distribution:

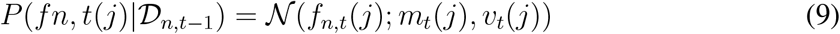

with mean

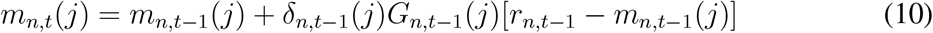

and variance

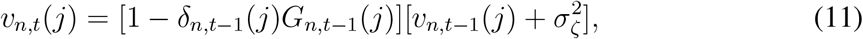

where *r_n,t_* is the received reward on trial *t* in round *n*, and *δ_n,t_*(*j*) = 1 if arm *j* was chosen on trial *t* in round *n* (0 otherwise). The “Kalman gain” (conceptually similar to a learning rate) term is given by:

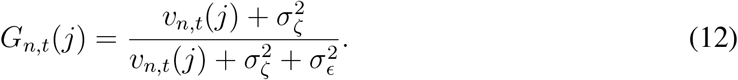

We set the diffusion and reward noise to 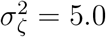 and 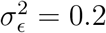, respectively. The diffusion noise models how fast options accumulate uncertainty if they are not sampled, whereas the reward noise models the assumed noise in the output of an arm.

### Gaussian Process regression

As the true generative model of the reward distributions on each round is a Gaussian Process (GP), we can also model structured function learning using the same framework (Rasmussen & Williams, 2006; Schulz, Speekenbrink, & Krause, 2018). Gaussian Process regression is a nonparametric Bayesian model of human function learning (Griffiths, Lucas, Williams, & Kalish, 2009; Lucas, Griffiths, Williams, & Kalish, 2015) that has been successfully applied as a model of how people generalize in contextual (Schulz, Konstantinidis, & Speekenbrink, 2017) and spatially correlated (Wu, Schulz, Speekenbrink, Nelson, & Meder, 2017) multi-armed bandits. We use the Gaussian Process to model structure learning and directed exploration within a round. According to a Gaussian Process regression model, structure learning is accomplished by learning about an underlying function which maps an option’s position to its expected rewards. Given the generative process described in the introduction, the posterior over the latent reward function *f* is also a GP:

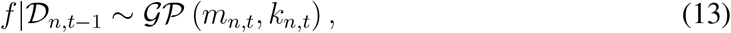

with posterior mean and kernel functions given by:

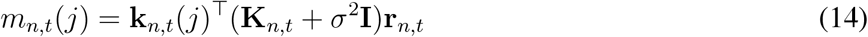

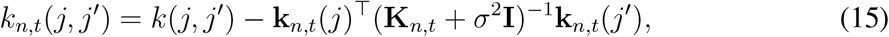

where **k**_*n,t*_(*j*) = [*k*(*a*_*n*,1_, *j*), …, *k*(*a_n, t_*, *j*)]^⊤^, **K**_*n,t*_ is the positive definite kernel matrix 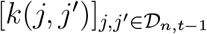, and **I** the identity matrix. The posterior variance of *f* for option *j* is:

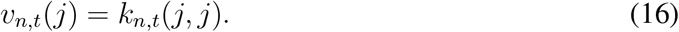

As the kernel directly corresponds to assumptions about the underlying function such as its smoothness (Schulz et al., 2015) and shape (Schulz, Tenenbaum, et al., 2017), we used two different kernels to model participants’ learning within each round. The first one was a linear kernel with parameters *θ* = (*θ*_1_, *θ*_2_):

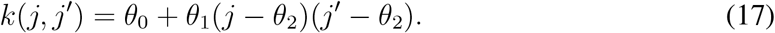

A Gaussian Process parameterized by a linear kernel is equivalent to performing Bayesian linear regression treating each option as a separate feature. This kernel therefore provided a good baseline for comparison with other more powerful regression models. The second kernel was the Radial Basis Function (RBF) kernel:

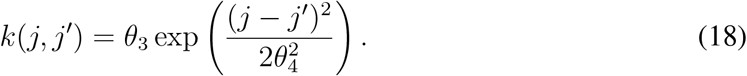

The RBF kernel is a universal function approximator that is able to asymptotically learn any stationary function provided that it is sufficiently smooth. It does that by assuming that similar (that is nearby) options will produce similar rewards. A Gaussian Process model can predict a “jump” from the leftmost option to the rightmost option after having accumulated evidence for an underlying linearly positive function of rewards over options. It does this by learning about and generalizing from the underlying structure.

### Clustering model

The Clustering model assumes that each round belongs to a cluster of rounds that share reward values. Generalization arises from inferring the assignments of rounds into clusters and inheriting previously learned statistics to speed up learning on a new round. In other words, a Clustering model can facilitate the generalization of which specific options have been good in earlier rounds, as opposed to the generalization of a class of functions (e.g., linear functions) on a current round.

Clustering models have been used to explain a variety of cognitive phenomena, including category learning (Sanborn, Griffiths, & Navarro, 2006) as well as Pavlovian (Gershman et al., 2010), operant conditioning (Collins & Frank, 2013; Lloyd & Leslie, 2013), and goal-directed navigation (Franklin & Frank, 2019). We use the clustering model to define and test a learning model which is able to generalize an option’s mean and variance over rounds. For example, knowing that there exist two scenarios in which either the leftmost or the rightmost arm produce the highest rewards can considerably speed up exploration on a new round.

Let c denote the set of cluster assignments. Conditional on data 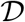, the posterior over cluster assignments is stipulated by Bayes’ rule:

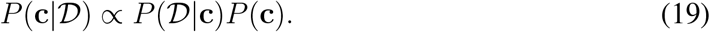

We assume a Gaussian likelihood function, where each cluster *z* is associated with a mean *μ_j,z,n_* and variance 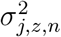 for each option *j* on round *n*. This parametrization is thus the same as the Kalman filter described in the previous section, except that (a) there is no diffusion of the mean across trials (i.e., 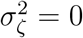), and (b) instead of resetting the means and variances to the prior at the beginning of each round, the Clustering model assumes that the means and variances can be sampled from past rounds, with some probability of generating a new cluster and resampling means and variances from the prior.

To capture the idea that participants assume a mix of reuse and resampling, we use a Bayesian nonparametric prior over clusters, known as the *Chinese restaurant process* (CRP; Aldous, 1985), that can accommodate an unbounded number of clusters, with a preference for fewer clusters. The CRP generates cluster assignments according to the following sequential stochastic process:

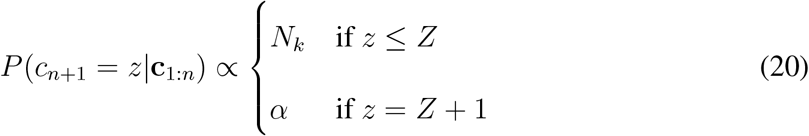

where *K* is the number of clusters that have already been created. A CRP clustering algorithm chooses the number of clusters parsimoniously based on the complexity of the encountered data (Gershman & Blei, 2012).

For inference, we use discrete particle variational inference (Saeedi, Kulkarni, Mansinghka, & Gershman, 2017) to approximate the posterior over cluster assignments. Briefly, the posterior over clustering is a discrete hypothesis space of deterministic assignments of rounds into clusters. This space is combinatorially large and we approximate this space with a fixed number (*M*) of hypotheses, or particles, that correspond to the *M* most probable under the posterior. Weights 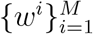 for a set of unique particles 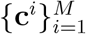 are updated on every trial such that the weight of each particle is proportional to its posterior probability. On a new round, the set of particles is augmented to generate all new possible cluster assignments for the new round consistent with the assignments of the particles. The top *M* particles (which we set to 30) are carried forward. Maximum likelihood estimation is used to estimate the mean 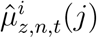 reward function and its variance 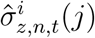 for each particle *i* and round *j* through trial *t*.

The Clustering model is used to estimate a mean and variance for each option by marginalizing over the particle approximation:

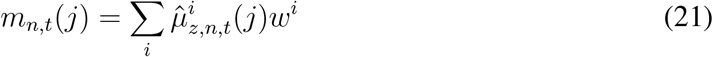

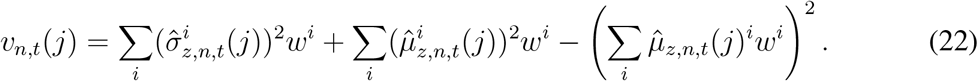

The Clustering model is sensitive to the statistics of reward for each option across rounds. As such, the Clustering model can learn that certain options (for example, the leftmost or rightmost options) have been better than other options in previous rounds and thereby guide exploration on a novel round, akin to a learning-to-learn mechanism.

### Bayesian structure learning model

Although both variants of the Gaussian Process regression model are able to learn about the underlying structure on a given round, these models assume a fixed functional form across all rounds of trials. Thus, they do not learn about the type of structure across rounds but instead, learn the specific function in each round independently. This contrasts with the Clustering model, which pools across multiple rounds to learn the task structure, and leaves the Gaussian Process regression models unable to produce the learning-to-learn patterns found in our experiments. We therefore develop another model that is able to learn across both trials and rounds through Bayesian inference over functional structures, called the *Bayesian structure learning model*. This Bayesian learner assumes there is a regular underlying function that describes the arm values but does not know which *a priori*. Instead, it infers it from data. It is important to note that participants in our task did not know about the latent structure a priori (and were not told there was a structure), but rather had to infer it.

Let *ω* denote a Gaussian Process model defined by a covariance kernel *k_ω_*. On trial *t* of block *n*, the posterior probability over models is given by Bayes’ rule:

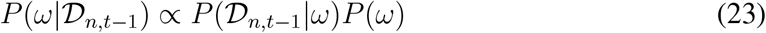

We assume an initial uniform prior *P*(*ω*) over the space of models Ω, and include three kernels, an RBF, a linear and a white noise kernel^2^, in our space of models. The RBF and linear kernel allow for function learning while the white noise kernel learns a value for each arm independently. The prior is updated progressively over time, such that the posterior probability over models on each round was used as the prior in the subsequent round. The likelihood, 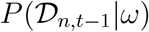, is defined by the Gaussian process and kernel for the data within each round. Thus, the model not only considers the evidence for each structure within the current round, but the accumulated evidence across rounds as well.

The model posterior is derived from a mixture of Gaussian processes. On each trial *t* in block *n*, we produce an estimate of the mean *m_n,t_*(*j*) and variance *v_n,t_*(*j*) conditioned on the posterior distribution over models, 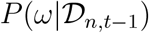:

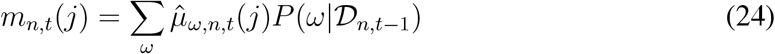

where 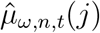 is the posterior mean for model *m* conditional on 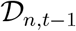. The variance is given by:

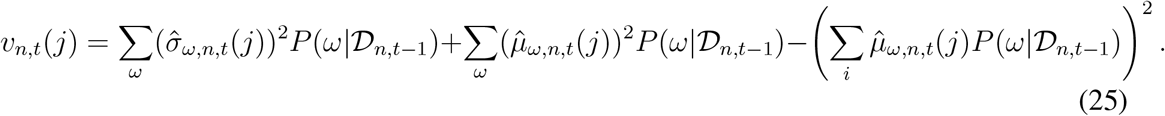

where 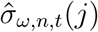 is the posterior standard deviation for model *m* conditional on 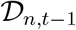.

The Bayesian structure learning model learns which kind of latent functions govern the rewards in our task over rounds and detects which structure is currently present on a given round. However, this model already approaches the task with particular structures under consideration and, importantly, with the knowledge that latent functions indeed govern the rewards in our task.

### Decision making

In order to fit the models to human choice behavior, we need to specify a policy that maps distributions of rewards to the probability of choosing a particular option. We use a variant of upper confidence bound (UCB; Auer et al., 2002) sampling as an action selection policy (Daw et al., 2006; Gershman, 2018). This adds uncertainty-guided exploration as an additional component to the model, as upper confidence bound sampling is a heuristic solution to the exploration-exploitation problem that selectively explores options that are more uncertain. Combined with a function learning model, this approach has been found to capture human behavior well in other bandit tasks that involve explicit function learning (Schulz, Konstantinidis, & Speekenbrink, 2017; Wu, Schulz, Speekenbrink, et al., 2018). It also has known performance guarantees (Srinivas, Krause, Kakade, & Seeger, 2012), and hence is normatively well-motivated.

Formally, we model the probability of choosing option *a* as a softmax function of decision value *Q_n,t_*(*j*):

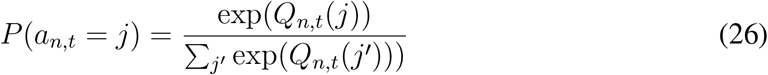

where

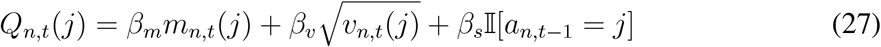

where 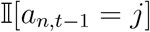 indicates whether option *j* was chosen on the previous trial; the parameter *β_s_* thus captures “stickiness” (autocorrelation) in choices. The parameters of this model define UCB exploration through a linear combination of an option’s expected means and standard deviations (Auer et al., 2002).

### Hybrid model of clustering and function learning

Our parameterization of UCB sampling as a linear combination of terms allows us to also compare combinations of different models of learning. This is particularly important because the GP models and the Clustering model rely on two different types of structure, as the GP models assume a shared structure over the arms within each round whereas the Clustering model assumes that the value of arms is structured across rounds. As such, we further consider a hybrid model that combines multiple learning rules to ask ask whether both uniquely contribute to participant behavior in this task:

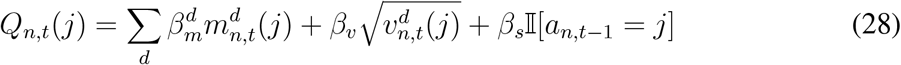

where *d* indexes different learning models.

We use the combination of the Clustering and Gaussian Process models in particular as these two mechanisms have previously been studied independently and are well-supported by the literature on learning and decision making (Collins & Frank, 2013; Franklin & Frank, 2019; Gershman et al., 2010; Schulz, Konstantinidis, & Speekenbrink, 2017; Wu, Schulz, Speekenbrink, et al., 2018). To foreshadow our modeling results, this model is indeed the winning model in all of our comparisons.

We also compare this model to two other hybrid models. The first alternative hybrid model also contains a clustering component but generalizes between arms using a linear kernel. The other hybrid model combines the GP-RBF model of generalization with the Kalman filter model. The propose of these additional hybrid models was to assess other combinations of reward tracking and generalization, and ensure that our results were not driven solely by the expressiveness of the hybrid model (see also Appendix E for additional model recovery analyses). Each of these two hybrid comparison models includes one structured component of the winning model (the means and standard deviations of either the GP-RBF or the Clustering model) and a second, plausible alternative structure learning model.

## Parameter estimation and model comparison

The parameters of the learning models are either held fixed or optimized endogenously (i.e., chosen to maximize the likelihood of the data observed so far). In particular, the diffusion and reward noise are set to 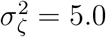 and 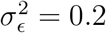 for the Kalman filter based on past empirical results (Speekenbrink & Konstantinidis, 2015) as well as simulations of model performance in our task. The hyper-parameters for the Gaussian Process regression model and the Clustering model are chosen to maximize the likelihood given the data so far.

We use hierarchical Bayesian estimation to estimate the parameters of the policy for each model, where each subject’s parameters are assumed to be drawn from a common distributions (Piray, Dezfouli, Heskes, Frank, & Daw, 2018; Wiecki, Sofer, & Frank, 2013). Formally, we assume that the *β* coefficients are distributed according to:

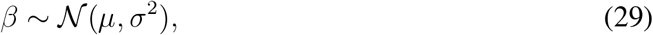

where we simultaneously estimate the group-level means *μ* and variances *σ*^2^. We place the following priors on the group-level parameters:

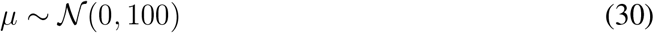

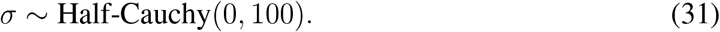

We estimate all parameters using Markov chain Monte Carlo sampling as implemented in `PyMC3` (Salvatier, Wiecki, & Fonnesbeck, 2016).

We compare the fitted models using Pareto smoothed importance sampling leave-one-out cross-validation (PSIS-LOO), a fully Bayesian information criterion (Vehtari, Gelman, & Gabry, 2017). While the details of PSIS-LOO have been covered elsewhere, briefly, PSIS-LOO can be understood as a Bayesian estimation of the leave-one-trial-out cross-validation error. It can be calculated by using weak priors and is also robust to outlier observations. It has therefore been compared favorably to other estimates such as the Watanabe-Akaike Information Criterion or weak approximations such as the Bayesian information criterion.

As a byproduct of the PSIS-LOO calculations, we also obtain approximate standard errors for every model’s estimated Bayesian cross-validation error and for comparison of errors between two models. Here, we scale the PSIS-LOO to be comparable to AIC. As with other information criteria, lower LOO-values indicate better model fits. As all participants are fitted with a hierarchical model, there is a single information criterion per model. We report the LOO for each model and its standard error, as well as the relevant pairwise comparisons (differential LOO, or dLOO) and standard error (dSE). We further report the LOO value of a chance model for comparison, which is equal to 2*N* log 8, where *N* is the total number of trials over all subject. This chance value can also be used to calculate a pseudo-*r*^2^ value for each model, similar to McFadden’s pseudo-*r*^2^ (McFadden et al., 1973):

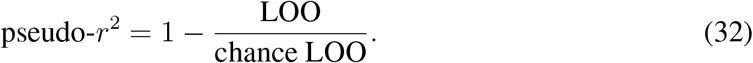

A pseudo-*r*^2^ = 0 corresponds to chance accuracy, while *r*^2^ = 1 corresponds to theoretically perfect accuracy.

We compare all of the learning models as described above in terms of their predictive performance (their PSIS-LOO values). Moreover, we create and test two additional hybrid models by linearly combining the models’ predicted means and standard deviations. Finally, we also assess a reduced “Sticky-choice” model with the influence of an option’s means and standard deviations set to zero for all arms, leaving only the sticky choice value intact; this model serves as another baseline to compare all of the other models against.

### Summary of models

We will briefly recapitulate all of the models and what predictions they make about participant behavior, before then presenting the results of our model comparison procedure. In total, we compare 10 different models based on their Bayesian leave-one-trial-out prediction error (Tab. 2).

**Table 2.** Summary of models, their components and individual predictions.

| Model | Components | Predictions |
| --- | --- | --- |
| Chance | None | All options are equally likely to be chosen. |
| Sticky-choice | Sticky choice | Previously chosen option will be chosen again. |
| Kalman filter | Mean tracking and upper confidence bound sampling | Learns about each option individually (without generalization) and uses uncertainty to explore options. |
| Linear regression | Linear kernel and upper confidence bound sampling | Learns about the underlying structure using a linear rule and uses uncertainty to explore options. |
| GP-RBF | RBF kernel and upper confidence bound sampling | Learns about the underlying structure based on the distance between options and use uncertainty to explore options. |
| Clustering model | Clustering and upper confidence bound sampling | Learns which options have been historically good and starts a round by sampling those options. Can learn-to-learn and uses uncertainty to explore. Does not generalize across options within a round. |
| Bayesian GP | Linear kernel, RBF kernel, white noise kernel, and upper confidence bound sampling | Performs Bayesian inference over each kernel within and across rounds. Can learn-to-learn and uses uncertainty to explore. |
| GP-RBF + Clustering | RBF kernel, clustering algorithm and upper confidence bound sampling | Combines similarity-based learning to generalize over options with clustering to learn-to-learn. Uses uncertainty to explore. |
| Linear + Clustering | Linear kernel, clustering and upper confidence bound sampling | Combines linear rule learning to generalize over options with clustering to learn-to-learn. Uses uncertainty to explore. |
| GP-RBF + Kalman | RBF kernel, mean tracking and upper confidence bound sampling | Combines similarity-based learning to generalize over options with simple mean tracking. Uses uncertainty to explore. Cannot learn to learn. |

The *Chance* model simply predicts that each option is equally likely to be chosen and serves as a comparative baseline. The *Sticky-Choice* model is another simple benchmark model and predicts that the previously chosen option is likely to be chosen again without any mechanism of learning. All of the other models make predictions about both the expected means and uncertainties of each option. Thus, testing whether or not uncertainties influence choice behavior tells us if participants engage in directed exploration, which would manifest itself as a significant effect of the predictive standard deviations onto choices. The *Kalman filter* model tracks rewards for each option separately. Although it does not generalize over arms at all, it is currently the best available approach to model human behavior in multi-armed bandit tasks (Gershman, 2018; Hotaling, Navarro, & Newell, 2018; Speekenbrink & Konstantinidis, 2015). We also test a class of models based on the assumption that people learn the latent function on each round. This mechanism can either be based on learning linear rules, as postulated by the *GP-Linear* model, or based on learning that similar (close by) arms will produce similar rewards, as postulated by the *GP-RBF* model. Thus, both of these models learn the functions, but differ in the underlying rules they apply. The *Clustering* model learns which arms have been historically good across rounds, and then initializes sampling on the good arms on a current round; although it can learn to sample better arms at the beginning of a round over time, it does not learn about the underlying latent functions within a round. The *Bayesian GP* model performs fully Bayesian inference over three different kernels: a linear, a RBF, and a white noise kernel. It learns how good the different kernels performed historically and on a given round. The GP-RBF and Clustering *hybrid model* uses a combination of similarity-based function learning to learn about the latent function, and clustering to learn which options have been historically good. We compare this model against two other hybrid models. The first one mixes the GP-Linear and the Clustering model, therefore testing a different algorithm of learning the latent function. The second ones mixes the GP-RBF model with a Kalman filter model to test another form of reward tracking that does not cluster arms over rounds.

### Results of model comparison

Over the suite of experiments, the best overall model was the hybrid model that combines the predictions of Gaussian Process regression, using a radial basis function kernel (GP-RBF), with the predictions generated by the Clustering model (Fig. 18; see also Table F1 in Appendix F for detailed results). Thus, generalization both within and across rounds is necessary to capture human behavior in our experiments.

**Figure 18.**
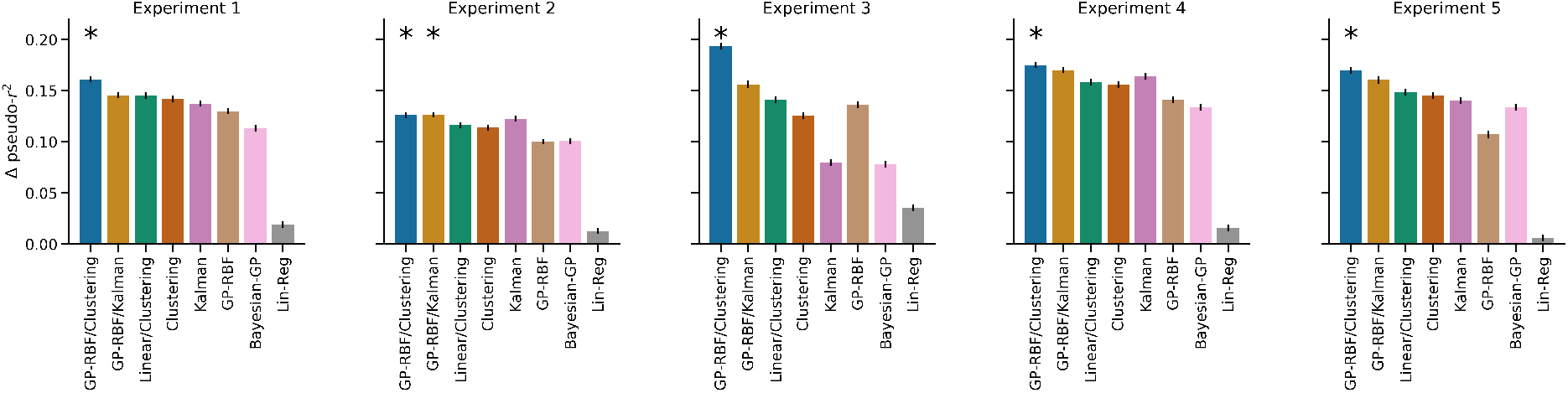
Results of model comparisons for all Experiments. Pseudo-*r*^2^ values are show in comparison to relative to a sticky-choice model. Error bars denote 95% highest posterior density interval. Best models are marked by an asterisk.

In Experiment 1, the hybrid model (LOO = 51423, pseudo-*r*^2^ = 0.61) outperformed the next best model, a hybrid combination of the GP-RBF and the Kalman filter (LOO=53652, pseudo-*r*^2^ = 0.59; see Appendix E for recovery simulations). In Experiment 2, our model comparison favored two models, a combination of the GP-RBF model and the Kalman model (LOO=43794, pseudo-*r*^2^ = 0.57) and the GP-RBF and the Clustering hybrid model (LOO=43859, pseudo-*r*^2^ = 0.57). While the GP-RBF and Kalman hybrid model predicted participants’ behavior best overall, the difference between these two models was not meaningful as the difference score was well within the standard error (dLOO = 65, dSE=133). Model comparison results for Experiment 3, in which the latent functional structure was randomly shifted on every round, again revealed that the hybrid GP-RBF/Clustering model predicted participants’ choices best (LOO = 93350, pseudo-*r*^2^ = 0.43), outperforming the next best model by a large margin (vs GP-RBF + Kalman: dLOO = 5280, dSE = 153). Experiment 4, in which participants either experienced structure early or late in the experiment, showed better model fit for the hybrid model (LOO=63591, pseudo-*r*^2^=0.45) over the next best model (GP-RBF + Kalman: dLOO = 670, dSE = 136). Finally, we see the same pattern of behavior in Experiment 5, where subjects experience change-point functions (GP-RBF/Clustering Hybrid: LOO=78336, pseudo-*r*^2^=0.4; vs GP-RBF/Kalman Hybrid dLOO=697, dSE=142).

Interestingly, subject behavior was not well-explained by the Bayesian model in any of the tasks (Figure 18). This means that participants were not best-described by a model maximally knowing about the task structure, but rather solved the task by a mix of two strategies, generalizing over the option’s rewards by applying function learning while at the same time clustering options over rounds. This is intuitive, because this hybrid model can capture participants increased learning within the structured rounds via the means of function learning as well as their learning-to-learn behavior via a method of clever reward initialization on each round. Importantly, the Bayesian model includes as a component a linear function over the arms, a model that only poorly described participants’ behavior on its own.

The GP-RBF and Clustering hybrid model also outperforms the Kalman-filter model in each of the experiments. As the only non-generalizing model in the considered set, the Kalman-filter is nonetheless a Bayesian learner that fully takes advantage of the reward and uncertainty of reward in each arm, while still treating each arm independently. Consequently, we can conclude that subjects generalize task structure and their results cannot be explained by an efficient use of data of each arm individually.

### Parameter estimates

Next we assessed the mean posterior parameter estimates of the winning hybrid model for each experiment. We examined the group-level effects with the posterior of the mean of each *β* coefficient (*μ* in equation 30). In the GP-RBF and Clustering hybrid model, there are four group effects of interest: the effect of the mean and standard deviation from both the GP-RBF and Clustering models. This allows us to tell which component of the structure learning models contributes to the hybrid models’ choice. Importantly, we can interpret significantly positive standard deviation weights as evidence for uncertainty-driven exploration. We assess statistical significance with the 95% highest posterior density interval over the parameters; if the zero is not included in this interval, we interpret this as a significant effect (J. Kruschke, 2014).

In each of the five experiments, the group-level effect of both means was significantly positive (Fig. 19, Table F1). Additionally, group-level effect of the standard deviation of the GP-RBF was also significantly positive across all of the experiments. This is consistent with previous studies showing that humans leverage both an option’s expected reward and its uncertainty to solve the exploration-exploitation dilemma (Gershman, 2018; Speekenbrink & Konstantinidis, 2015).

**Figure 19.**
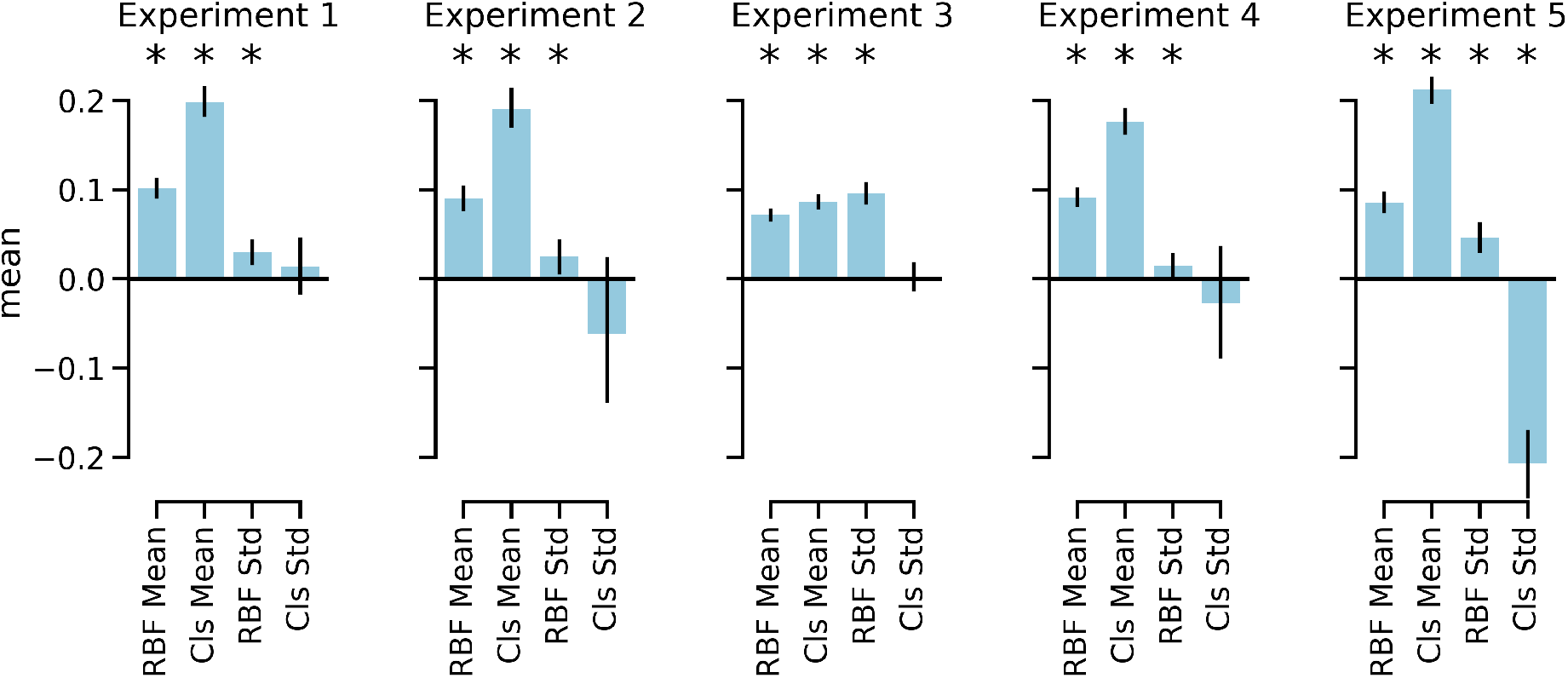
Posterior parameter estimates for the winning hybrid model for each experiment. Error bars indicate the 95% highest posterior density set.

Interestingly, we did not find evidence that the uncertainty of the clustering process positively contributed to choice behavior in any experiment. It is worth noting that while there are performance guarantees for upper confidence bound sampling in Gaussian processes (Srinivas et al., 2012), the same is not true for clustering models of generalization. Indeed, while clustering models of generalization have substantial empirical support in a variety of cognitive domain (Collins & Frank, 2013; Franklin & Frank, 2019; Gershman & Niv, 2010; Lloyd & Leslie, 2013; Sanborn et al., 2006), in reinforcement learning domains the performance of clustering models can be arbitrarily bad if it misspecifies the task domain (Franklin & Frank, 2018), and it is not clear if using the uncertainty of the Clustering model is a normative strategy. This is most apparent in Experiment 5, where clustering uncertainty related negatively to choice value. In this experiment, the standard deviation of reward across rounds in Experiment 5 was highest for the arms at the end of the keyboard (Figure 15). These arms were also never the highest rewarded arm. Because the clustering model is sensitive to the statistics of reward across rounds, the negative clustering uncertainty bonus may reflect the adaptive strategy of avoiding the arms of at each end.^3^

### Comparing to the Bayesian Structure Learning model

It might be somewhat surprising that the Bayesian structure learning model did not provide the best account of human behavior in our experiments. When we examine the trial-by-trial model fit (i.e., PSIS-LOO) in Experiment 1, the Bayesian structure learning model has a higher PSIS-LOO (i.e., worse model fit) across all trials when compared to the the hybrid GP-RBF and clustering model (Figure 20A), thus providing a worse account for subject choice behavior. This difference between the two models is largest in the first few trials of each round, which are trials that are likely linked to exploratory choices.

**Figure 20.**
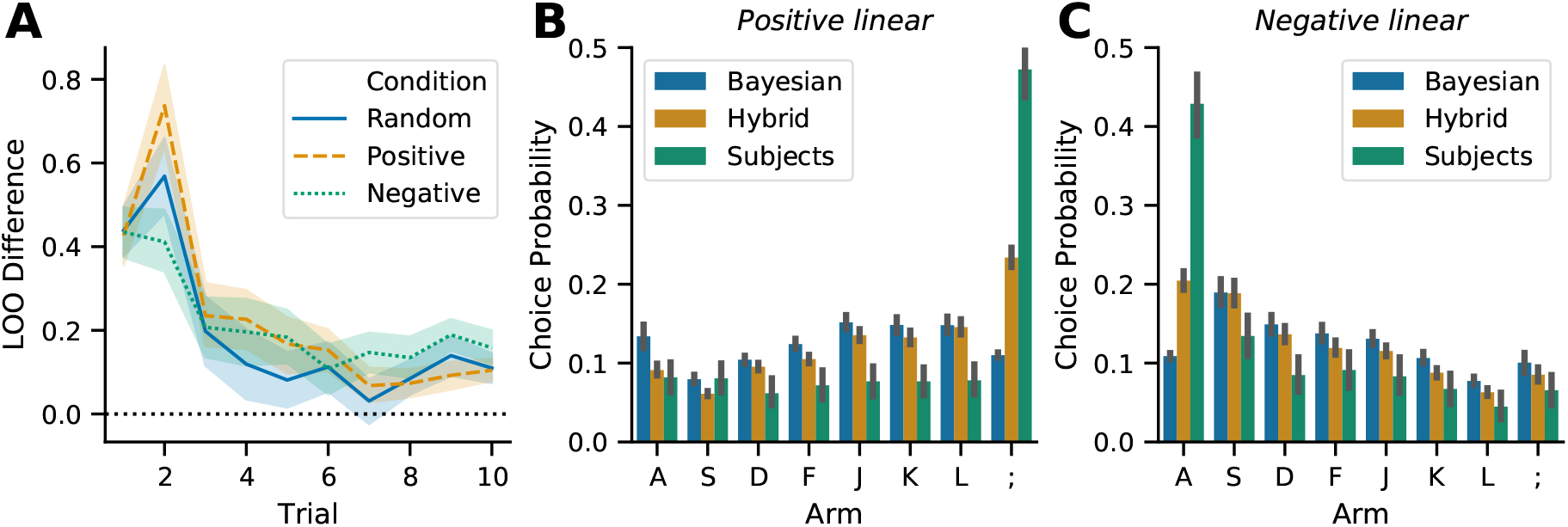
Experiment 1: Bayesian Structure Learning Model **A.** Difference in model fit (LOO) between Bayesian structure learning model and hybrid GP-RBF, Clustering model across trials in all rounds. Positive value reflect greater evidence for hybrid model. **B-C.** Choice probability in trial 2 following a non-optimal response shown the positive (**B**) and negative (**C**) linear conditions.

Why is this the case? In Experiment 1, an efficient strategy is to choose either the leftmost or rightmost arm and then continue to choose that arm if it is highly rewarded and switch to the other end if it is not. This pattern of behavior can be seen in subject data by looking at trial 2 following a sub-optimal response in trial 1 (Figure 20B,C). Across all rounds, subjects transition to the optimal response with high frequency (Figure 20B,C), switching from the leftmost arm to the rightmost arm in the positive linear condition in 47.2% of relevant trials and switching from the rightmost arm to the leftmost arm 42.8% of relevant trials. The hybrid GP-RBF and clustering model shows a similar pattern, albeit to a lesser degree (optimal transitions, positive linear: 23.3%; negative linear: 20.5%), whereas the Bayesian model fails to make these optimal transitions above chance rates (optimal transitions, positive linear: 11.0%; negative linear: 10.9%), instead tending to make transitions of sub-optimal length. This is likely the case because the Bayesian model does not immediately consider the other extreme option first, but rather averages the predictions of structured and white noise kernels, leading to less extreme transitions than what is empirically observed. Hence, we find the hybrid GP-RBF model captures this pattern of structured exploration found in subject data better than the Bayesian model^4^.

### Generative simulations

Since the hybrid model combining the Clustering and the GP-RBF model was the best model across all of our model comparisons, we examine the generative performance of this model to ensure that the observed qualitative behavioral patterns found in the human data can also be produced by the model. As a comparison, we also examine the generative performance of the Kalman filter model. This comparison highlights which aspects of human-like behavior are uniquely captured by the generalization mechanisms of the hybrid model. It also facilitates testing whether the two models can be recovered with our model comparison procedure. We focus on the Kalman filter as an alternative model as it is the standard model applied in multi-armed bandit tasks (Gershman, 2015; Navarro, Tran, & Baz, 2018; Speekenbrink & Konstantinidis, 2015).

#### Simulating human-like behavior for Experiment 1

We first assess whether both the hybrid and Kalman models could successfully generate human-like behavior, a variant of posterior model falsification (Palminteri, Wyart, & Koechlin, 2017), in Experiment 1. We simulate 100 participants in Experiment 1 with both models. For the Kalman filter model, we combine the predicted means and standard deviations to an upper confidence bound by using equal weights (both *β_m_* = 1). For the hybrid model, we combine the GP-RBF model’s predicted means and standard deviations with the Clustering model’s predicted means to create each options utility, again by setting all *β_m_* = 1. We do not use the Clustering model’s predicted uncertainty as the effect of this component was not reliably different from 0 across all our model comparisons. We also do not add a stickiness parameter into this simulation as we wanted to see what the behavior of the models would look like without this nuisance parameter.

Figure 21 shows the results of the Kalman filter model simulation. The Kalman filter model did not produce higher rewards in the linear-positive rounds than in the random rounds (*t*(99) = 1.08, *p* = .28, *d* = 0.10, *BF* = 0.2), nor did it produce higher rewards in the linear-negative rounds than in the random rounds (*t*(99) = −0.12, *p* = .90, *d* = 0.01, *BF* = 0.1). Although the Kalman filter model generated better rewards over trials (*ρ* = 0.37, *p* < .001), it did not significantly improve over rounds (*ρ* = 0.01, *p* = .18). Indeed, the Kalman filter showed no indicators of learning-to-learn-effects; there was no significant correlation between rounds and the difference between structured and random rounds (both *p* > .05), no significantly larger moves in the correct directions after observing low rewards (see Fig. 21E), and no significant increase in how frequently it sampled the leftmost or rightmost options across rounds (*r* = 0.23, *p* > .22). Thus, the Kalman filter model cannot generate human-like behavior in our task.

**Figure 21.**
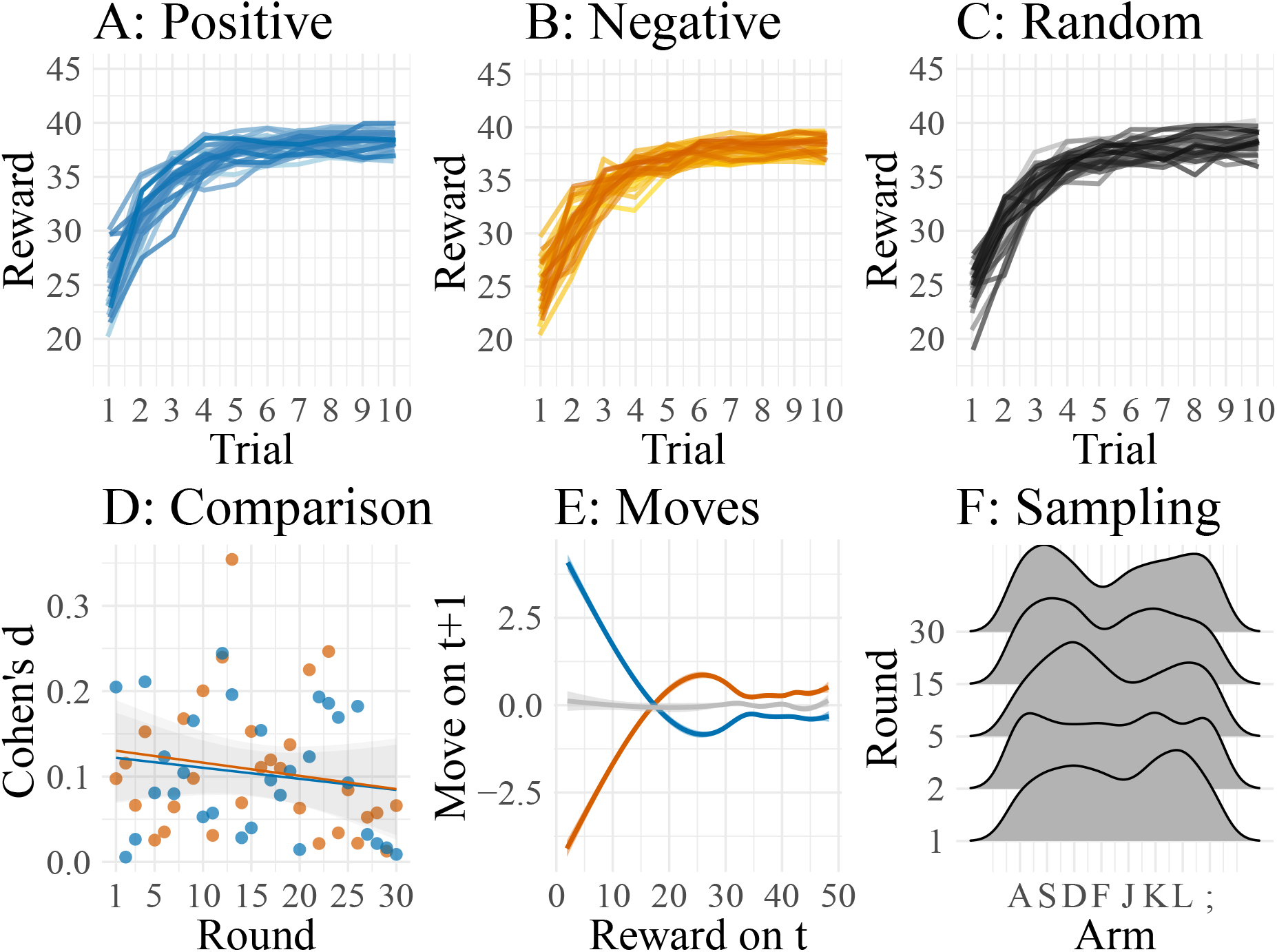
Simulated behavior of the Kalman filter model. **A-C:** Mean rewards over trials averaged by round with darker colors representing later rounds. **D:** Effect size (Cohen’s d) when comparing rewards of the linear-positive (blue) and the linear-negative (orange) condition with rewards of the random condition per round. Lines indicate means of linear least-square regressions. **E:** Participants’ moves on a trial *t* + 1 after having observed a reward on the previous trial *t*. Lines were smoothed using a generalized additive regression model. **F:** Kernel smoothed density of sampled arms during the first 5 trials on different rounds.

Figure 22 shows the results of the hybrid model. The hybrid model produced higher rewards in the linear-positive (*t*(99) = 4.67, *p* < .001, *d* = 0.47, *BF* > 100) and the linear-negative (*t*(99) = 5.33, *p* < .001, *d* = 0.53, *BF* > 100) rounds than in the random rounds. Furthermore, the hybrid model generated higher rewards over both trials (*ρ* = 0.39, *p* < .001) and rounds (*ρ* = 0.34, *p* < .001). Just as in the human data, the hybrid model also showed signatures of learning-to-learn behavior as the effect sizes when comparing the structured and the random rounds increased over time (*r* = 0.44, *p* < .001), the model moved in the right directions after observing low rewards (in particular for the linear-positive rounds; see Fig. 22D), and sampled the leftmost and rightmost options during earlier trials more frequently over time (*r* = 0.75, *p* < .001). We therefore conclude that the hybrid model is an acceptable generative model of human-like behavior in our task.

**Figure 22.**
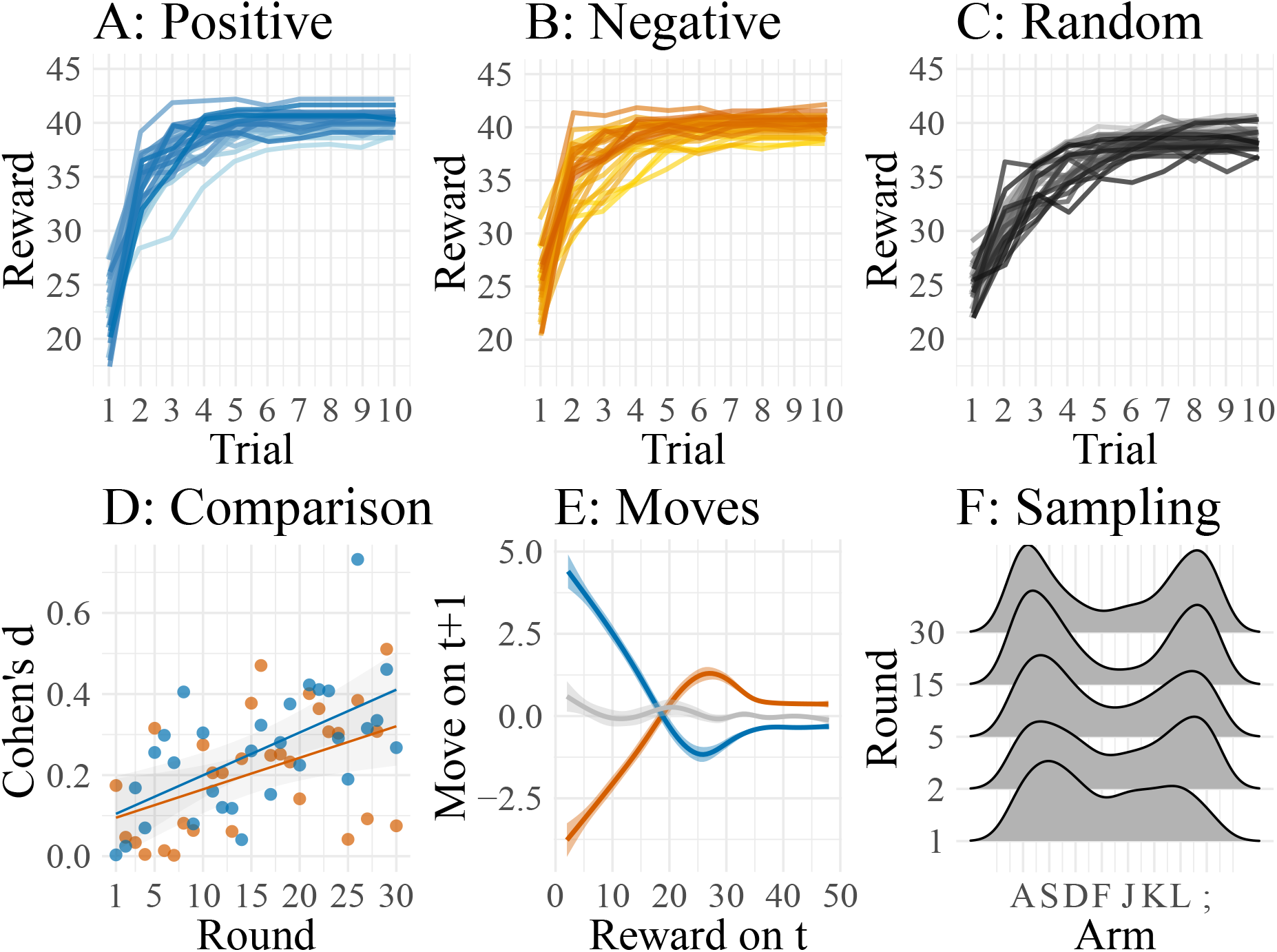
Simulated behavior of GP-RBF/Clustering hybrid model. **A-C:** Mean rewards over trials averaged by round with darker colors representing later rounds. **D:** Effect size (Cohen’s d) when comparing rewards of the linear-positive (blue) and the linear-negative (orange) condition with rewards of the random condition per round. Lines indicate means of linear least-square regressions. **E:** Participants’ moves on a trial *t* + 1 after having observed a reward on the previous trial *t*. Lines were smoothed using a generalized additive regression model. **F:** Smoothed density of sampled arms during the first 5 trials on different rounds.

#### Model recovery for Experiment 1

Next, we assess the recoverability of our generative modeling. We do this to validate our model comparison procedure and to account for the possibility of over-fitting. Specifically, we want to rule out the possibility that the hybrid model would best fit any dataset as a consequence of its expressiveness and the Bayesian parameter estimation (see Appendix F for further recovery simulations). Thus, the model recovery consists of simulating data using a *generating model* specified as the two models described before. We then perform the same model comparison procedure to fit a *recovering model* to this simulated data.

The results of this model recovery analysis are shown in Figure 23. When the hybrid model generated the data, the hybrid model achieved a predictive accuracy of *r*^2^ = .66 and LOO of 18570.6, whereas the Kalman filter model only achieved a LOO of 34570.6, with an average predictive accuracy of *R*^2^ = .37. When the Kalman filter generated the data, the hybrid model achieved a predictive accuracy of *r*^2^ = 0.27 and a LOO of 39871.6, whereas the Kalman filter model achieved a predictive accuracy of *r*^2^ = 0.71 and a LOO of 15839.5.

**Figure 23.**
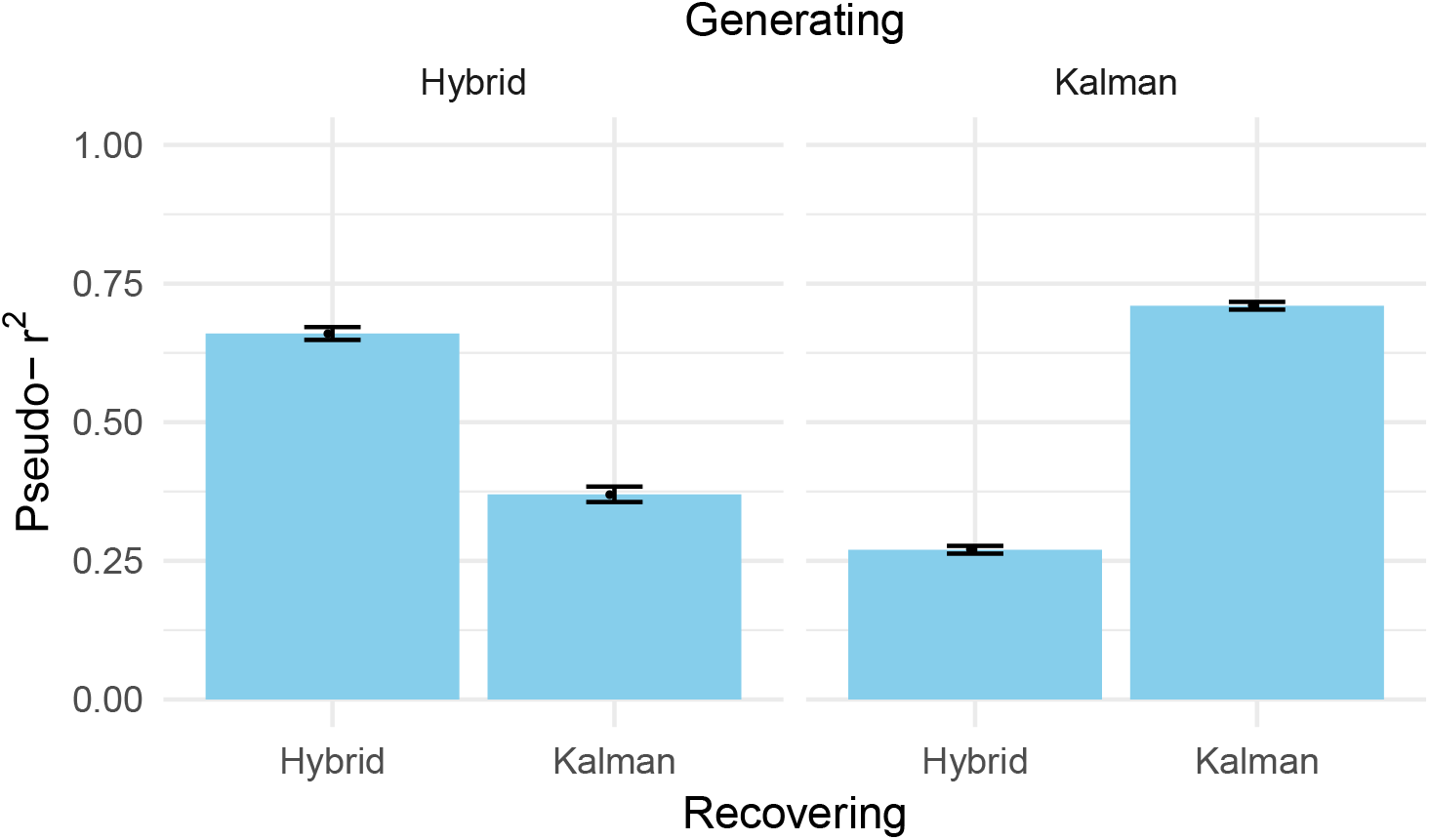
Results of the recovery simulation. Left: fits to data generated by the hybrid model. Right: fits to data generated by the Kalman filter. Error bars represent the 95% highest density intervals.

In summary, we find evidence that the models are discriminable, and that the hybrid model is unlikely to overfit data generated by the wrong model. Both models were thus recoverable.

#### Simulating human-like behavior for Experiment 4

We are also interested if the hybrid model can capture other effects observed in our experiments, in particular the dynamic effects of structure learning observed in Experiment 4. One of the main findings of Experiment 4 was that participants’ performance in later rounds was sensitive to the structure in earlier rounds. The SRS group performed better than the RSR group during the first and last 10 structured rounds and, importantly, performed as well as the RSR group during the 10 random rounds from round 11 to round 20. This means that learning-to-learn effects are not simply a result of repeatedly encountering structure, it also matters at what time during learning structure first appears.

We assess if the hybrid model is able to reproduce this effect by simulating behavior for Experiment 4. Doing so, we simulate 50 participants for the SRS condition and 50 participants for the RSR condition. The results of this simulation are shown in Figure 24.

**Figure 24.**
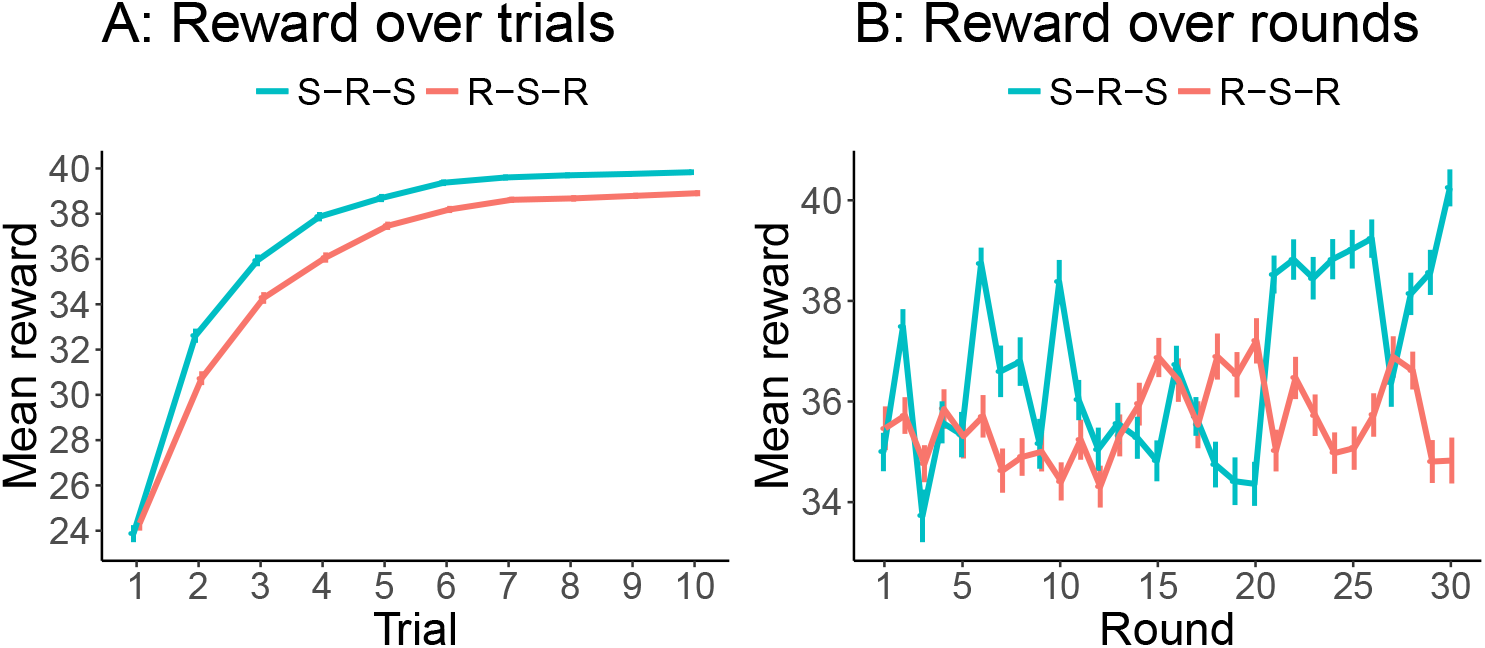
Results of the hybrid model simulation for Experiment 4. **A:** Average reward over trials. **B:** Average reward over rounds. Error bars represent the 95% confidence interval.

The simulated SRS participants performed better than the RSR participants overall (*t*(98) = 4.5, *p* < .001, *d* = 0.89, *BF* > 100), during the first 10 rounds (*t*(98) = 2.89, *p* = .004, *d* = 0.58, *BF* = 8) and during the last 10 rounds (*t*(98) = 7.11, *p* < .001, *d* = 1.42, *BF* > 100). Crucially, the two groups did not differ during the 10 middle rounds (*t*(98) = 1.58, *p* = .12, *d* = 0.32, *BF* = 0.6). These results show that the hybrid model is an acceptable generative model of human behavior, and can replicate the behavioral effects found in Experiment 4. Therefore, the hybrid model not only captures the occurrence of a learning-to-learn effect, but also *when* this effect is likely to appear.

## General discussion

The studies reported in this paper demonstrate that participants can discover latent functional structure in a structured multi-armed bandit task, that they use this knowledge to generalize and thereby improve their performance, and that they learn to learn across rounds, progressively improving at identifying the latent structure (Experiments 1). Our studies also rule out some heuristics, like only sampling the extreme options (Experiment 2) or satisficing (Experiment 3). We also find a clear order effect: participants especially benefit from exposure to latent structure early on (Experiment 4). Additionally, participants can also detect and use structure in tasks where the underlying function was nonlinear (Experiment 5).

Our model comparison assessed a pool of 10 models in total, revealing that three ingredients are necessary to describe human behavior in our task. First, participants clearly generalize across arms in our task. We captured this ability using a Gaussian Process regression model that learns to generalize across arms within a round. The second ingredient is generalization across rounds. We captured this ability by a model that clustered the reward distribution across rounds, giving rise to an inductive bias for successive rounds. The final ingredient is uncertainty-guided exploration: participants tend to sample options with a higher predictive uncertainty. We captured this exploratory behavior by incorporating an uncertainty bonus into the policy. This strategy is equivalent to upper confidence bound sampling, a heuristic solution to the exploration-exploitation dilemma, which inflates current expectations of rewards by their uncertainty (Auer et al., 2002; Srinivas et al., 2012), and detectable in subject behavior by significantly positive estimates of the GP-RBF’s uncertainty weight in the choice function. Putting these three components together leads to a powerful model of human adaptive behavior in our task, which can generate human-like behavior and is recoverable.

It might be surprising that a hybrid model (linearly combining the predictions of the Clustering and the GP-RBF model) predicted participants’ choices best. For example, one might have expected that the Bayesian structure learning Gaussian Process model, which estimated the posterior of each structure over rounds, would win the model comparison, since it more closely maps onto the generative process underlying our experiment. However, this model already starts out with an explicit assumption about the existence of latent functional structures, whereas participants in our task had to discover this themselves. Indeed, our results of Experiment 4 showed that participants do not pick up on the latent structure when the first rounds of the experiment contained no structure.

### Related work

Much of the previous work on bandit tasks has relied on reinforcement learning models (e.g., Busemeyer & Stout, 2002) or learning from past examples (e.g., Plonsky, Teodorescu, & Erev, 2015). Here we argued that people learn something deeper about the task—its structure, represented by functional relations.

We are not the first to investigate how generalization affects behavior in multi-armed bandit tasks. Gershman and Niv (2015) studied a task in which the rewards for multiple options were drawn from a common distribution, which changed across contexts. Some contexts were “rich” (options tended to be rewarding) and some contexts were “poor” (options tended to be non-rewarding). Participants learned to sample new options more often in rich environments than in poor environments. We believe that our model could also capture these generalization effects by learning about the features of each context.

Wu, Schulz, Speekenbrink, et al. (2018) told people explicitly that the rewards were a smooth function of the options’ positions in a spatial layout; behavior on this task was well-captured by a Gaussian Process model of function learning and exploration. Schulz, Wu, Huys, Krause, and Speekenbrink (2018) used the same paradigm to study scenarios in which participants had to cautiously optimize functions while never sampling below a given threshold. Our results go beyond these studies as we show that participants learn about functional structure incidentally, i.e., without knowing that there exists a latent structure from the start, and show signatures of learning-to-learn. Moreover, we have presented a hybrid model of clustering and generalization that captured participants’ behavior across a whole range of structured multi-armed bandit tasks.

Schulz, Konstantinidis, and Speekenbrink (2017) investigated how contextual information (observable signals associated with each option) can aid generalization and exploration in tasks where the context is linked to an option’s quality by an underlying function. We find that functional generalization even matters if no explicit features are presented.

A number of tasks have used bandits that lack spatial or perceptual cues, but nonetheless have an underlying correlation structure. Wimmer, Daw, and Shohamy (2012) had participants perform a 4-armed bandit task in which pairs of options were perfectly correlated (i.e., the true number of “latent” options was 2). Participants then used this correlation structure to infer that when one option of each pair changed its value, the other option should change correspondingly. Reverdy, Srivastava, and Leonard (2014) showed how a correlation structure between arms can enhance decision making performance over short time horizons. Stojic, Analytis, and Speekenbrink (2015) found that participants can use features to guide their exploration in a contextual multi-armed bandit. Stojic, Schulz, Analytis, and Speekenbrink (2018) used a similar paradigm to test how participants explore novel options.

Multiple studies have shown that people can learn to leverage latent structure such as hierarchical rules (Badre et al., 2010), similarity across contexts (Collins, 2017; Collins & Frank, 2013) or between arms (Acuna & Schrater, 2010), even when not cued to do so. This suggests that structure learning may be incidental, and perhaps even obligatory. To the best of our knowledge, we are first to show how functional generalization and clustering interact to create complex behavior in human reinforcement learning tasks.

Gureckis and Love (2009) observed that participants can use the covariance between changing payoffs and the systematic movement of a state cue to generalize experiences from known to novel states. To explain this phenomenon, they postulated a model that uses a linear network to extrapolate the experienced payoff curves. This model can estimate action values for different choices by extrapolating rewards to unexperienced states, and matched well with participants’ behavior in several experiments (Gureckis & Love, 2009; Otto, Gureckis, Markman, & Love, 2009). Although this model is similar to a function learning approach of generalization in bandit tasks with explicit state cues, here we have focused on situations without explicit cues and payoff changes.

Finally, several studies have investigated how participants learn about functions which relate features to outcomes, both when the outcome is a continuous variable (i.e., function learning – see, for example, Busemeyer, Byun, Delosh, & McDaniel, 1997; Carroll, 1963; Hammond, 1955; Kalish, Lewandowsky, & Kruschke, 2004) and when it is a discrete variable (i.e., category learning – see, for example, J. K. Kruschke, 1992; Medin & Schaffer, 1978; Nosofsky, 1984). A related paradigm stemming from this tradition is multiple cue probability learning (J. K. Kruschke & Johansen, 1999), in which participants are shown many cues that are probabilistically related to an outcome and have to learn the underlying function between the cues and expected outcomes. However, there is no need for exploration in these tasks and participants are explicitly told to learn about the underlying function.

### Outstanding questions

In this work we have mostly focused on an underlying linear structure (but see Exp. 5), which is known to be relatively easy for participants to learn (DeLosh, Busemeyer, & McDaniel, 1997). However, people are able to learn about non-linear functions as well (Kalish, 2013; Schulz, Tenenbaum, et al., 2017). Thus, an obvious next step is to assess how well people can detect and use other types of latent structure such as non-linear functions. Currently, our results suggest that participants benefit from an underlying non-linear structure, at least in our change point experiment (Exp. 5). However, the maximum of the change point function changed over rounds, and therefore learning about the underlying structure was particularly difficult. We also did not ask participants to generalize to completely novel scenarios (Gershman & Niv, 2015) or to predict the reward for new arms (Wu, Schulz, Garvert, Meder, & Schuck, 2018), an important test of our models that should be assessed in future studies.

Furthermore, the reward in all of our experiments was continuous, which might have made learning about the underlying structure easier. Future studies could therefore investigate the effects of having categorical rewards such as binary feedback of wins and losses. We expect that binary rewards would provide a less informative reward signal and thus that both participants and our models would perform substantially worse.

Finally, we have modeled function learning and clustering as separate processes that are linearly combined. While this was the best fitting model in statistical terms and represents dissociable computational strategies, it is an open question whether these strategies are represented separately in the human brain. One possibility is that the spatial representation required for function learning in our task might be supported by the hippocampus, medial temporal cortex and ventral prefrontal cortex, regions of the brain hypothesized to be important for spatial and conceptual task representations (Constantinescu, O’Reilly, & Behrens, 2016; Stachenfeld, Botvinick, & Gershman, 2017; Wilson, Takahashi, Schoenbaum, & Niv, 2014). Theoretical modeling has proposed that fronto-cortical and striatal interactions support clustering-based generalization (Collins & Frank, 2013), and empirical work has linked clustering-based generalization with fronto-central EEG signal (Collins & Frank, 2016). Whether these two processes are linked by a shared neural representation is unknown and a topic for future research.

### Conclusion

We studied how people search for rewards in a paradigm called *structured multi-armed bandits*. In this paradigm, the expected reward of an option is a function of its spatial position. Behaviorally, we found that participants were able to detect and use the latent functional structure, with rapid learning across trials and learning-to-learn improvements over rounds. This behavior was best captured by a hybrid model combining function learning within rounds and clustering options across rounds. These results shed new light on how humans explore complex environments using a combination of three powerful mechanisms: functional generalization, contextual clustering, and uncertainty-based exploration. Our results also show that human behavior, even in tasks that look quite similar to traditional bandit tasks, can be a lot richer than what traditional accounts of human reinforcement learning postulated. In particular, combining two mechanisms that have previously been investigated independently, functional generalization and clustering, with an upper confidence bound sampling algorithm, we arrive at a model that is strikingly similar to human behavior. This model can capture participants’ ability to learn extremely fast if a bandit is governed by a latent functional structure and also their capacity to learn-to-learn, clustering previous scenarios and thereby speeding up exploration. We hope that this model can bring us closer to capturing the humanly ability to search for rewards in complex environments and to navigate the exploration-exploitation dilemma as gracefully as they do. We also believe that it captures a fundamental truth about behavior in the real world, where the space of options is frequently vast and structured.

## Acknowledgements

We are grateful to Charley Wu for feedback on an earlier draft. This research was supported by the the Office of Naval Research under grant N000141712984. ES is supported by the Harvard Data Science Initiative.

## Appendix A Materials

### Statistical analyses and model comparison

The code for all statistical analyses and full model comparison is available at:

- https://github.com/nicktfranklin/StructuredBandits

### Experiments

All structured multi-armed bandit tasks can be played under the following links:

- Experiment 1: https://ericschulz.github.io/explin/index1.html
- Experiment 2: https://ericschulz.github.io/explin/index2.html
- Experiment 3: https://ericschulz.github.io/explin/index3.html
- Experiment 4: https://ericschulz.github.io/explin/index4.html
- Experiment 5: https://ericschulz.github.io/explin/index5.html

### Comprehension check questions

Figure A1 shows the comprehension check questions used in all of our experiments.

**Figure A1.**
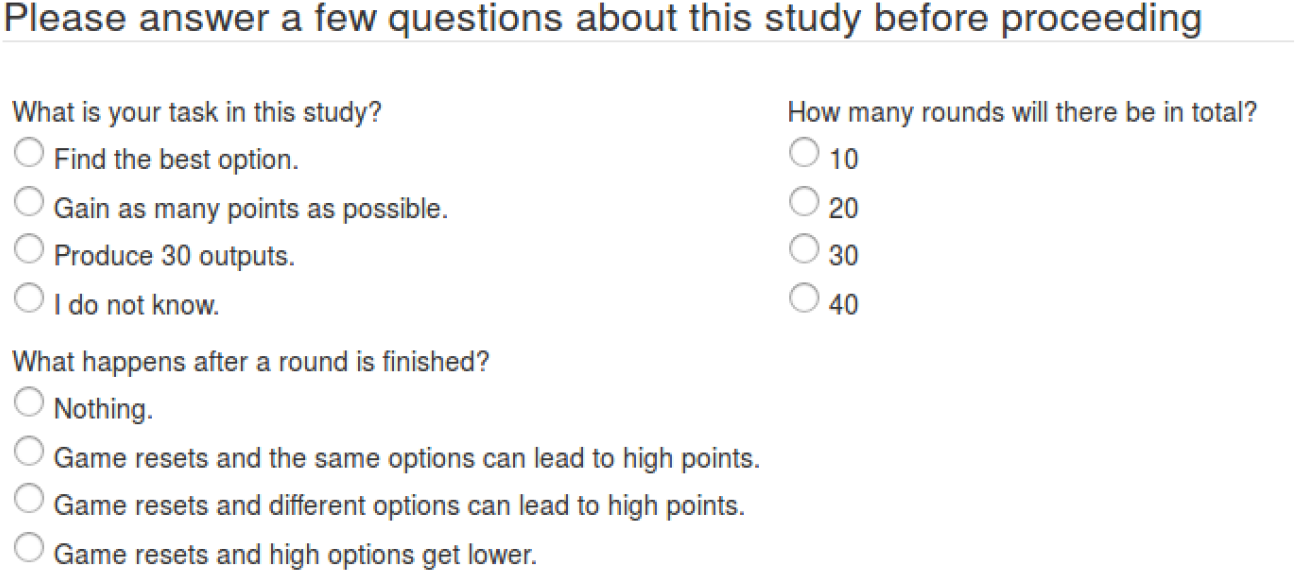
Comprehension check questions used in our experiments. Participants had to answer all of the questions correctly to proceed with the experiment. If they answered some of the questions incorrectly, they were directed back to the instructions of the experiment.

## Appendix B Statistical analysis

### Frequentist tests and Bayes factors

We report all tests assessing mean group difference by using both frequentists and Bayesian statistics. All frequentist tests are accompanied by their effect sizes, i.e., Cohen’s d (Cohen’s d; J. Cohen, 1988). The Bayesian statistics are expressed as Bayes factors (BFs). The Bayes factor quantifies the likelihood of the data under the alternative hypothesis *H_A_* relative to the likelihood of the data under the null hypothesis *H*_0_. As an example, a *BF* of 10 indicates that the data are 10 times more likely under *H_A_* than under *H*_0_, while a *BF* of 0.1 indicates that the data are 10 times more likely under *H*_0_ than under *H_A_*. We use the “default” Bayesian t-test for independent samples as proposed by Rouder and Morey (2012) for comparing group differences, using a Jeffreys-Zellner-Siow prior with its scale set to 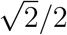. The Bayes factor for the correlational analyses is based on Jeffrey’s test for linear correlation (Ly, Verhagen, & Wagenmakers, 2016). Non-informative priors are assumed for the means and variances of the two populations, which we implement by a shifted and scaled beta-prior.

### Behavioral analysis

Markdown files for all behavioral analyses can be found under the following links:

- Experiment 1: https://ericschulz.github.io/explin/exp1.html
- Experiment 2: https://ericschulz.github.io/explin/exp2.html
- Experiment 3: https://ericschulz.github.io/explin/exp3.html
- Experiment 4: https://ericschulz.github.io/explin/exp4.html
- Experiment 5: https://ericschulz.github.io/explin/exp5.html

## Appendix C Results of pilot experiment

We report the results of a pilot experiment which we ran to assess if participants can learn in our structured experiment and also to gauge the range of expected bonuses in our task. This experiment was almost exactly the same as Experiment 1, with the only difference being that participants did not experience a 2-second delay after having received the reward of a sampled option, but rather could click through the experiment in a self-paced manner. We recruited 60 participants (19 females, mean age=32.83, SD=8.85) from Amazon Mechanical Turk. Participants performed better on the linear-positive than on the random rounds (*t*(59) = 4.87, *d* = 0.63, *BF* > 100) and also better on the linear-negative than on the random rounds (*t*(59) = 7.25, *d* = 0.94, *BF* > 100). Participants also showed signatures of learning-to-learn, sampling the leftmost and rightmost options more frequently during the first five trials over round, *r* = 0.19, *p* < .001. We therefore had good reasons to believe that these effects would replicate in our main experiments. Furthermore, we used participants’ performance in the pilot experiment to calculate the expected range of bonuses, which we then used to inform participants in our main experiments. The range of achieved bonuses in the pilot experiment was between $0.65 and $1.04. We therefore told participants in the main experiments that the expected range of rewards would be between $0.65 and $1.05.

## Appendix D Heuristic Strategies

We evaluate two heuristic strategies in addition to the models described in the main text as to how well they describe participants’ behavior in Experiment 1. The first strategy (labeled “LeftToRight”) tries every option from left to right and then chooses the one with maximum reward, sampling it until the end of the round. The second strategy (labeled “LeftRightCenter”) tries the extremes first and then samples two options from the middle and then chooses and samples the best option until the end of a round. These strategies were instantiated as vectors *m^h^* and were used in combination with sticky-choice to create choice values:

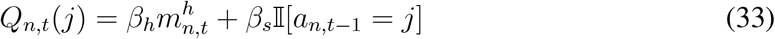

These choice values are used in the decision function (Equation 26), and hierarchical model fitting is performed as before. Figure D1 shows the results of the model fitting for all models evaluate in Experiment 1, including the two heuristic strategies. Although the two heuristic models perform better than the Sticky-Choice rule alone, they perform substantially worse than the winning hybrid model.

**Figure D1.**
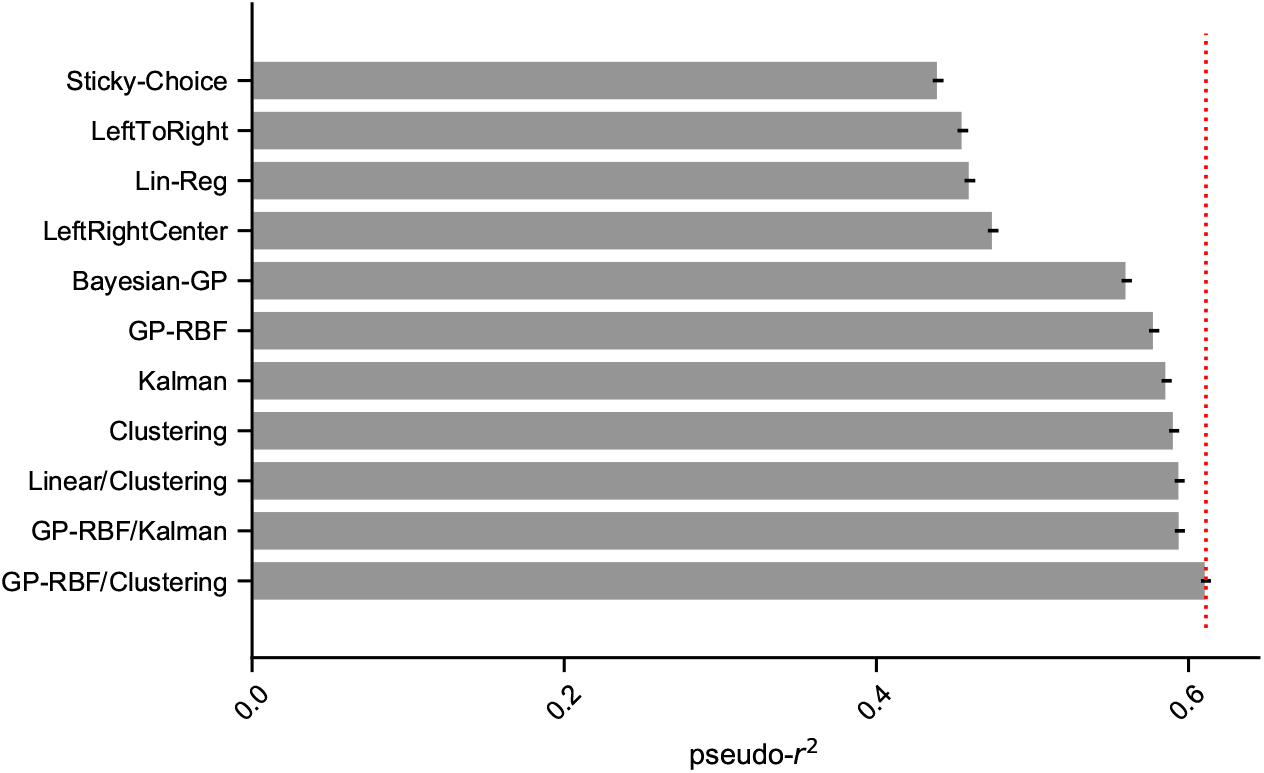
Pseudo-*r*^2^ for each model evaluated in Experiment 1.

## Appendix E Ruling out the GP-RBF + Kalman hybrid model

The hybrid combination of the GP-RBF and the Kalman filter model also performed reasonably well, particularly in the scrambled Experiment 2, where it explained participants’ behavior as good as the GP-RBF/Clustering model (but see Experiment 3 for contrasting effects). Here, we further rule out this model using generative model simulations and model recovery analysis. We simulate 100 participants using the GP-RBF/Kalman model to assess its performance in Experiment 1. Even though the GP-RBF/Kalman model produces higher rewards in the linear-positive (*t*(99) = 14.05, *p* < .001, *d* = 1.41, *BF* > 100) and linear-negative (*t*(99) = 14.27, *p* < .001, *d* = 1.43, *BF* > 100) rounds than in the random rounds, it cannot reproduce the empirically observed learning-to-learn effect, showing no correlation between rounds and rewards (*ρ* = 0.003, *p* = .86) as well as rounds and the advantage of structured rounds (*r* = 0.09, *p* = .61). Finally, we assess how well either the GP-RBF/Kalman or the GP-RBF/Clustering model are able to model the data generated by the GP-RBF/Kalman model. The GP-RBF/Kalman model achieves a predictive accuracy of *r*^2^ = 0.58 and a LOO of 22861.2, whereas the GP-RBF/Clustering model achieves a lower accuracy of *r*^2^ = 0.38 and a loo of 33590. Thus, the GP-RBF/Kalman model could have been identified as the correct model in our model comparison procedure.

## Appendix F Model comparison results

**Table F1.** Model comparison results for Experiment 1 (Linear functions). LOO, differential LOO and the standard error of LOO shown for each model. Mean (group level) parameter estimates (β) are shown for each model. For hybrid models, β_μ_1__ and β_μ_2__ refer to the means of the first and second model respectively. Asterisks (*) denote parameter estimates that do not include 0 in the 95% posterior high density interval.

| Model | LOO | dLOO | St. Error | $\beta_{\mu_1}$ | $\beta_{\sigma_1}$ | $\beta_{\mu_2}$ | $\beta_{\sigma_2}$ | $\beta_{\text{sticky}}$ |
| --- | --- | --- | --- | --- | --- | --- | --- | --- |
| *Chance* | 132253 | — | — | — | — | — | — | — |
| Sticky-choice | 74117 | 22693 | 458 | — | — | — | — | 2.82* |
| Linear regression | 71422 | 19998 | 452 | 0.19* | -0.29* | — | — | 2.65* |
| Bayesian GP | 58138 | 6714 | 438 | 0.17* | -0.01 | — | — | 1.66* |
| GP-RBF | 55815 | 4391 | 434 | 0.17* | -0.00 | — | — | 1.68* |
| Kalman filter | 54757 | 3334 | 429 | 0.19* | 0.08 | — | — | 1.49* |
| Clustering model | 54121 | 2698 | 426 | 0.28* | -0.15 | — | — | 1.20* |
| Linear Reg + Clustering | 53652 | 2228 | 423 | 0.07* | -0.01 | 0.29* | -0.13* | 1.17* |
| GP-RBF + Kalman | 53632 | 2208 | 426 | 0.08* | 0.02* | 0.13* | 0.06* | 1.50* |
| **GP-RBF + Clustering** | **51423** | **0** | **245** | **0.10\*** | **0.03\*** | **0.20\*** | **0.01** | **1.16\*** |

**Table F2.** Model comparison results for Experiment 2 (Scrambled functions). LOO, differential LOO and the standard error of LOO shown for each model. Mean (group level) parameter estimates (β) are shown for each model. For hybrid models, β_μ_1__ and β_μ_2__ refer to the means of the first and second model respectively. Asterisks (*) denote parameter estimates that do not include 0 in the 95% posterior high density interval.

| Model | LOO | dLOO | St. Error | $\beta_{\mu_1}$ | $\beta_{\sigma_1}$ | $\beta_{\mu_2}$ | $\beta_{\sigma_2}$ | $\beta_{\text{sticky}}$ |
| --- | --- | --- | --- | --- | --- | --- | --- | --- |
| *Chance* | 102309 | — | — | — | — | — | — | — |
| Sticky-choice | 61624 | 17826 | 392 | — | — | — | — | 2.62* |
| Linear regression | 59844 | 16046 | 390 | 0.13* | −0.32* | — | — | 2.49* |
| GP-RBF | 47516 | 3718 | 385 | 0.16* | −0.01 | — | — | 1.66* |
| Bayesian GP | 47439 | 3641 | 394 | 0.17* | −0.00 | — | — | 1.61* |
| Clustering model | 45596 | 1798 | 379 | 0.27* | −0.18* | — | — | 1.32* |
| Linear Reg + Clustering | 45429 | 1450 | 378 | 0.04* | −0.08 | 0.26* | −0.18* | 1.29* |
| Kalman filter | 44355 | 557 | 393 | 0.19* | 0.02 | — | — | 1.35* |
| **GP-RBF + Clustering** | **43858** | **60** | **378** | **0.09\*** | **0.02\*** | **0.19\*** | **0.06** | **1.33\*** |
| **GP-RBF + Kalman** | **43798** | **0** | **378** | **−0.01** | **0.06\*** | **0.23\*** | **−0.05** | **1.33\*** |

**Table F3.** Model comparison results for Experiment 3 (Shifted functions). LOO, differential LOO and the standard error of LOO shown for each model. Mean (group level) parameter estimates (β) are shown for each model. For hybrid models, β_μ_1__ and β_μ_2__ refer to the means of the first and second model respectively. Asterisks (*) denote parameter estimates that do not include 0 in the 95% posterior high density interval.

| Model | LOO | dLOO | St. Error | $\beta_{\mu_1}$ | $\beta_{\sigma_1}$ | $\beta_{\mu_2}$ | $\beta_{\sigma_2}$ | $\beta_{\text{sticky}}$ |
| --- | --- | --- | --- | --- | --- | --- | --- | --- |
| *Chance* | 163444 | – | – | – | – | – | – | – |
| Sticky-choice | 120859 | 27309 | 430 | – | – | – | – | 2.15* |
| Linear regression | 115895 | 22345 | 444 | 0.24* | –0.22* | – | – | 1.97* |
| Bayesian GP | 109823 | 16272 | 458 | 0.07* | –0.13* | – | – | 1.08* |
| Kalman filter | 109614 | 16063 | 459 | 0.04* | –0.04* | – | – | 1.07* |
| Clustering model | 103177 | 9626 | 468 | 0.10* | –0.04* | – | – | 0.84* |
| GP-RBF | 101631 | 8081 | 471 | 0.11* | 0.03* | – | – | 1.61* |
| Linear Reg + Clustering | 100949 | 7399 | 468 | 0.15* | 0.06* | 0.10* | –0.03* | 0.82* |
| GP-RBF + Kalman | 98831 | 5280 | 473 | 0.09* | 0.08* | 0.02* | –0.07* | 1.20* |
| **GP-RBF + Clustering** | **93550** | **0** | **477** | **0.07\*** | **0.10\*** | **0.09\*** | **0.00** | **1.06\*** |

**Table F4.** Model comparison results for Experiment 4 (Structure early vs. late). LOO, differential LOO and the standard error of LOO shown for each model. Mean (group level) parameter estimates (β) are shown for each model. For hybrid models, β_μ_1__ and β_μ_2__ refer to the means of the first and second model respectively. Asterisks (*) denote parameter estimates that do not include 0 in the 95% posterior high density interval.

| Model | LOO | dLOO | St. Error | $\beta_{\mu_1}$ | $\beta_{\sigma_1}$ | $\beta_{\mu_2}$ | $\beta_{\sigma_2}$ | $\beta_{\text{sticky}}$ |
| --- | --- | --- | --- | --- | --- | --- | --- | --- |
| *Chance* | 140986 | – | – | – | – | – | – | – |
| Sticky-choice | 88262 | 24671 | 452 | – | – | – | – | 2.48* |
| Linear regression | 86065 | 22471 | 451 | 0.13* | –0.23* | – | – | 2.36* |
| Bayesian GP | 69448 | 5857 | 451 | 0.16* | –0.06* | – | – | 1.39* |
| GP-RBF | 68376 | 4785 | 450 | 0.15* | –0.01 | – | – | 1.48* |
| Clustering model | 66262 | 2671 | 445 | 0.26* | –0.14* | – | – | 1.13* |
| Linear Reg + Clustering | 65932 | 2341 | 443 | 0.05* | –0.07* | 0.26* | –0.13* | 1.10* |
| Kalman filter | 65121 | 1530 | 454 | 0.18* | 0.00 | – | – | 1.12* |
| GP-RBF + Kalman | 64291 | 670 | 450 | 0.05* | 0.03* | 0.16* | –0.01 | 1.21* |
| **GP-RBF + Clustering** | **63591** | **0** | **443** | **0.09\*** | **0.01\*** | **0.18\*** | **–0.03** | **1.06\*** |

**Table F5.** Model comparison results for Experiment 5 (Changepoint functions). LOO, differential LOO and the standard error of LOO shown for each model. Mean (group level) parameter estimates (β) are shown for each model. For hybrid models, β_μ_1__ and β_μ_2__ refer to the means of the first and second model respectively. Asterisks (*) denote parameter estimates that do not include 0 in the 95% posterior high density interval.

| Model | LOO | dLOO | St. Error | $\beta_{\mu_1}$ | $\beta_{\sigma_1}$ | $\beta_{\mu_2}$ | $\beta_{\sigma_2}$ | $\beta_{\text{sticky}}$ |
| --- | --- | --- | --- | --- | --- | --- | --- | --- |
| *Chance* | 163444 | — | — | — | — | — | — | — |
| Sticky-choice | 102277 | 23921 | 486 | — | — | — | — | 2.54* |
| Linear regression | 101426 | 23070 | 485 | 0.04* | −0.38* | — | — | 2.49* |
| GP-RBF | 87189 | 8834 | 473 | 0.12* | −0.01 | — | — | 1.78* |
| Bayesian-GP | 83466 | 5110 | 463 | 0.13* | −1.08* | — | — | 0.95* |
| Kalman filter | 82541 | 4185 | 485 | 0.15* | −0.14* | — | — | 0.81* |
| Clustering model | 81779 | 3424 | 482 | 0.25* | −0.29* | — | — | 1.05* |
| Linear + Clustering | 81325 | 2970 | 480 | 0.03* | −0.20* | 0.25* | −0.29* | 1.05* |
| GP-RBF + Kalman | 79683 | 1327 | 482 | 0.05* | 0.07* | 0.15* | −0.18* | 0.78* |
| **GP-RBF + Clustering** | **78356** | **0** | **476** | **0.09\*** | **0.05\*** | **0.21\*** | **−0.21\*** | **1.04\*** |

## Footnotes

1 We use Spearman’s *ρ* to account for possible nonlinearities in participants’ learning curves.

2 A white noise kernel estimates the level of observation noise *σ* if *j* = *j*′ and is 0 otherwise. It is equivalent to a Kalman filter model without any innovation of uncertainty across trials.

3 Re-running the model fits for a second time revealed similar parameter estimates. We therefore believe that the negative estimates of the Clustering model’s uncertainty are more likely to be the result of participants’ strategy than of a model unidentifiability.

4 See Appendix D for further comparisons with heuristic models of human behavior.

## References

Acuna, D. E., & Schrater, P. (2010). Structure Learning in Human Sequential Decision-Making. PLoS Computational Biology, 6. doi: 10.1371/journal.pcbi.1001003

Aldous, D. J. (1985). Exchangeability and related topics., 1–198. Retrieved from http://link.springer.com/10.1007/BFb0099421 doi: 10.1007/BFb0099421

Andrychowicz, M., Denil, M., Gomez, S., Hoffman, M. W., Pfau, D., Schaul, T.,… De Freitas, N. (2016). Learning-to-learn by gradient descent by gradient descent. In Advances in Neural Information Processing Systems (pp. 3981–3989).

Auer, P., Cesa-Bianchi, N., & Fischer, P. (2002). Finite-time analysis of the multiarmed bandit problem. Machine Learning, 47, 235–256.

Badre, D., Kayser, A. S., & D’Esposito, M. (2010). Frontal cortex and the discovery of abstract action rules. Neuron, 66, 315–326.

Bechara, A., Damasio, H., Tranel, D., & Damasio, A. R. (2005). The Iowa Gambling Task and the somatic marker hypothesis: Some questions and answers. Trends in Cognitive Sciences, 9, 159–162.

Börgers, T., & Sarin, R. (2000). Naive reinforcement learning with endogenous aspirations. International Economic Review, 41, 921–950.

Boyan, J. A., & Moore, A. W. (1995). Generalization in reinforcement learning: Safely approximating the value function. In Advances in neural information processing systems (pp. 369–376).

Busemeyer, J. R., Byun, E., Delosh, E. L., & McDaniel, M. A. (1997). Learning functional relations based on experience with input-output pairs by humans and artificial neural networks. In K. Lamberts & D. R. Shanks (Eds.), Knowledge, concepts and categories. studies in cognition. (pp. 408–437). Cambridge, MA, US: MIT Press.

Busemeyer, J. R., & Stout, J. C. (2002). A contribution of cognitive decision models to clinical assessment: decomposing performance on the bechara gambling task. Psychological assessment, 14, 253.

Carroll, J. D. (1963). Functional learning: The learning of continuous functional mappings relating stimulus and response continua. ETS Research Bulletin Series, 1963, i–144.

Chapelle, O., & Li, L. (2011). An empirical evaluation of thompson sampling. In Advances in Neural Information Processing Systems (pp. 2249–2257).

Cohen, J. (1988). Statistical power analysis for the behavioral sciences. 2nd. Hillsdale, NJ: erlbaum.

Cohen, J. D., McClure, S. M., & Angela, J. Y. (2007). Should I stay or should I go? How the human brain manages the trade-off between exploitation and exploration. Philosophical Transactions of the Royal Society B: Biological Sciences, 362, 933–942.

Collins, A. G. (2017). The cost of structure leanring. Journal of Cognitive Neuroscience, 29, 1646–1655. doi: 10.1162/jocn

Collins, A. G., & Frank, M. J. (2013). Cognitive control over learning: Creating, clustering, and generalizing task-set structure. Psychological Review, 120, 190.

Collins, A. G., & Frank, M. J. (2016). Neural signature of hierarchically structured expectations predicts clustering and transfer of rule sets in reinforcement learning. Cognition, 152, 160–169.

Constantinescu, A. O., O’Reilly, J. X., & Behrens, T. E. (2016). Organizing conceptual knowledge in humans with a gridlike code. Science, 352, 1464–1468.

Daw, N. D., O’Doherty, J. P., Dayan, P., Seymour, B., & Dolan, R. J. (2006). Cortical substrates for exploratory decisions in humans. Nature, 441, 876–879. doi: 10.1038/nature04766

DeLosh, E. L., Busemeyer, J. R., & McDaniel, M. A. (1997). Extrapolation: The sine qua non for abstraction in function learning. Journal of Experimental Psychology: Learning, Memory, and Cognition, 23, 968.

Duvenaud, D. (2014). Automatic model construction with Gaussian processes (Unpublished doctoral dissertation). University of Cambridge.

Franklin, N. T., & Frank, M. J. (2018). Compositional clustering in task structure learning. PLoS Vomputational Biology, 14, e1006116.

Franklin, N. T., & Frank, M. J. (2019). Generalizing to generalize: humans flexibly switch between compositional and conjunctive structures during reinforcement learning. bioRxiv. doi: 10.1101/547406

Gershman, S. J. (2015). A unifying probabilistic view of associative learning. PLoS Computational Biology, 11, e1004567.

Gershman, S. J. (2018). Deconstructing the human algorithms for exploration. Cognition, 173, 34–42.

Gershman, S. J., & Blei, D. M. (2012). A tutorial on Bayesian nonparametric models. Journal of Mathematical Psychology, 56, 1–12.

Gershman, S. J., Blei, D. M., & Niv, Y. (2010). Context, learning, and extinction. Psychological Review, 117, 197.

Gershman, S. J., Malmaud, J., & Tenenbaum, J. B. (2017). Structured representations of utility in combinatorial domains. Decision, 4, 67–86.

Gershman, S. J., & Niv, Y. (2010). Learning latent structure: Carving nature at its joints. Current Opinion in Neurobiology, 20, 251–256.

Gershman, S. J., & Niv, Y. (2015). Novelty and inductive generalization in human reinforcement learning. Topics in Cognitive Science, 7, 391–415.

Gittins, J. C. (1979). Bandit processes and dynamic allocation indices. Journal of the Royal Statistical Society. Series B (Methodological), 148–177.

Goldstone, R. L., & Ashpole, B. C. (2004). Human foraging behavior in a virtual environment. Psychonomic bulletin & review, 11, 508–514.

Griffiths, T. L., Lucas, C., Williams, J., & Kalish, M. L. (2009). Modeling human function learning with gaussian processes. In Advances in neural information processing systems (pp. 553–560).

Gureckis, T. M., & Love, B. C. (2009). Short-term gains, long-term pains: How cues about state aid learning in dynamic environments. Cognition, 113, 293–313.

Hammond, K. R. (1955). Probabilistic functioning and the clinical method. Psychological Review, 62, 255–262.

Harlow, H. F. (1949). The formation of learning sets. Psychological Review, 56, 51.

Hastie, T. J. (2017). Generalized additive models. In Statistical models in S (pp. 249–307). Routledge.

Hotaling, J., Navarro, D., & Newell, B. (2018). Skilled bandits: Learning to choose in a reactive world.

Kalish, M. L. (2013). Learning and extrapolating a periodic function. Memory & Cognition, 41, 886–896.

Kalish, M. L., Lewandowsky, S., & Kruschke, J. K. (2004). Population of linear experts: knowledge partitioning and function learning. Psychological Review, 111, 1072.

Kruschke, J. (2014). Doing bayesian data analysis: A tutorial with r, jags, and stan. Academic Press.

Kruschke, J. K. (1992). ALCOVE: An exemplar-based connectionist model of category learning. Psychological Review, 99, 22–44. doi: 10.1037/0033-295X.99.1.22

Kruschke, J. K., & Johansen, M. K. (1999). A model of probabilistic category learning. Journal of Experimental Psychology: Learning, Memory, and Cognition, 25, 1083.

Lloyd, K., & Leslie, D. S. (2013). Context-dependent decision-making: A simple Bayesian model. Journal of The Royal Society Interface, 10, 20130069.

Lucas, C. G., Griffiths, T. L., Williams, J. J., & Kalish, M. L. (2015). A rational model of function learning. Psychonomic bulletin & review, 22, 1193–1215.

Ly, A., Verhagen, J., & Wagenmakers, E.-J. (2016). Harold Jeffreys’s default Bayes factor hypothesis tests: Explanation, extension, and application in psychology. Journal of Mathematical Psychology, 72, 19–32.

McFadden, D., et al. (1973). Conditional logit analysis of qualitative choice behavior.

Medin, D. L., & Schaffer, M. M. (1978). Context theory of classification learning. Psychological Review, 85, 207–238.

Mehlhorn, K., Newell, B. R., Todd, P. M., Lee, M. D., Morgan, K., Braithwaite, V. A.,… Gonzalez, C. (2015). Unpacking the exploration–exploitation tradeoff: A synthesis of human and animal literatures. Decision, 2, 191.

Navarro, D. J., Newell, B. R., & Schulze, C. (2016). Learning and choosing in an uncertain world: An investigation of the explore–exploit dilemma in static and dynamic environments. Cognitive Psychology, 85, 43–77.

Navarro, D. J., Tran, P., & Baz, N. (2018). Aversion to option loss in a restless bandit task. Computational Brain & Behavior, 1, 151–164.

Nosofsky, R. M. (1984). Choice, similarity, and the context theory of classification. Journal of Experimental Psychology: Learning, Memory, and Cognition, 10, 104–114. doi: 10.1037/0278-7393.10.1.104

Otto, A. R., Gureckis, T. M., Markman, A. B., & Love, B. C. (2009). Navigating through abstract decision spaces: Evaluating the role of state generalization in a dynamic decision-making task. Psychonomic Bulletin & Review, 16, 957–963.

Palminteri, S., Wyart, V., & Koechlin, E. (2017). The importance of falsification in computational cognitive modeling. Trends in Cognitive Sciences, 21, 425–433.

Piray, P., Dezfouli, A., Heskes, T., Frank, M. J., & Daw, N. D. (2018). Hierarchical bayesian inference for concurrent model fitting and comparison for group studies. bioRxiv. doi: 10.1101/393561

Plonsky, O., Teodorescu, K., & Erev, I. (2015). Reliance on small samples, the wavy recency effect, and similarity-based learning. Psychological review, 122, 621.

Rasmussen, C. E., & Williams, C. K. I. (2006). Gaussian Processes for Machine Learning. MIT Press.

Reverdy, P. B., Srivastava, V., & Leonard, N. E. (2014). Modeling human decision making in generalized Gaussian multiarmed bandits. Proceedings of the IEEE, 102, 544–571.

Rouder, J. N., & Morey, R. D. (2012). Default Bayes Factors for Model Selection in Regression., 47, 877–903. doi: 10.1080/00273171.2012.734737

Saeedi, A., Kulkarni, T. D., Mansinghka, V. K., & Gershman, S. J. (2017). Variational particle approximations. The Journal of Machine Learning Research, 18, 2328–2356.

Salvatier, J., Wiecki, T. V., & Fonnesbeck, C. (2016). Probabilistic programming in Python using PyMC3. PeerJ Computer Science, 2, e55.

Sanborn, A., Griffiths, T., & Navarro, D. (2006). A more rational model of categorization.

Schulz, E., & Gershman, S. J. (2019). The algorithmic architecture of exploration in the human brain. Current opinion in neurobiology, 55, 7–14.

Schulz, E., Konstantinidis, E., & Speekenbrink, M. (2017). Putting bandits into context: How function learning supports decision making. Journal of Experimental Psychology: Learning, Memory, and Cognition.

Schulz, E., Speekenbrink, M., & Krause, A. (2018). A tutorial on Gaussian process regression: Modelling, exploring, and exploiting functions. Journal of Mathematical Psychology, 85, 1–16.

Schulz, E., Tenenbaum, J. B., Duvenaud, D., Speekenbrink, M., & Gershman, S. J. (2017). Compositional inductive biases in function learning. Cognitive Psychology, 99, 44–79.

Schulz, E., Tenenbaum, J. B., Reshef, D. N., Speekenbrink, M., & Gershman, S. (2015). Assessing the perceived predictability of functions. In Cogsci.

Schulz, E., Wu, C. M., Huys, Q. J. M., Krause, A., & Speekenbrink, M. (2018). Generalization and search in risky environments. Cognitive Science, 42, 2592–2620. doi: 10.1111/cogs.12695

Shepard, R. N. (1987). Toward a universal law of generalization for psychological science. Science, 237, 1317–1323.

Speekenbrink, M., & Konstantinidis, E. (2015). Uncertainty and exploration in a restless bandit problem. Topics in Cognitive Science, 7, 351–367.

Srinivas, N., Krause, A., Kakade, S., & Seeger, M. (2012, May). Information-Theoretic Regret Bounds for Gaussian Process Optimization in the Bandit Setting. IEEE Transactions on Information Theory, 58, 3250–3265. doi: 10.1109/TIT.2011.2182033

Stachenfeld, K. L., Botvinick, M. M., & Gershman, S. J. (2017). The hippocampus as a predictive map. Nature neuroscience, 20, 1643.

Steingroever, H., Wetzels, R., Horstmann, A., Neumann, J., & Wagenmakers, E.-J. (2013). Performance of healthy participants on the iowa gambling task. Psychological assessment, 25, 180.

Stojic, H., Analytis, P. P., & Speekenbrink, M. (2015). Human behavior in contextual multi-armed bandit problems. In Proceedings of the Thirty-Seventh Annual Conference of the Cognitive Science Society (pp. 2290–2295).

Stojic, H., Schulz, E., Analytis, P. P., & Speekenbrink, M. (2018). It’s new, but is it good? how generalization and uncertainty guide the exploration of novel options. PsyArXiv.

Tenenbaum, J. B., & Griffiths, T. L. (2001). Generalization, similarity, and bayesian inference. Behavioral and Brain Sciences, 24, 629–640.

Vehtari, A., Gelman, A., & Gabry, J. (2017). Practical Bayesian model evaluation using leave-one-out cross-validation and WAIC. Statistics and Computing, 27, 1413–1432.

Whittle, P. (1980). Multi-Armed Bandits and the Gittins Index. Journal of the Royal Statistical Society. Series B (Methodological), 42, 143–149.

Wiecki, T. V., Sofer, I., & Frank, M. J. (2013). Hddm: hierarchical bayesian estimation of the drift-diffusion model in python. Frontiers in Neuroinformatics, 7, 14.

Wilson, R. C., Takahashi, Y. K., Schoenbaum, G., & Niv, Y. (2014). Orbitofrontal cortex as a cognitive map of task space. Neuron, 81, 267–279.

Wimmer, G. E., Daw, N. D., & Shohamy, D. (2012). Generalization of value in reinforcement learning by humans. European Journal of Neuroscience, 35, 1092–1104.

Wu, C. M., Schulz, E., Garvert, M. M., Meder, B., & Schuck, N. W. (2018). Connecting conceptual and spatial search via a model of generalization. bioRxiv, 258665.

Wu, C. M., Schulz, E., Speekenbrink, M., Nelson, J. D., & Meder, B. (2017). Mapping the unknown: The spatially correlated multi-armed bandit. bioRxiv. doi: 10.1101/106286

Wu, C. M., Schulz, E., Speekenbrink, M., Nelson, J. D., & Meder, B. (2018). Generalization guides human exploration in vast decision spaces. Nature Human Behaviour, 2, 915.

Zhang, S., & Yu, A. J. (2013). Forgetful Bayes and myopic planning: Human learning and decision-making in a bandit setting. In Advances in Neural Information Processing Systems (pp. 2607–2615).

